# Intelligence-associated Polygenic Scores Predict g, Independent of Ancestry, Parental Educational Levels, and Color among Hispanics in comparison to European, European-African, and African Americans

**DOI:** 10.1101/2020.09.24.312074

**Authors:** Bryan J. Pesta, John G. R. Fuerst, Davide Piffer, Emil O. W. Kirkegaard

## Abstract

Polygenic scores for educational attainment and intelligence (eduPGS), genetic ancestry, and cognitive ability have been found to be inter-correlated in some admixed American populations. We argue that this could either be due to causally-relevant genetic differences between ancestral groups or be due to population stratification-related confounding. Moreover, we argue that it is important to determine which scenario is the case so to better assess the validity of eduPGS. We investigate the confounding vs. causal concern by examining, in detail, the relation between eduPGS, ancestry, and general cognitive ability in East Coast Hispanic and non-Hispanic samples. European ancestry was correlated with *g* in the admixed Hispanic (*r* = .30, *N* = 506), European-African (*r* = .26, *N* = 228), and African (*r* = .084, *N* = 2,179) American samples. Among Hispanics and the combined sample, these associations were robust to controls for racial / ethnic self-identification, genetically predicted color, and parental education. Additionally, eduPGS predicted *g* among Hispanics (*B* = 0.175, *N* = 506) and all other groups (European: *B* = 0.230, *N* = 4914; European-African: *B* = 0.215, *N* = 228; African: *B* = 0.126, *N* = 2179) with controls for ancestry. Path analyses revealed that eduPGS, but not color, partially statistically explained the association between *g* and European ancestry among both Hispanics and the combined sample. Of additional note, we were unable to account for eduPGS differences between ancestral populations using common tests for ascertainment bias and confounding related to population stratification. Overall, our results suggest that eduPGS derived from European samples can be used to predict *g* in American populations. However, owing to the uncertain cause of the differences in eduPGS, it is not yet clear how the effect of ancestry should be handled. We argue that more research is needed to determine the source of the relation between eduPGS, genetic ancestry, and cognitive ability.

## 1. Introduction

General intelligence is perhaps the most powerful variable in social science, as it often strongly predicts numerous academic, economic, occupational, social, and health related outcomes (Schmidt, 2002; Gottfredson, 2003; Deary, 2010). Owing to its relation to general human well-being in contemporary society, understanding the causes of individual differences in general intelligence is of significant social importance. Moreover, the search for the causes of these differences has led many to investigate the environmental and genetic determinants of general intelligence (Plomin, Defries, Knopik, & Neiderhiser, 2014). More recently, large scale Genome Wide Association Studies (GWAS) have been conducted to identify the genetic variants underlying the hereditary component of individual differences (see, e.g., Lee et al., 2018; Sniekers et al., 2017; Savage et al., 2018). At present, among Europeans, 4-10% of the variance in cognitive ability can be explained by intelligence and educational polygenic scores (eduPGS; Plomin & von Stumm, 2018).

GWAS studies have mostly been conducted on European populations. However, among certain groups (e.g., African Americans), European-based eduPGS have been found to display attenuated predictive accuracy with respect to cognitive ability (Rabinowitz et al., 2019; Lasker, Pesta, Fuerst, & Kirkegaard, 2019; Guo, Lin, & Harris, 2019). There are several possible reasons for this attenuation. One explanation for the attenuated predictive accuracy appeals to possible lower within-group heritability in non-European groups (e.g., Rabinowitz et al., 2019). This is a theoretically plausible account, since predictive accuracy is a function of heritability (Daetwyler, Villanueva, & Woolliams, 2008). However, because the heritability of IQ is similar across racial and ethnic groups (for a meta-analytic review, see Pesta, Kirkegaard, te Nijenhuis, Lasker, & Fuerst, 2020), a more likely possibility is decay of linkage disequilibrium (LD), which results in different correlations between SNPs across different ancestry groups (Zanetti & Weale, 2018).

The significance of LD decay in attenuating transethnic predictive accuracy depends on the specific populations in question, as the impact of LD decay will be modified by assortative mating, selection, admixture, genetic drift and other factors, which vary across populations. For example, although the predictive accuracy of eduPGS was found to be attenuated among African Americans, this did not seem to be the case in an admixed, Brazilian sample (Horta, Hartwig & Victora, 2018). Nor was it the case for Asian and non-Black Hispanic Americans (Guo, Lin, & Harris, 2019). A similar pattern has been discovered for other, medically related PGS, with the accuracy of PGS being attenuated the most for African Americans (Duncan et al., 2019, Figure 2). This is likely because African Americans are primarily a sub-Saharan African group, and because sub-Saharan Africans are the continental lineage most genetically distant from Europeans (and other major races; Duncan et al., 2019). Generally, the validity of eduPGS has to be tested on a population by population basis and can not be assumed to generalize.

Moreover, among admixed populations, evaluating the validity of PGS is complicated when ancestry, PGS, and traits of interest covary. In these situations, as detailed by Lawson et al. (2020) the covariance between ancestry, PGS, and the trait could result from a combination of ascertainment bias and confounding related to population stratification, or could be a result of genetic differences related to the trait. Depending on the scenario, corrections for ancestry may either be accurate, overcorrect, or undercorrect. Thus, it is important to evaluate possible scenarios.

In the case of admixed American populations, previous research suggests that ancestry, eduPGS, and cognitive ability scores are inter-correlated (Lasker et al., 2019). This is found when examining self-identified racial/ethnic (SIRE) groups separately. For example, Kirkegaard, Woodley of Menie, Williams, Fuerst, and Meisenberg (2019), Lasker et al. (2019), Warne (2020), and Guo, Lin, and Harris (2019) found that cognitive ability was associated with admixture components within ethnic groups (e.g., self-identifying African Americans, Hispanic, Minorities, African-and Black Hispanics). These findings were robust to controls for SIRE, despite the fact that, as Fang et al. (2019, *p*. 764) noted, SIRE “acts as a surrogate to an array of social, cultural, behavioral, and environmental variables” and so “stratifying on SIRE has the potential benefits of reducing heterogeneity of these non-genetic variables and decoupling the correlation between genetic and non-genetic factors.”

There are two obvious scenarios which could explain such results. These are depicted in Figure 1. The first, (a), is the confounding scenario. Here, ancestry is associated with causally irrelevant eduPGS-related loci as a result of ascertainment bias and confounding related to population stratification (e.g., Martin et al., 2017; Kim, Patel, Teng, Berens, & Lachance, 2018); simultaneously, ancestry is also associated with cognitive ability by way of the environment. The environmental differences cause socioeconomic ones which, in turn, cause cognitive ability ones. In this scenario, eduPGS would have a spurious relation to *g* between groups, despite a causal one within groups. The second, (b), is a causal scenario. In this case, ancestry is associated with causally relevant eduPGS-related loci due to evolutionary history. These genetic differences cause cognitive ability ones which, in turn, lead to socioeconomic ones. In this case, eduPGS would be a component with constitutive explanatory relevance in the relation between ancestry and *g*.^1^

**Figure 1.**
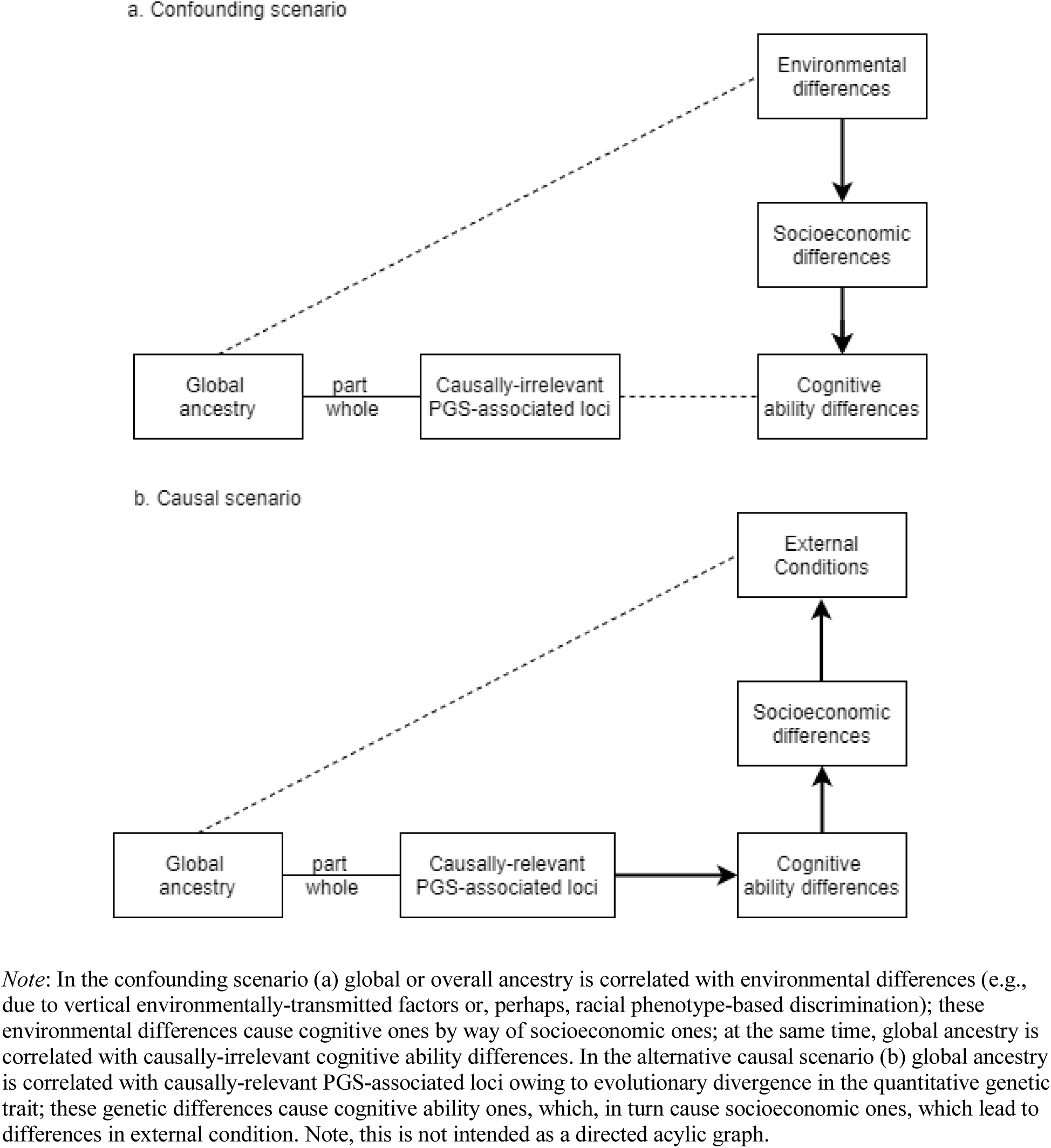
Theoretical Scenarios Depicting the Relation Between Ancestry, eduPGS, Cognitive Ability, and Socioeconomic Differences.

**Figure 2.**
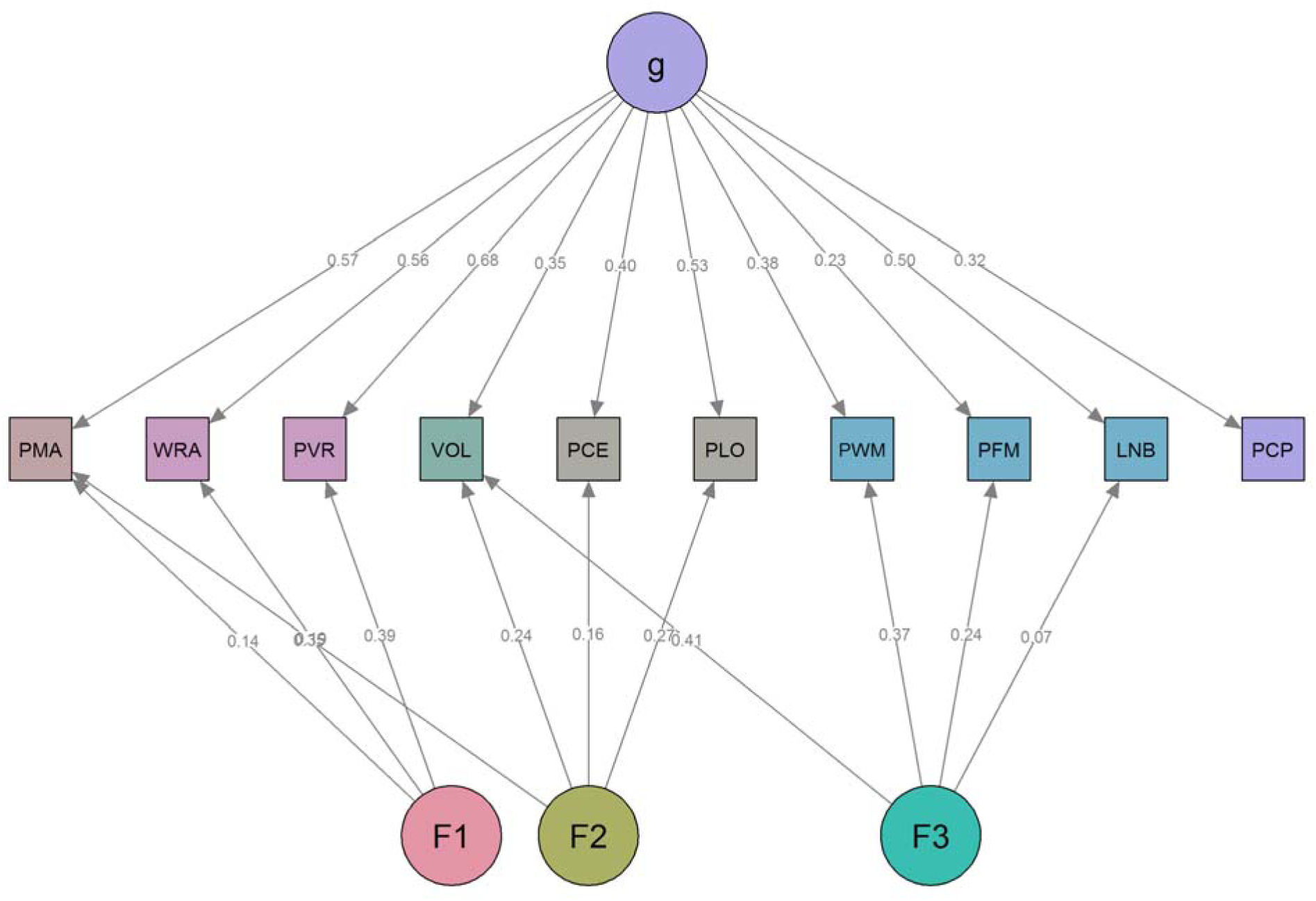
Three Factor Model for the Penn Cognitive Battery.

Determining which is the case is critical for an accurate interpretation of PGS in admixed American populations (Lawson et al., 2020). This is because if the causal scenario is correct, controlling for ancestry can attenuate the true effect of PGS. However, if the confounding scenario is correct, leaving ancestry unadjusted can spuriously inflate the effect of PGS. We do not attempt to determine which scenario is correct in this paper. Rather, we investigate if there is a robust confounding vs. causal problem with respect to cognitive ability that needs to be solved by future research. This would be the case if the data was found consistent with either a causal or confound scenario as illustrated above.

To do this, we used the Trajectories of Complex Phenotype sample, which is a large population-representative Philidelphian sample. This sample has a number of advantages over other available ones. First, it is a local sample and so the issue of ancestry-related geographic confounding (Lawson et al., 2020; Kirkegaard et al. 2019) is not of significant concern. Second, the cognitive battery is well designed and allows for a robust examination of psychometric bias between ethnic groups. Third, the heritability of cognitive ability has been previously reported for the two largest ethnic groups (African and European Americans; Mollon et al., 2018); this is important as the predictive validity of PGS depends on the trait heritability (Pesta et al., 2020).

We focus here on the relation between ancestry, eduPGS, and general intelligence in the Hispanic sample and compare these results with those from European, biracial European-African, and African American samples. As far as we are aware, no previous research has investigated this issue using a largely Caribbean Hispanic sample.

For background, Hispanics residing in Philadelphia are largely a Caribbean-origin, admixed population, with a predominantly Puerto Rican component (69% Puerto Rican; US Census Bureau, 2016). As of 2010, the majority of self-identifying Puerto Ricans (60%) reside stateside, with the largest communities in New York City, Philadelphia, and Chicago. Of all stateside Puerto Ricans, 70% are born in the continental U.S. (mean age = 29 years), with a higher stateside-born proportion in younger cohorts (Motel & Patten, 2012). Hispanics, in general, have been found to score significantly below European Americans on measures of general intelligence, with Roth et al. (2017; Table 12) calculating meta-analytic effects of *d* = -0.65 to -1.04. Puerto Ricans, both in Puerto Rico and on the mainland, score similarly below US non-Hispanic Europeans, and have been found to do so since the early 1900s (Malloy, 2014).

Given these typically large differences, we first assess measurement invariance to ensure that our measure of cognitive ability functions the same for European and Hispanic Americans. After, we report the bivariate correlations for eduPGS, ancestry, parental education, and color. Next, via regression, we examine whether associations between genetic ancestry and general cognitive ability can be accounted for by either SIRE, skin color, or parental education.

Following this, we evaluate the predictive validity of four constructions of eduPGS. We next examine whether the validity of eduPGS is robust to controls for genetic ancestry and color. Using path modeling, we also explore the extent to which eduPGS can statistically explain the association between European genetic ancestry and general cognitive ability.

After, we apply Jensen’s Method of Correlated Vectors (MCV; Jensen, 1998) to examine the relation between eduPGS and *g*-loadings. The expectation is that eduPGS effects will be largest on the most *g*-loaded subtests since genetic effects, in contrast to environmental effects, tend to be *g*-loaded (Rushton & Jensen, 2010; te Nijenhuis, Choi, van den Hoek, Valueva, & Lee, 2019). This phenomenon, of a positive correlation between the vector of *g*-loading and some other vector, is referred to as a “Jensen Effect” (Woodley of Menie, Fernandes, & Hopkins, 2015). As it is, eduPGS is differentially associated with subtest scores (de la Fuente et al., 2020; Genç et al., 2020). However, these samples do not allow for a robust evaluation of whether eduPGS exhibits a Jensen Effect owing to a limited number of subtests or small sample sizes. Thus, we evaluate the matter here. We further examine the relation between *g*-loadings, group differences, and ancestry. If these differences are primarily in *g* (i.e., Spearman’s hypothesis; Jensen, 1998), then Jensen Effects are expected. Moreover, since a causal model would predict a positive manifold of Jensen Effects for eduPGS, heritability, ancestry, and group differences, these analyses can show if the data are at least as would be expected by a causal model.

Finally, we examine the eduPGS scores for common forms of ascertainment bias and confounding related to population stratification to see if we can easily account for the effects of population structure. If we can, there may be no confounding vs. causal scenario in need of resolving. To be clear, though, it is outside the scope of this paper to investigate all possible forms of confounding. We therefore restrict consideration to some forms of bias discussed in the literature. The overall goal is to better understand how eduPGS, ancestry, SIRE, color, and cognitive ability are associated with one another and to evaluate whether there is a robust confound vs. causal concern in need of further research.

## 2. Materials and Methods

The Trajectories of Complex Phenotypes (TCP) study was conducted by the Center for Applied Genomics at the Children’s Hospital of Philadelphia, and the Brain Behavior Laboratory at the University of Pennsylvania. Participants were English-speaking Philadelphians, aged 8-21 years at the time of testing (which was done primarily between 2010 to 2013). Those with severe cognitive or medical impairments were excluded from the final sample. Access to the data was obtained through dbGaP (phs000607.v3.p2) via Bryan J. Pesta.

### 2.1 Cognitive Ability

Participants were administered the Penn Computerized Neurocognitive Battery (PCNB; Gur et al., 2010; Moore, Reise, Gur, Hakonarson, & Gur, 2015), a psychometrically robust cognitive battery that incorporates tasks linked to specific brain systems. This is a widely used 1-hour neurocognitive battery, which has been previously validated for this sample (Moore et al. 2015; Satterthwaite et al., 2016), which has an age range of 8 to 22 years. The 14 PCNB subtests were designed to measure five broad behavioral domains: Executive Control, Episodic Memory, Complex Cognition, Social Cognition, and Sensorimotor Speed. The subtests are as follows: 1. Executive Control: Penn Conditional Exclusion Test (PCET; meant to assess Mental Flexibility), Penn Continuous Performance Test (PCPT; Attention), and Letter N-Back Task (LNB; Working Memory); 2. Episodic Memory: Penn Word Memory Task (PWMT; Verbal Memory), Penn Face Memory Task (PFMT; Face Memory), and Visual Object Learning Test (VOLT; Spatial Memory); 3. Complex Cognition: Penn Verbal Reasoning Test (PVRT; Language Reasoning), Penn Matrix Reasoning Test (PMRT; Nonverbal Reasoning), and Penn Line Orientation Test (PLOT; Spatial Ability); 4. Social Cognition: Penn Emotion Identification Test (PEIT; Emotion Identification), Penn Emotion Differentiation Test (PEDT; Emotion Differentiation), and Penn Age Differentiation Test (PADT; Age Differentiation). 5. Sensorimotor Speed: Motor Praxis Test (MP; Sensorimotor Speed), and Finger Tapping (Tap; Sensorimotor Speed) (Lasker et al., 2019). Participants additionally completed the Wide Range Achievement Test (WRAT), a highly-reliable broad ability measure (Moore et al., 2015).

We excluded the approximately 1.5% of individuals for whom data were missing for at least half the tests. We then imputed values for the remaining cases using IRMI (iterative robust model-based imputation; Templ, Kowarik, Filzmoser. 2011; Templ, Kowarik, Alfons, Prantner, 2019). The effects of age and sex on subtest scores have previously been detailed (Gur et al., 2012; Roalf et al., 2014). To handle non-linear effects, we residualized the subtest variables for age and sex using a natural (i.e., restricted cubic) spline model before performing factor analysis. We ran both exploratory factor analysis (EFA) and confirmatory factor analysis (CFA). The model with Social Cognition and the Sensorimotor Speed factors exhibited a lack of indicator coherence and did not converge. As such, we removed the five subtests related to these two factors and ran EFA/CFA for the 10 remaining ones. A model, based on the remaining tests, with Complex Cognition, Executive Control, and Episodic Memory factors as specified by Moore et al.’s (2015) five factor model also did not converge. However, we identified a similar three factor model (with analogous Executive Control, Episodic Memory, Complex Cognition broad factors), which had good fit among European and Hispanic Americans. This is shown in Figure 2. This same bifactor model also had a good fit among European and African Americans.

To ensure that the measure was unbiased between our major ethnic groups, we performed Multi-group confirmatory factor analyses (MGCFA) to determine if Measurement Invariance (MI) held for European and African Americans and for the European and Hispanic Americans (Dolan & Hamaker, 2001; Lubke, Dolan, Kelderman, & Mellenbergh, 2003). Results for European and African Americans were not interpretively different from those reported by Lasker et al. (2019), despite, in this case, using Full Information Maximum Likelihood (FIML) instead of listwise deletion to handle missing subtest scores. Results for European and Hispanic Americans are detailed in the Supplementary Material and also discussed below. The three-factor baseline model specified by Lasker et al. (2019) fit well in both groups. As detailed in Supplementary File 1, full factorial invariance held. Additionally, the weak form of Spearman’s hypothesis (Jensen, 1998), in which *g* accounts for the majority of variance in subtest scores, did not fit worse than baseline. In this model, 69-72% of the between-group differences are accounted for by *g.* In contrast, the contra models, in which the majority of subtest score differences were not attributable to *g*, and, arguably, the strong model, in which *g* accounts for all of variance in subtest scores, fit worse (see the detailed discussion in Supplemental File 1). The resulting mean differences for the weak Spearman’s hypothesis model are shown in Table 1, while results for the strong Spearman’s hypothesis model are provided in Supplementary File 1. Since it is often difficult to distinguish between Spearman Hypothesis models (e.g., strong vs. various forms of weak SH), and since the magnitude of *g* differences depends on the specification, for replicability and less dependence on researcher choice, we used *g*-scores from the exploratory factor analysis. These correlated at *r* = .97 with scores from the MGCFA model above.

**Table 1.** Factor Score Differences between Hispanic and European Americans based on the Weak Spearman’s Hypothesis Model.

| Factor | Estimate | SE | Lower 95% CI | Upper 95% CI |
| --- | --- | --- | --- | --- |
| $g$ | 0.668 | 0.064 | 0.543 | 0.793 |
| Complex Cognition | 0 | - | - | - |
| Executive Functioning | -0.342 | 0.124 | -0.585 | -0.099 |
| Episodic Memory | -0.234 | 0.075 | -0.382 | -0.087 |

*Note:* Positive values indicate higher European American scores and vice-versa. Estimates are in terms of Hedge's $g$ . Combined $N = 5,454$ with 515 Hispanic Americans and 4,939 European Americans.

As there was a relatively wide age range (8 to 22 years), on a reader’s request, we additionally investigated the validity of these scores across age groups. To do this we used parental education as a predictor and *g*-scores as the criterion. We used education as a predictor, since this variable is a known correlate of *g*, eduPGS, and genetic ancestry, and also since most cases had this variable. For this analysis, we limited the analytic sample to those with ancestry, *g*, and education. In Figure 3, we show the regression plot for parental education and *g* by age group, with participants grouped into three age distributions (for purposes of illustration). As is evident, age grouping has little effect. Moreover, in a regression model with parental education predicting *g*, the interaction term for age had a trivial, albeit statistically significant, effect (ß = =.008; *p* = .006; *N* = 7,846), likely due to the large sample size. Generally, parental education predicts *g*-scores more or less equally well across the age distribution, suggesting that our natural spline model effectively captured the age-related effects on subtests.

**Figure 3.**
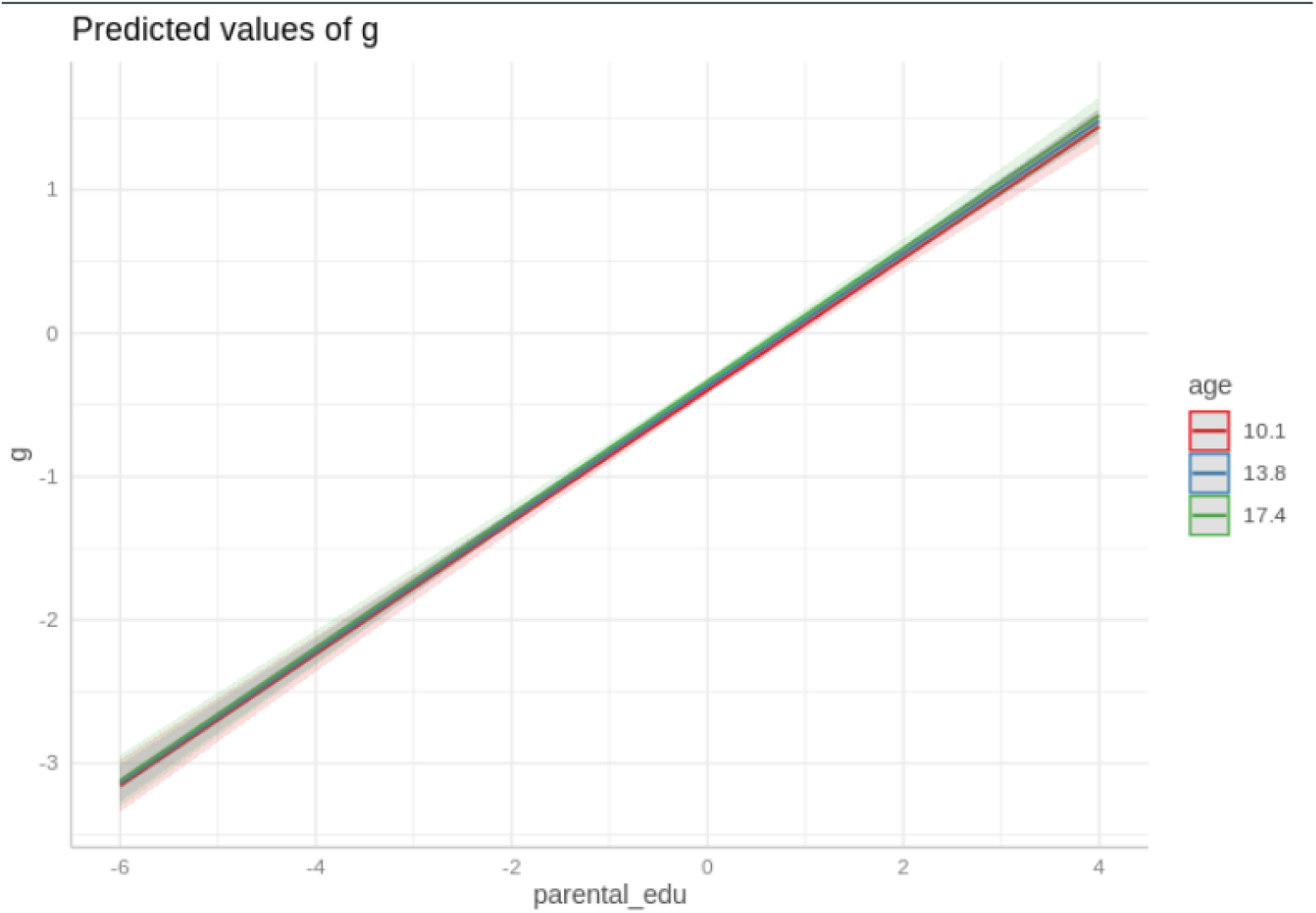
Regression Plot for Parental Education and g by Three Age Groups.

Means and standard deviations for *g* and the subtests are presented in Table 2. These (also including results for the five psychometrically biased subtests) are provided for the four main groups. Hispanic subgroup results, which were not used in any of the analyses in this paper, are additionally reported in Supplemental File 1

**Table 2.** General Intelligence (*g*) and Subtest Means and Standard Deviations for European (EA), European-African (EA-AA), African (AA), and Hispanic (HI) Americans.

|  | EA |  | EA-AA |  | AA |  | HI |  |
| --- | --- | --- | --- | --- | --- | --- | --- | --- |
| $g$ | 0.00 | 1.01 | -0.14 | 1.05 | -1.01 | 1.07 | -0.57 | 1.13 |
| PLOT* | -0.01 | 1.00 | -0.22 | 0.96 | -0.71 | 0.96 | -0.36 | 1.04 |
| PCPT* | 0.00 | 1.00 | -0.04 | 1.03 | -0.33 | 1.16 | -0.26 | 1.26 |
| PCET* | 0.00 | 1.00 | 0.03 | 0.97 | -0.45 | 1.08 | -0.26 | 1.08 |
| LNB* | 0.00 | 1.00 | -0.02 | 0.98 | -0.47 | 1.18 | -0.29 | 1.12 |
| VOLT* | 0.00 | 1.00 | -0.09 | 0.96 | -0.35 | 1.11 | -0.27 | 1.06 |
| TAP | 0.00 | 1.00 | 0.04 | 1.07 | -0.07 | 1.06 | 0.02 | 0.98 |
| PMRT* | 0.01 | 1.00 | -0.06 | 1.05 | -0.56 | 0.94 | -0.25 | 0.98 |
| MP | 0.00 | 1.00 | -0.16 | 1.10 | -0.11 | 1.04 | -0.07 | 1.18 |
| PEDT | 0.00 | 1.01 | -0.06 | 0.95 | -0.15 | 1.16 | -0.15 | 1.09 |
| PVRT* | 0.00 | 1.00 | -0.15 | 1.07 | -0.98 | 1.13 | -0.58 | 1.18 |
| PEIT | -0.01 | 1.01 | -0.02 | 1.07 | -0.05 | 1.10 | 0.02 | 1.02 |
| PFMT* | -0.01 | 1.00 | 0.10 | 1.03 | -0.04 | 1.08 | 0.06 | 1.04 |
| PADT | 0.00 | 1.01 | -0.06 | 0.93 | 0.00 | 1.12 | -0.06 | 1.02 |
| PWMT* | 0.00 | 1.01 | 0.07 | 1.03 | -0.17 | 1.23 | -0.11 | 1.16 |
| WRAT* | 0.00 | 1.00 | -0.13 | 1.08 | -0.85 | 0.95 | -0.48 | 1.05 |

*Note:* \*Denotes that the subtests were used in computing $g$ scores.

### 2.2 Parental Education

Following Lasker et al. (2019), we computed *z*-scores individually for paternal and maternal education and then averaged these. The average score was then *z*-scored again. In this sample, paternal and maternal education were the only available measures of socioeconomic status (SES). While this could be seen as a limitation, parental education is arguably the most relevant socioeconomic related variable as it has the strongest correlation with educational-based PGS (Lee et al., 2018; Figure 4) and as it is as highly correlated with cognitive ability / scholastic achievement as are other commonly used parental-based indicators, such as occupation and income (Sirin, 2005; Table 3). That said, parental education is not a complete measure of SES, and we do not treat it as such. To note, using a much broader index of SES, but a narrower index of cognitive ability, Guo, Lin, and Harris (2019; Table 3) report descriptively similar results, though the effect of eduPGS on ability is attenuated more among Black and Black Hispanics when controlling for these variables.

**Figure 4.**
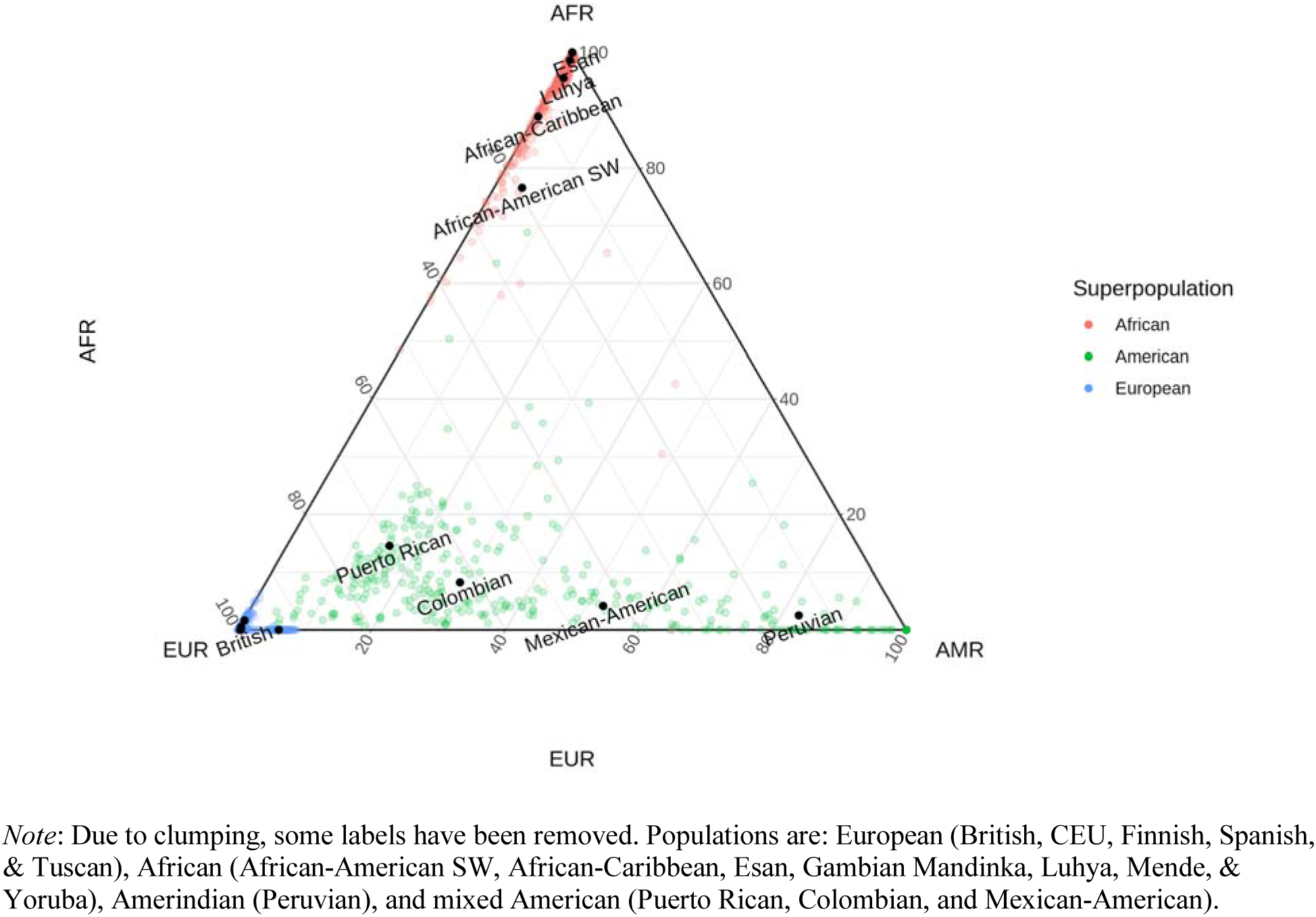
Admixture Ternary-Plot for 1000 Genomes Reference Samples.

**Table 3.**
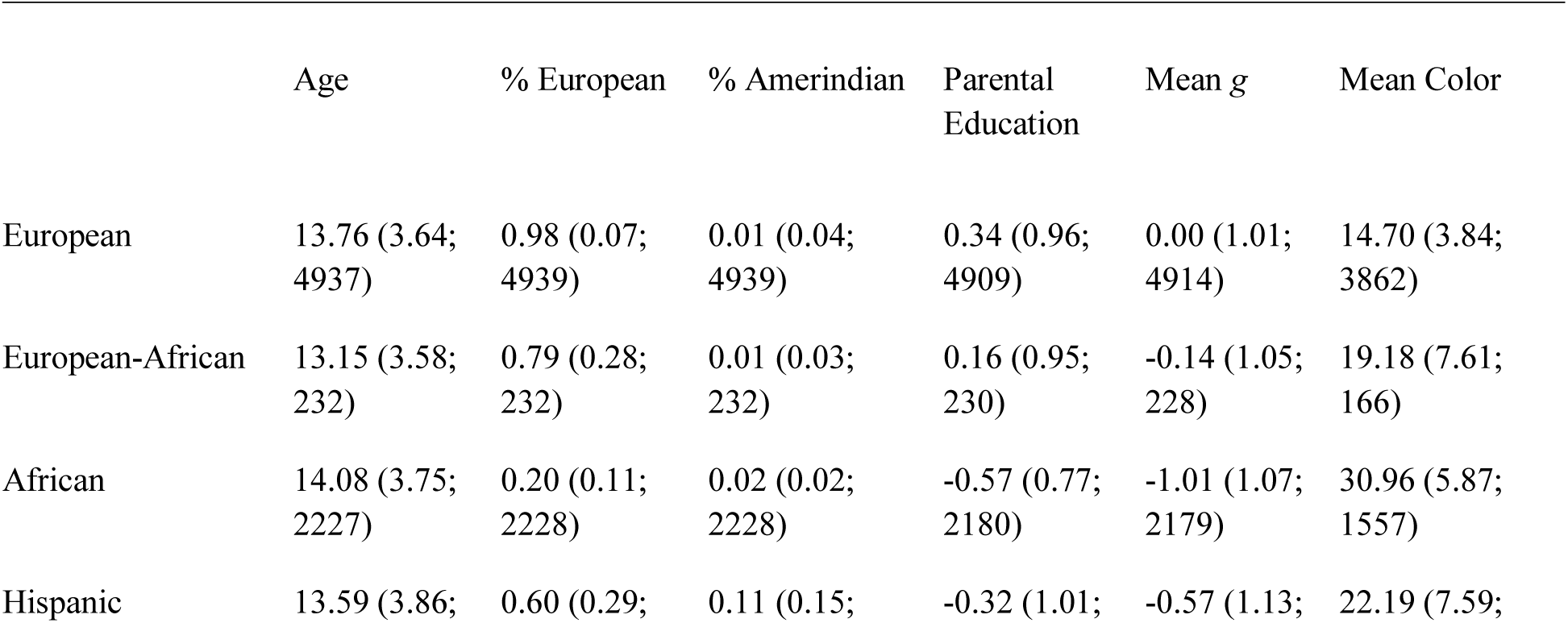
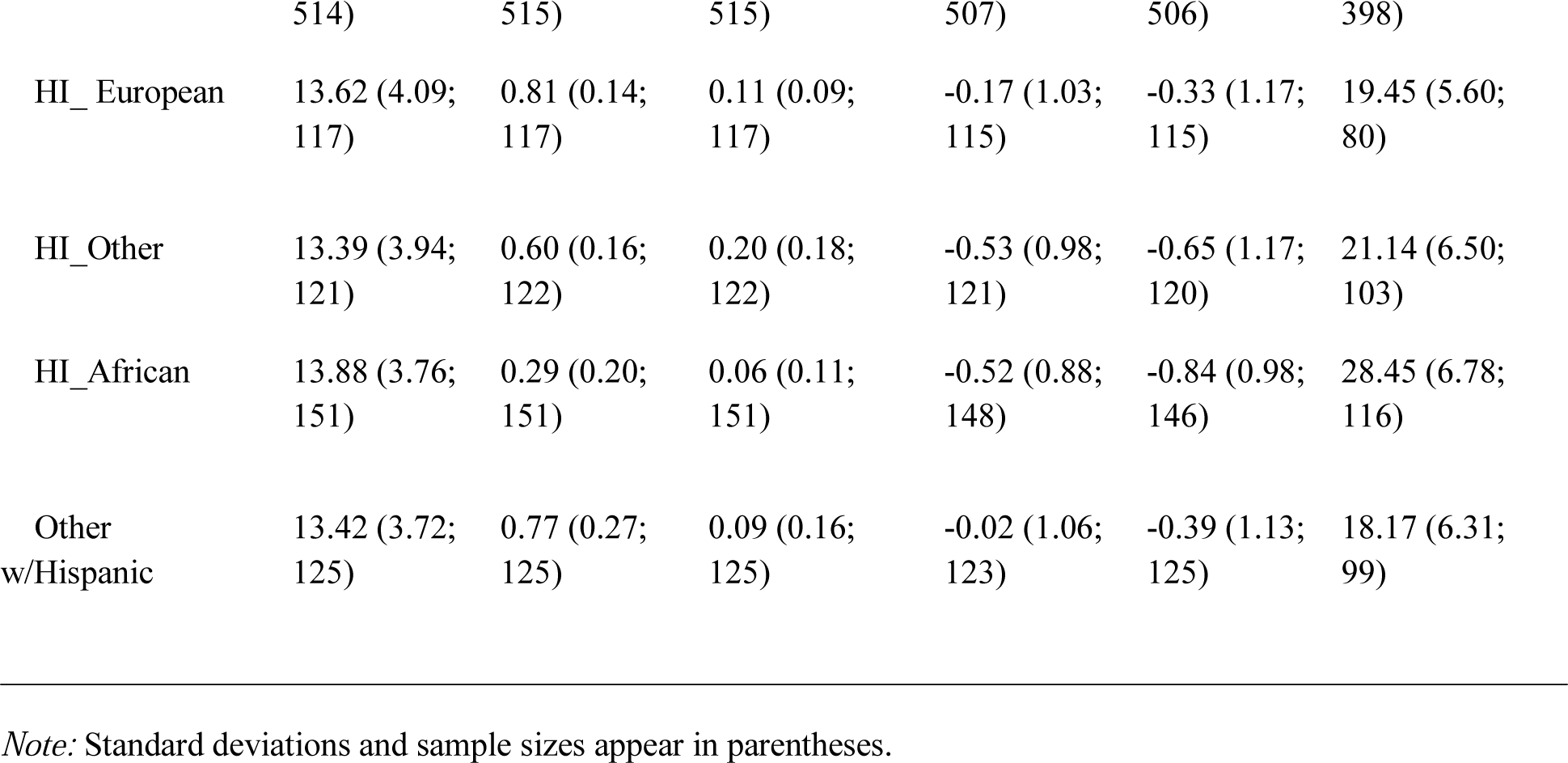
Sample Characteristics for the Participants.

### 2.3 SIRE

Self-identified race/ethnicity (SIRE) was based on yes/no questions for which the participants could select multiple races or ethnicities out of the following set of choices: Black or African-American; American Indian or Alaskan Native; Asian; European-American; Hispanic/Latino; Native Hawaiian/Pacific Islander; Other; and Not Available/Pending Validation. Those who identified as European only and did not report being Hispanic were coded as European American, while those who identified as African only and did not report being Hispanic were coded as African American, and those who identified as both European and African and did not report being Hispanic were coded as European-African American. Those who identified as Hispanic, regardless of self-identified race, were coded as such. Among Hispanics, we also identified individuals by self-identified race: European, African, Other, and a mixed category of all others (e.g., European-African Hispanic).

### 2.4 Genetic Ancestry Percentages

Lasker et al. (2019) previously described the TCP genotype data. Different arrays covered different variants, so to obtain overlapping sets of single nucleotide polymorphisms (SNPs), we imputed variants with the Michigan Imputation Server (https://imputationserver.sph.umich.edu/index.html). We used this server with the Minimac3 imputation algorithm, 1000G Phase 3 v5 as reference panel, and Eagle v2.3 Phasing. For computational efficiency when calculating admixture percentages, we filtered the 15.5 M variants available to the 6.5 M with a minor allele frequency (MAF) of at least 0.05. For color scores (based on a total possible of 35 variants), we did not filter by MAF, so as not to lose variants. Imputation was done using PLINK v1.90b6.8 (Chang et al., 2015). For individual ancestry estimates, we used ADMIXTURE version 1.3.0 D.H. (Alexander, Novembre, & Lange, 2009). Prior, we merged the TCP with 1000 Genomes reference samples: European (British, CEU, Finnish, Spanish, & Tuscan), African (African-American SW, African-Caribbean, Esan, Gambian Mandinka, Luhya, Mende, & Yoruba), Amerindian (Peruvian), and mixed American (Puerto Rican, Colombian, and Mexican-American). We did this to gain a reliable estimate of Amerindian ancestry, since Hispanics in this sample were a three-way admixed population. We then ran ADMIXTURE with k = 3 genetic clusters. The results for the 1000 Genomes reference populations appear in Figure 4, while those for the TCP European, European-African, African, and Hispanic samples appear in Figure 5. Note that the Hispanics in the sample were largely an Afro-European descent group, with an ancestry profile matching the predominantly Puerto Rican origin of the Philadelphia Hispanic population (69% Puerto Rican; 7% Mexican; 2% Cuban; 22% Other; US Census Bureau, 2016).

**Figure 5.**
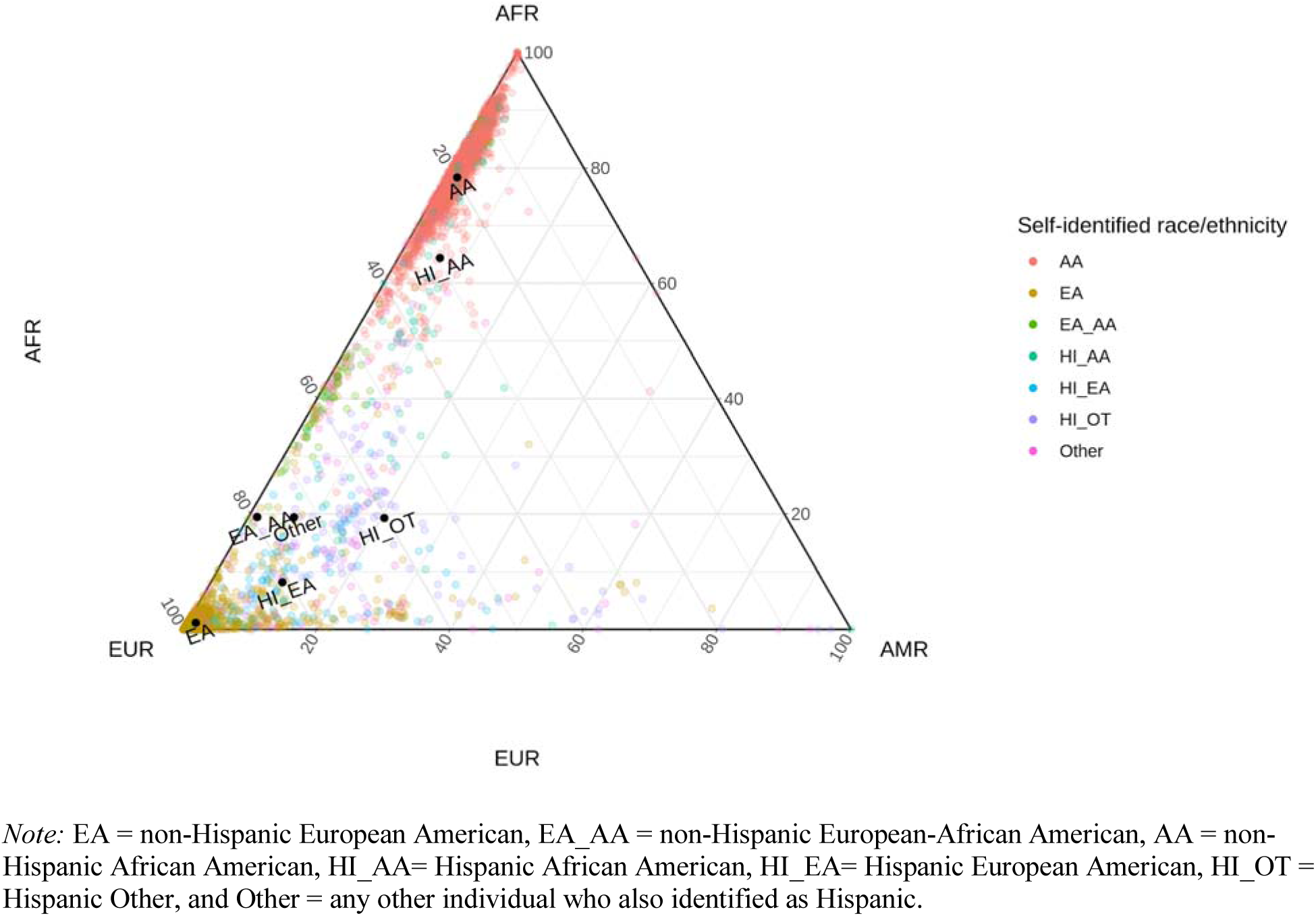
Admixture Ternary-Plot for TCP Self-identified SIRE Groups.

### 2.5 Skin, Hair, and Eye Color

Because the data did not include measures of appearance, we followed Lasker et al. (2019) and calculated phenotypic scores from genotypic data. Namely, we imputed phenotype based on genotype using the HIrisPlex-S web application (https://hirisplex.erasmusmc.nl/). Developed for use by the U.S. Department of Justice in forensic investigations, and validated on thousands of people from around the world (Chaitanya et al., 2018), the HIrisPlex-S imputes skin, hair, and eye color probabilities from 41 SNPs (with overlaps: 6 for eye color, 22 for hair color, and 36 for skin color), with a high degree of accuracy. We focus on skin color since this trait is given primacy by proponents of race-associated phenotypic discrimination (“colorism”) models (e.g., Dixon & Telles, 2017; Marira & Mitra, 2013), and since the other traits have a higher rate of missing data, owing to poor tagging of SNPs in some of the microarrays.

HIrisPlex-S gives probabilities of The Fitzpatrick Scale skin type, which range from Type I “palest; freckles” to Type VI “deeply pigmented dark brown to darkest brown”. The Fitzpatrick Scale skin types correspond with scores on Von Luschan’s chromatic scale. For example, a Fitzpatrick Scale Type I classification corresponds with a Von Luschan’s chromatic scale score of 0–6. This correspondence allowed us to transform the HIrisPlex-S skin type probabilities into a single, color measure. This was done by weighting the median score of each color type by the HIrisPlex-S predicted probability of each type. Owing to poor tagging of SNPs in the arrays, it was possible to compute color scores for only 3,862 European, 166 European-African, 1,557 African, and 398 Hispanic Americans.

The correlation between skin color and European ancestry for the combined sample was *r* = -.87 (*N* = 6,050), as shown in Figure 6. For Hispanics alone, the correlation was *r* = -.67 (*N* = 398), as shown in Figure 7. Note, the expected correlations are population specific, owing to differences in admixture range, admixture components, assortative mating, etc. (Kim, Edge, Goldberg, & Rosenberg, 2019). In this case, the estimate found for Philadelphian Hispanics is similar to those reported by others for comparable populations (e.g., Puerto Rican: rho = .63; Parra, Kittles, & Shriver, 2004; R-square = .417 / *r* = .65; Bonilla, Shriver, Parra, Jones, & Fernández, 2004). For the African American only (monoracial) sample, the *r* = .39 was also similar to previously reported comparable ones (e.g., African Americans from Washington, D.C.: *r*_s_ = .44; Parra, Kittles, & Shriver, 2004), so it is likely that our color estimates are reasonably precise.

**Figure 6.**
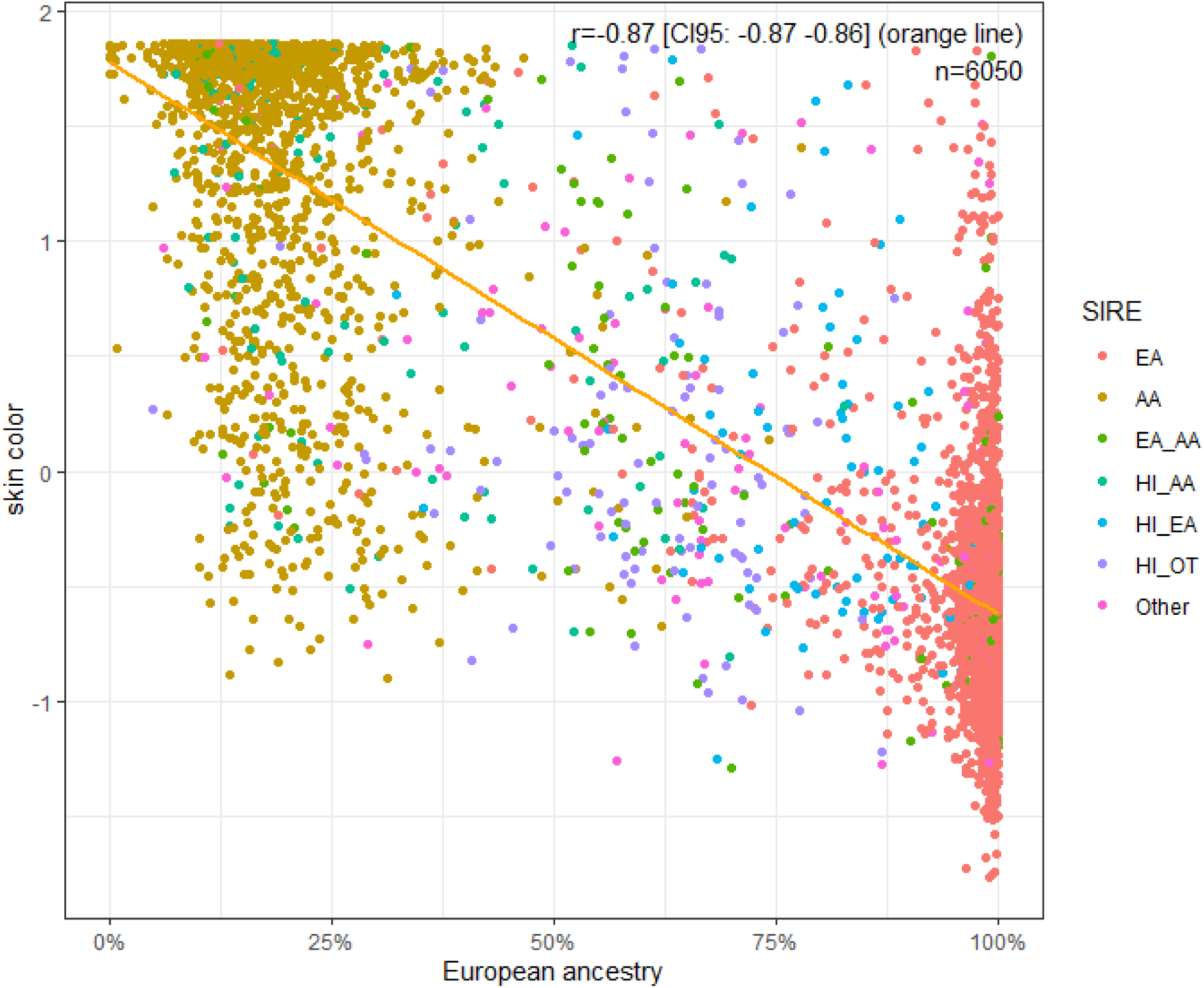
Regression Plot of the Relation Between Color (with Higher Values indicating Darker Color) and European Genetic Ancestry in the Combined Sample.

**Figure 7.**
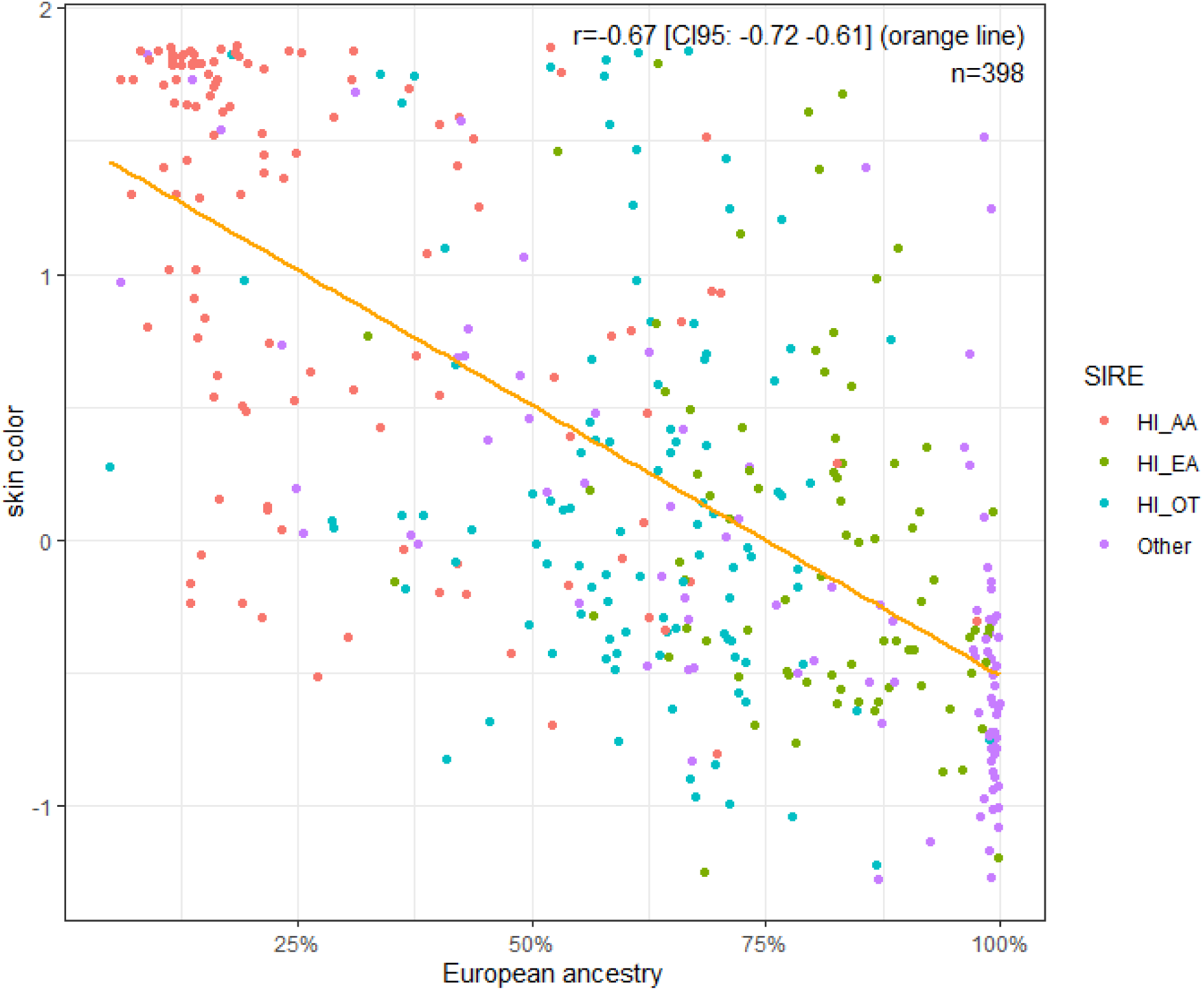
Regression Plot of the Relation Between Color (with Higher Values indicating Darker Color) and European Genetic Ancestry Among Hispanics.

Further, consistent with observations previously reported (e.g., Bonilla et al., 2004), the mean score for Hispanics was 22.19 which is intermediate to Type III (sometimes mild burn, tans uniformly) and Type IV (burns minimally, always tans well, moderate brown). Moreover, Hispanics who identified as European had a color score of 19.45, which was significantly darker than that for non-Hispanic European Americans at 14.70 (*t* (3,940) = 10.645). And, Hispanics who identified as African had a color score of 28.45, which was significantly lighter than that for non-Hispanic African Americans at 30.96 (*t* (1,671) = -4.393). Generally, these color values are consistent with known population values.

### 2.6 Cognitive Ability-Related Polygenic Scores

Lasker et al. (2019) detailed the rationale for eduPGS selection. Briefly here, we used four overlapping eduPGS from Lee et al. (2018): The eduPGS with all variants trained without the 23andMe cohort (with *N* = 7,762,369 SNPs in the present dataset), the multi-trait analysis of genome-wide association study (MTAG) eduPGS 10k SNPs (with *N* = 8,442 SNPs in the present dataset), the MTAG eduPGS lead SNPs (with *N* = 1,558 SNPs in the present dataset), and finally Lee et al.’s (2018) putatively causal variants (with *N* = 111 SNPs in the present dataset).

The eduPGS with all variants is predicted to show high within discovery population validity, but poor validity in non-discovery populations owing to high LD decay bias as a result of including a large number of SNPs, irrespective of their significance (Zanetti & Weale, 2018). Both the MTAG eduPGS 10k SNPs and MTAG-lead SNPs are predicted to show moderate within-discovery population validity and moderate validity in non-discovery populations. This is because ’lead’ or ’clumped’ SNPs with greater statistical significance are more likely to be causal or very close to a causal variant, which are also more likely to be transethnically valid (Marigorta & Navarro, 2013; Spencer, Cox, & Walters, 2014; Wang & Teo, 2015; Grinde et al., 2018). Finally, because causal SNPs are more likely to be transethnically valid, Lee et al.’s (2018) putative causal SNPs are likely to have the highest transethnic validity. However, as there are only 111 of these, the validities are predicted to be low in both discovery and non-discovery populations (e.g., Lasker et al., 2019).

In a supplementary analysis, we additionally examined the effect of computing MTAG 10k eduPGS using population-GWAS versus within family Beta weights for the SNPs. For the population-GWAS weights, we used Lee et al.’s (2018) published MTAG weights computed based on 1.1 million individuals. For the within family weights, we contacted Lee et al. (2018), who provided us with the within family weights from their analyses of 22,000 sibling pairs. For these analyses, we computed eduPGS using the subset of MTAG SNPs for which there were both within and population-GWAS weights.

## 3 Results

### Descriptive Statistics

Descriptive statistics for all groups appear in Table 3. Self-identifying European, European-African, African, and Hispanic Americans were ancestrally 98%, 79%, 20%, and 60% European, respectively. Further, among Hispanics, those who identified as European were 81% ancestrally European, while those who identified as African were 29% ancestrally European. This difference in ancestry is consistent with those reported previously (e.g., Table S5; Bryc et al., 2015). The Hispanic_Other group had the largest percentage of Amerindian ancestry, at 20%, which would be consistent with origins in Central or South American, as opposed to the Caribbean (Table S5; Bryc et al., 2015).

The Hispanic/European difference in *g* came to *d* = 0.56 (European vs. Hispanic: IQ = 100.00 versus IQ = 91.63). This difference is somewhat smaller than typically reported (e.g., Roth et al., 2017). The discrepancy may have resulted because inclusion into the TCP sample was limited to English speaking individuals. Among Hispanics, the difference in *g* between SIRE Europeans and Africans was *d* = 0.50 (i.e. European-Hispanic vs. African-Hispanic: IQ = 95.16 vs. IQ = 87.67). This is consistent with results typically reported in the literature (e.g., Kirkegaard & Fuerst, 2016). In comparison, the difference in *g* between European and African Americans came out to *d* = 0.98 (European vs. African American: IQ = 100.00 versus IQ = 85.27), while that between European and European-African American came out to *d* = 0.14 (European vs. European-African American: IQ = 100.00 versus IQ = 97.90).

### 3.2 Bivariate Relationships Among the Variables for Hispanics

We first report the bivariate correlations. These allow comparison with effects sizes for the association between ancestry and SES reported previously (e.g., Kirkegaard, Wang, & Fuerst, 2017). Table 4 shows the correlations for Hispanics. Consistent with having a predominantly Puerto Rican sample (e.g., Via et al., 2011), Amerindian ancestry was only weakly (negatively) correlated with European ancestry. Cognitive ability and parental education were positively related to European ancestry; negatively related to African ancestry, and unrelated to Amerindian ancestry. Notably, the correlations between cognitive ability, parental education, and European and African ancestry are higher than reported by Lasker et al. (2019) for their monoracial African American sample. This is likely due to the greater variance in admixture among Hispanics in this sample (*SD*_European_ = 29%). The correlation between European ancestry and parental education in this sample was also higher than that reported for European ancestry and socioeconomic status among Puerto Ricans (*r* = .16, *K* = 3, *N* = 1,943; Kirkegaard, Wang, & Fuerst, 2017).

**Table 4.** Pairwise Correlations among Self-Identified Hispanic-Americans.

|  | Cognitive Ability | Parental Education | Euro. Ancestry | Afr. Ancestry | Amer. Ancestry | SIRE European | Color | EduPGS |
| --- | --- | --- | --- | --- | --- | --- | --- | --- |
| Cognitive Ability | 1<br>(506) |  |  |  |  |  |  |  |
| Parental Education | .287*<br>(500) | 1<br>(507) |  |  |  |  |  |  |
| European Ancestry | .300*<br>(506) | .239*<br>(507) | 1<br>(515) |  |  |  |  |  |
| African Ancestry | -.294*<br>(506) | -.210*<br>(507) | -.874*<br>(515) | 1<br>(515) |  |  |  |  |
| Amerindian Ancestry | .015<br>(506) | -.035<br>(507) | -.160*<br>(515) | -.340*<br>(515) | 1<br>(515) |  |  |  |
| SIRE: European | .199*<br>(506) | .223*<br>(507) | .643*<br>(515) | -.544*<br>(515) | -.139*<br>(515) | 1<br>(515) |  |  |
| Color | -.161*<br>(391) | -.077<br>(392) | -.670*<br>(398) | .637*<br>(398) | -.009<br>(398) | -.409*<br>(398) | 1<br>(398) |  |
| EduPGS | .293*<br>(506) | .330*<br>(507) | .557*<br>(515) | -.544*<br>(515) | .027<br>(515) | .355*<br>(515) | -.385*<br>(398) | 1<br>(515) |

Note: \*Significant at $p < 0.01$ . Pairwise $N$ in parentheses; EduPGS = MTAG 10k EduPGS.

Table 5 additionally shows the correlations for the combined group. As expected, the correlations for European ancestry are higher in the combined sample, since there is greater variability. The correlation between European ancestry and parental education was also higher than that between European ancestry and SES (*r* = .17, *K* = 15, *N* = 15,980.50; Kirkegaard, Wang, and Fuerst, 2017) previously reported for multi-ethnic and/or unspecified North and Latin American samples. This may again be due to this sample’s higher variability in ancestry.

**Table 5.** Pairwise Correlations among all participants in this sample.

|  | Cognitive Ability | Parental Education | Euro. Ancestry | Afr. Ancestry | Amer. Ancestry | SIRE European | Color | EduPGS |
| --- | --- | --- | --- | --- | --- | --- | --- | --- |
| Cognitive Ability | 1<br>(7,920) |  |  |  |  |  |  |  |
| Parental Education | .401**<br>(7,846) | 1<br>(7,846) |  |  |  |  |  |  |
| European Ancestry | .405**<br>(7,920) | .403**<br>(7,921) | 1<br>(8,009) |  |  |  |  |  |
| African Ancestry | -.406**<br>(7,920) | -.400**<br>(7,921) | -.988**<br>(8,009) | 1<br>(8,009) |  |  |  |  |
| Amerindian Ancestry | -.025*<br>(7,920) | -.047**<br>(7,921) | -.150**<br>(8,009) | -.003<br>(8,009) | 1<br>(8,009) |  |  |  |
| SIRE: European | .396**<br>(7,920) | .405**<br>(7,921) | .938**<br>(8,009) | -.930**<br>(8,009) | -.124**<br>(8,009) | 1<br>(8,009) |  |  |
| Color | -.350**<br>(5,991) | -.341**<br>(5,989) | -.865**<br>(6,050) | .859**<br>(6,050) | .102**<br>(6,050) | -.806**<br>(6,050) | 1<br>(6,050) |  |
| EduPGS | .399**<br>(7,920) | .442**<br>(7,921) | .667**<br>(8,009) | -.666**<br>(8,009) | -.058**<br>(8,009) | .629**<br>(8,009) | -.603**<br>(6,050) | 1<br>(8,009) |

Note: \*\*Values are significant at $p < 0.0001$ ; \*values are significant at $p < 0.05$ . Pairwise $N$ in parentheses; EduPGS = MTAG 10k EduPGS.

Finally, the association between cognitive ability and European ancestry for the combined sample is depicted in Figure 8. A Bayesian Generalized Additive Model line (in blue) is superimposed on a regression line (orange); moreover, self-identified racial groups are color coded. As can be seen, the association is nearly linear. Additionally, European ancestry is positively and significantly associated with *g* in the admixed SIRE groups (*r* _European-African_ = .26, *r*_African_ = .084, *r*_Hispanic_ = .30), though not for European Americans (*r*_European_ = .02) among whom there is little non-European admixture.

**Figure 8.**
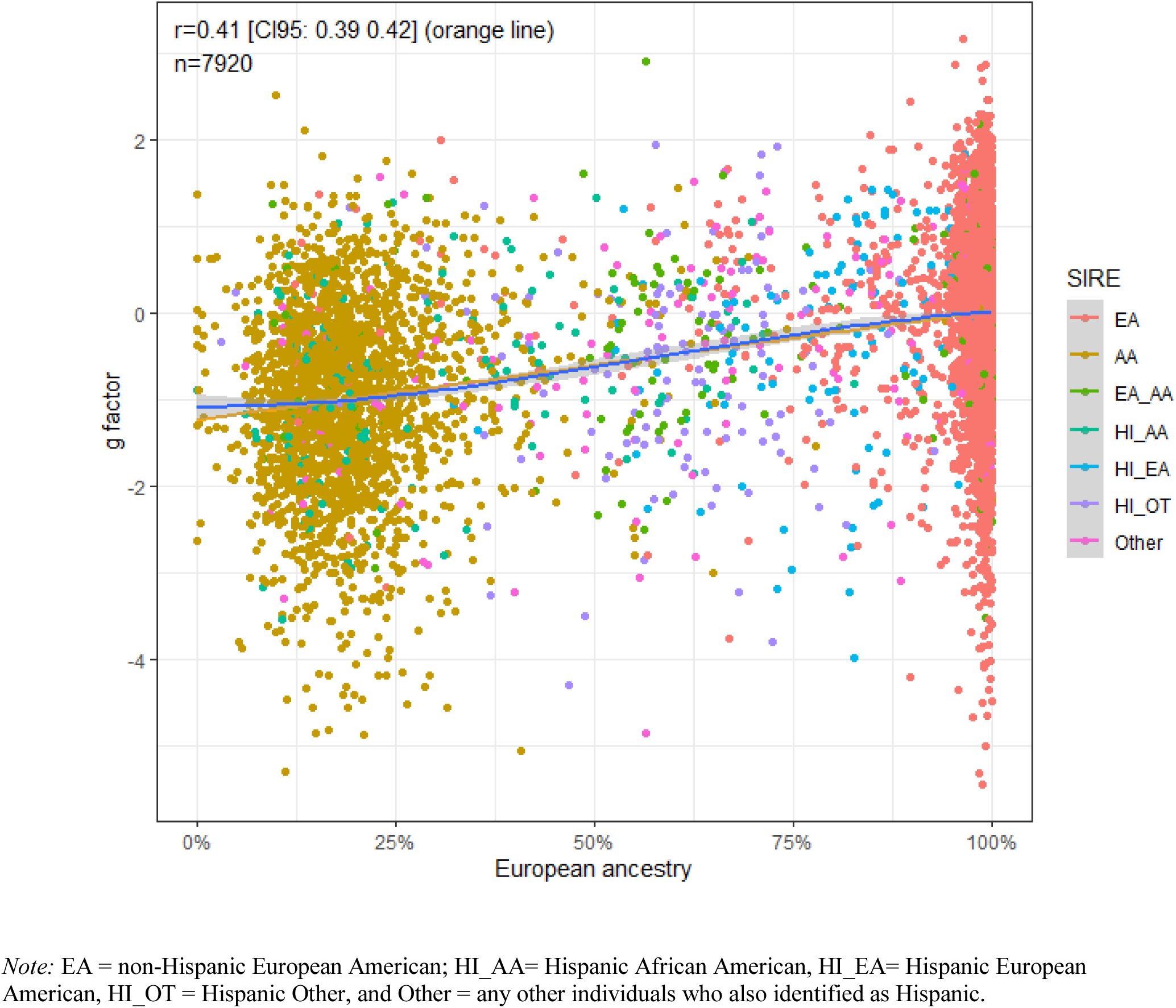
Regression Plot of the Relation Between g and European Genetic Ancestry in the Combined Sample.

**Figure 9.**
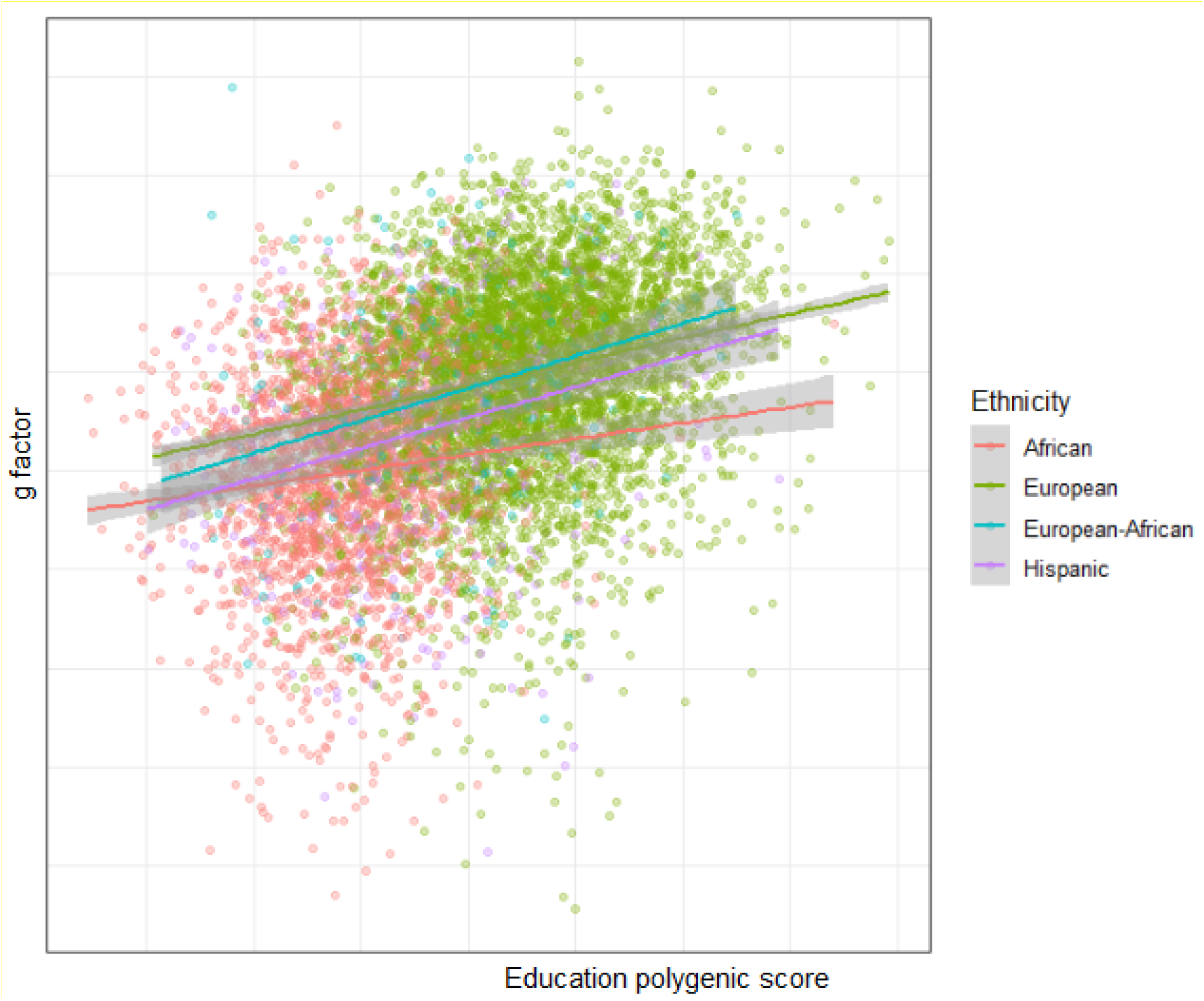
Regression Plot for the Predictive Validity of MTAG 10k eduPGS with Respect to g in the Hispanic (Purple), European (Green), European-African (Blue), and African American (Red) Samples.

### 3.3 Regression Analyses

Multiple regression analysis is preferable to bivariate analysis, since there is a possibility of confounding with social and environmental factors, particularly ones correlated with SIRE (Fang et al., 2019) and race-associated phenotype (Conley & Fletcher, 2017), and also of a non-independence of admixture components. In the initial regression analysis, shown in Table 6, we deal exclusively with those who identify as Hispanic American. In the first, Model (1a), we include only African and Amerindian ancestry as independent variables. In Model 1b, we add self-identified race to see if ancestry retains predictivity and has independent explanatory power. In Model 2, we approach the issue from the perspective of “colorism” (that is, discrimination based on race-associated phenotype, specifically color). As such, in Model 2a, we include only color. We then add ancestry in 2b to see which variables retain validity. In Model 2c we further add parental education to the model with color and ancestry. Note, all values except genetic ancestry are standardized. This allows the Beta coefficients for ancestry to be interpreted as a change in a standard deviation of cognitive ability going from 0% to 100% ancestry. Doing so allows results from different samples with different variances in ancestry to be compared on the same metric.

**Table 6.**
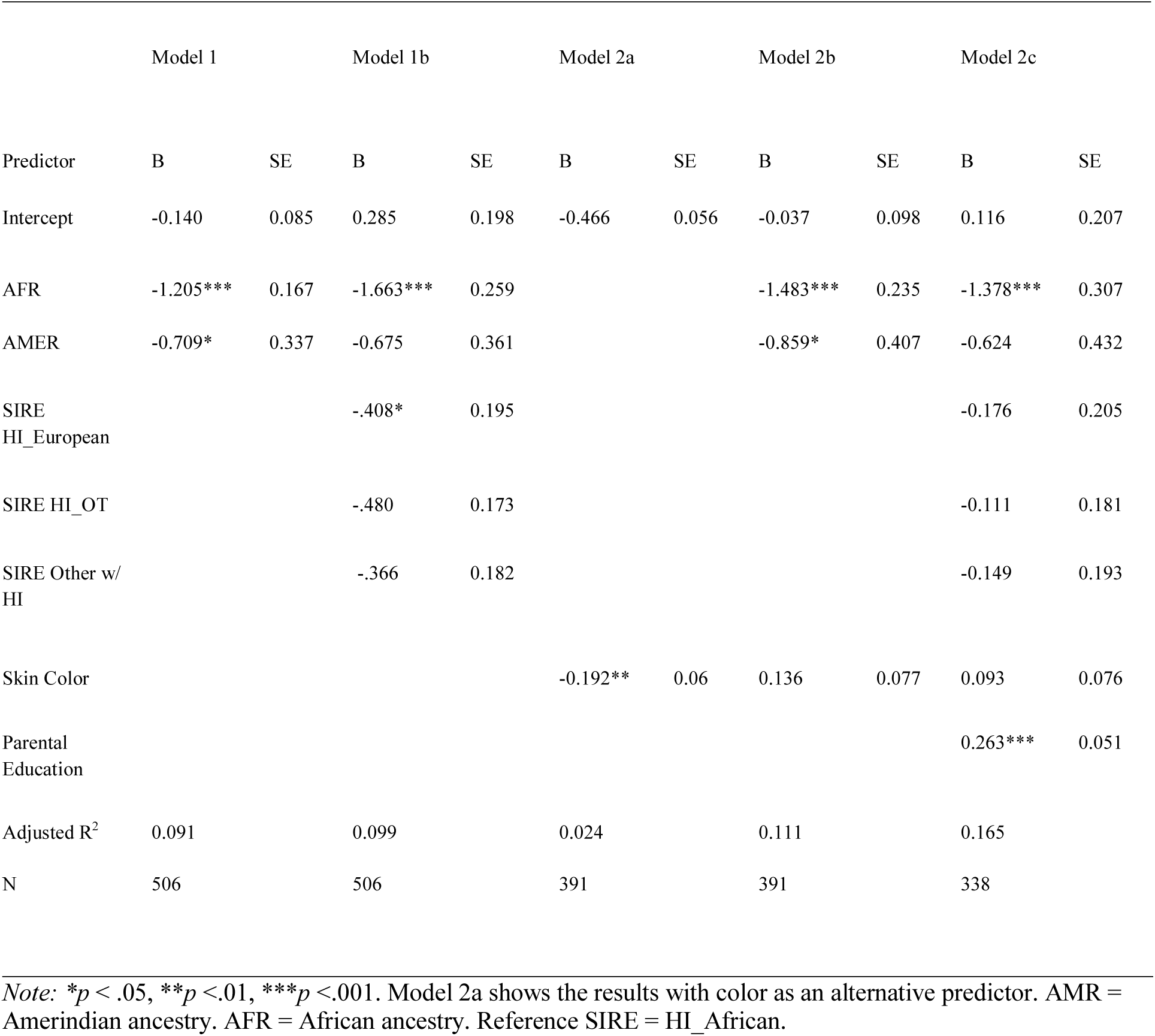
Regression Analysis for European Ancestry as a Predictor of *g* among Hispanic Americans with Controls for Skin Color (Model 2b), and Parental Education (Model 2c) Added.

As can be observed, African ancestry retained validity when either SIRE or color were added. In contrast, with the inclusion of SIRE, Amerindian ancestry became a nonsignificant predictor. However, the direction and magnitude of the effect was as previously reported (Kirkegaard et al., 2019), and the non-significance may simply be due to a lack of statistical power, given the low variance in Amerindian ancestry. As seen in Model 2a, darker color was a significant predictor of lower cognitive ability, consistent with previously reported findings (e.g., Hu, Lasker, Kirkegaard, & Fuerst, 2019). However, the effect became nonsignificant and the sign reversed with the inclusion of ancestry. This is consistent with the findings of Lasker et al. (2019), and generally supports the view that the association between color and ability is simply indexing that between ancestry and ability. Note, Model 1 and Model 2 above have different numbers, since there were fewer cases with color data. However, running the results with the same sample sizes (listwise deletion) did not lead to a difference of interpretation.

Next, we repeat the analysis with the other groups (EA, EA-AA, and AA). The results appear in Table 7, and are descriptively similar to those for the Hispanic-only sample. As before, Amerindian ancestry loses validity on inclusion of SIRE. This time, however, the effect size of Amerindian ancestry is close to zero. Statistically, this happens because among Philadelphian Europeans, the association between Amerindian ancestry and ability is non-significantly positive (*B* = 0.500, S.E. = 0.391, *N* = 4914, *p* = .20), while the association between African ancestry and ability is significantly negative (*B* = -0.616, S.E. = 0.266, *N* = 4914, *p* < .05). Why this is the case is not clear. In Model 2a, color alone has validity, but it becomes nonsignificant and changes direction on inclusion of ancestry (2b). This remains the case when parental education is added in model 2c.

**Table 7.** Regression Analysis for European Ancestry as a Predictor of *g* in the combined sample with Controls for Skin Color (Model 2), and Parental Education (Model 3) Added.

|  | Model 1a |  | Model 1b |  | Model 2a |  | Model 2b |  | Model 2c |  |
| --- | --- | --- | --- | --- | --- | --- | --- | --- | --- | --- |
| Predictor | B | SE | B | SE | B | SE | B | SE | B | SE |
| Intercept | 0.009 | 0.015 | 0.010 | 0.015 | -0.291*** | 0.013 | 0.016 | 0.023 | -0.089 | 0.023 |
| AFR | -1.274*** | 0.032 | -0.988*** | 0.108 |  |  | -1.262*** | 0.073 | -0.785*** | 0.129 |
| AMER | -0.532* | 0.208 | 0.043 | 0.242 |  |  | -0.299 | 0.253 | 0.148 | 0.281 |
| SIRE<br>African |  |  | -0.248* | 0.088 |  |  |  |  | -0.107 | 0.095 |
| SIRE<br>European_African |  |  | 0.042 | 0.072 |  |  |  |  | 0.044 | 0.081 |
| SIRE_<br>HI_European |  |  | -0.2632** | 0.100 |  |  |  |  | -0.091 | 0.115 |
| SIRE<br>HI_African |  |  | -0.208 | 0.112 |  |  |  |  | -0.098 | 0.121 |
| SIRE<br>HI_OT |  |  | -0.4823*** | 0.109 |  |  |  |  | -0.127 | 0.115 |
| SIRE<br>Other w/ HI |  |  | -0.097 | 0.076 |  |  |  |  | 0.035 | 0.082 |
| Skin Color |  |  |  |  | -0.390*** | 0.014 | -0.003 | 0.026 | 0.002 | 0.025 |
| Parental<br>Education |  |  |  |  |  |  |  |  | 0.331*** | 0.014 |
| Adjusted R2 | 0.165 |  | 0.168 |  | 0.122 |  | 0.164 |  | 0.237 |  |
| N | 7920 |  | 7920 |  | 5991 |  | 59991 |  | 5940 |  |

*Note:* $*p < .05$ , $**p < .01$ , $***p < .001$ . Model 1b shows the results with color as an alternative predictor. AMR = Amerindian ancestry. AFR = African ancestry. Reference SIRE = European.

### 3.4. Cognitive Ability and Education-related PGS (eduPGS)

We next report bivariate associations between cognitive ability, parental education, and four eduPGS from Lee et al. (2018). Hispanic results appear in Table 8. And the results for European, Europe-African, and African Americans are shown in Tables 9 to 11. In the Hispanic-only sample, all but the putatively causal eduSNPs were significantly associated with cognitive ability. Moreover, there were no statistically significant differences in the magnitudes of the correlations between Hispanic and European Americans for the putative causal eduPGS, the GWAS eduPGS, and the MTAG 10K PGS. However, the MTAG-lead PGS were significantly more predictive for Hispanics (*z* = 2.11, two-tail-*p* = 0.0349). In the case of GWAS eduPGS, the lack of difference in predictivity is somewhat surprising, since this is predicted to show high LD decay, and thus relatively low predictivity in non-European samples. It is possible that the association is spuriously high in the Hispanic sample owing to confounding with genetic ancestry. Among both European and European-African Americans, all eduPGS were significantly predictive of *g*. Since the European-African group was 79% European in ancestry, the effect of LD decay may have been minimal. For African Americans, in contrast, the validities were markedly reduced (e.g., MTAG 10k: *r*_European_ = .227 vs. *r*_African_ = .112). This is consistent with the general finding of reduced PGS validity among Afro-descent groups (Duncan et al., 2019).

**Table 8.**
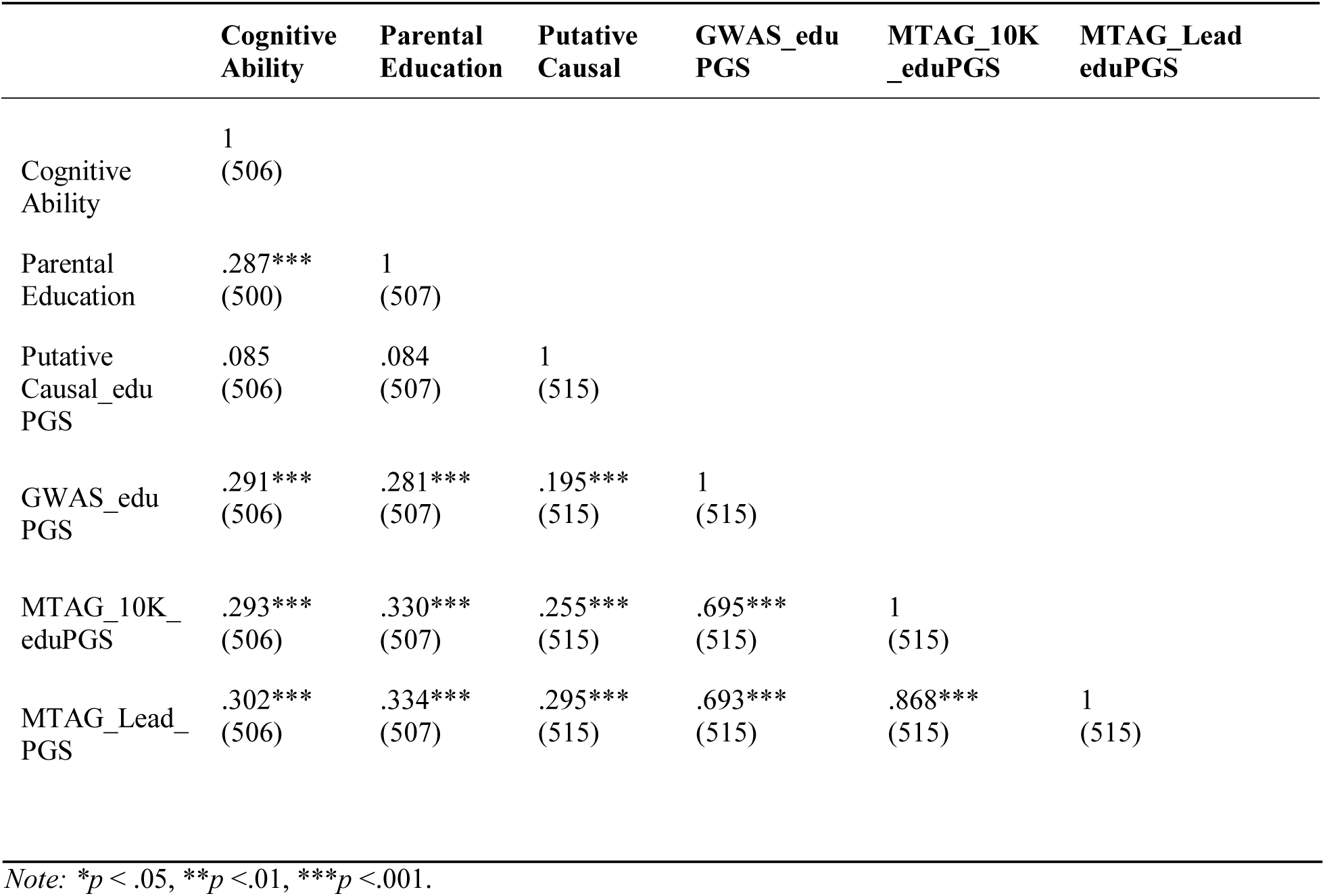
Pairwise Correlations Between Cognitive Ability and Education/Intelligence Related Polygenic Scores among Hispanic-Americans.

**Table 9.** Pairwise Correlations Between Cognitive Ability and Education/Intelligence Related Polygenic Scores among European-Americans.

|  | Cognitive Ability | Parental Education | Putative Causal | GWAS_edu PGS | MTAG_10K_eduPGS | MTAG_Lead_eduPGS |
| --- | --- | --- | --- | --- | --- | --- |
| Cognitive Ability | 1<br>(4914) |  |  |  |  |  |
| Parental Education | .297***<br>(4886) | 1<br>(4909) |  |  |  |  |
| Putative Causal_edu PGS | .058***<br>(4914) | .094***<br>(4909) | 1<br>(4939) |  |  |  |
| GWAS_edu PGS | .226***<br>(4914) | .306***<br>(4909) | .217***<br>(4939) | 1<br>(4939) |  |  |
| MTAG_10K_eduPGS | .227***<br>(4914) | .288***<br>(4909) | .315***<br>(4939) | .646***<br>(4939) | 1<br>(4939) |  |
| MTAG_Lead_PGS | .210***<br>(4914) | .249***<br>(4909) | .348***<br>(4939) | .575***<br>(4939) | .837***<br>(4939) | 1<br>(4939) |

Note: \* $p < .05$ , \*\* $p < .01$ , \*\*\* $p < .001$ .

**Table 10.** Pairwise Correlations Between Cognitive Ability and Education/Intelligence Related Polygenic Scores among European-African Americans.

|  | **Cognitive Ability** | **Parental Education** | **Putative Causal** | **GWAS_edu PGS** | **MTAG_10K_eduPGS** | **MTAG_Lead_eduPGS** |
| --- | --- | --- | --- | --- | --- | --- |
| Cognitive Ability | 1<br>(228) |  |  |  |  |  |
| Parental Education | .391***<br>(227) | 1<br>(230) |  |  |  |  |
| Putative Causal_edu PGS | .170*<br>(228) | .201*<br>(230) | 1<br>(232) |  |  |  |
| GWAS_edu PGS | .302***<br>(228) | .400***<br>(230) | .276***<br>(232) | 1<br>(232) |  |  |
| MTAG_10K_eduPGS | .308***<br>(228) | .381***<br>(230) | .453***<br>(232) | .734***<br>(232) | 1<br>(232) |  |
| MTAG_Lead_PGS | .312***<br>(228) | .409***<br>(230) | .462***<br>(232) | .736***<br>(232) | .895***<br>(232) | 1<br>(232) |

Note: \* $p < .05$ , \*\* $p < .01$ , \*\*\* $p < .001$ .

**Table 11.**
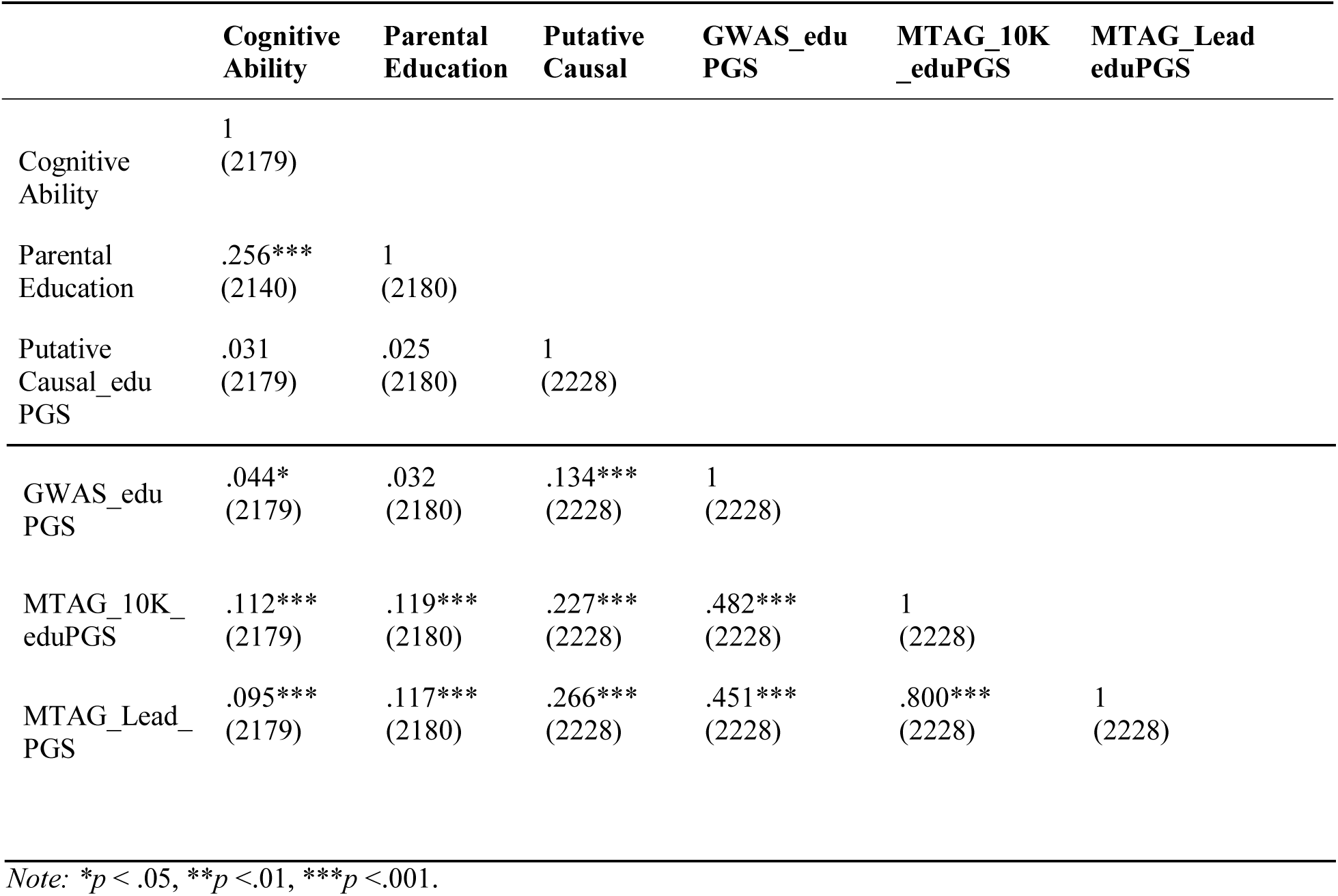
Pairwise Correlations Between Cognitive Ability and Education/Intelligence Related Polygenic Scores among African-Americans.

|  | **Cognitive Ability** | **Parental Education** | **Putative Causal** | **GWAS_edu PGS** | **MTAG_10K_eduPGS** | **MTAG_Lead_eduPGS** |
| --- | --- | --- | --- | --- | --- | --- |
| Cognitive Ability | 1<br>(2179) |  |  |  |  |  |
| Parental Education | .256***<br>(2140) | 1<br>(2180) |  |  |  |  |
| Putative Causal_edu PGS | .031<br>(2179) | .025<br>(2180) | 1<br>(2228) |  |  |  |
| GWAS_edu<br>PGS | .044*<br>(2179) | .032<br>(2180) | .134***<br>(2228) | 1<br>(2228) |  |  |
| MTAG_10K_<br>eduPGS | .112***<br>(2179) | .119***<br>(2180) | .227***<br>(2228) | .482***<br>(2228) | 1<br>(2228) |  |
| MTAG_Lead_<br>PGS | .095***<br>(2179) | .117***<br>(2180) | .266***<br>(2228) | .451***<br>(2228) | .800***<br>(2228) | 1<br>(2228) |

Note: \* $p < .05$ , \*\* $p < .01$ , \*\*\* $p < .001$ .

**Table 12.**
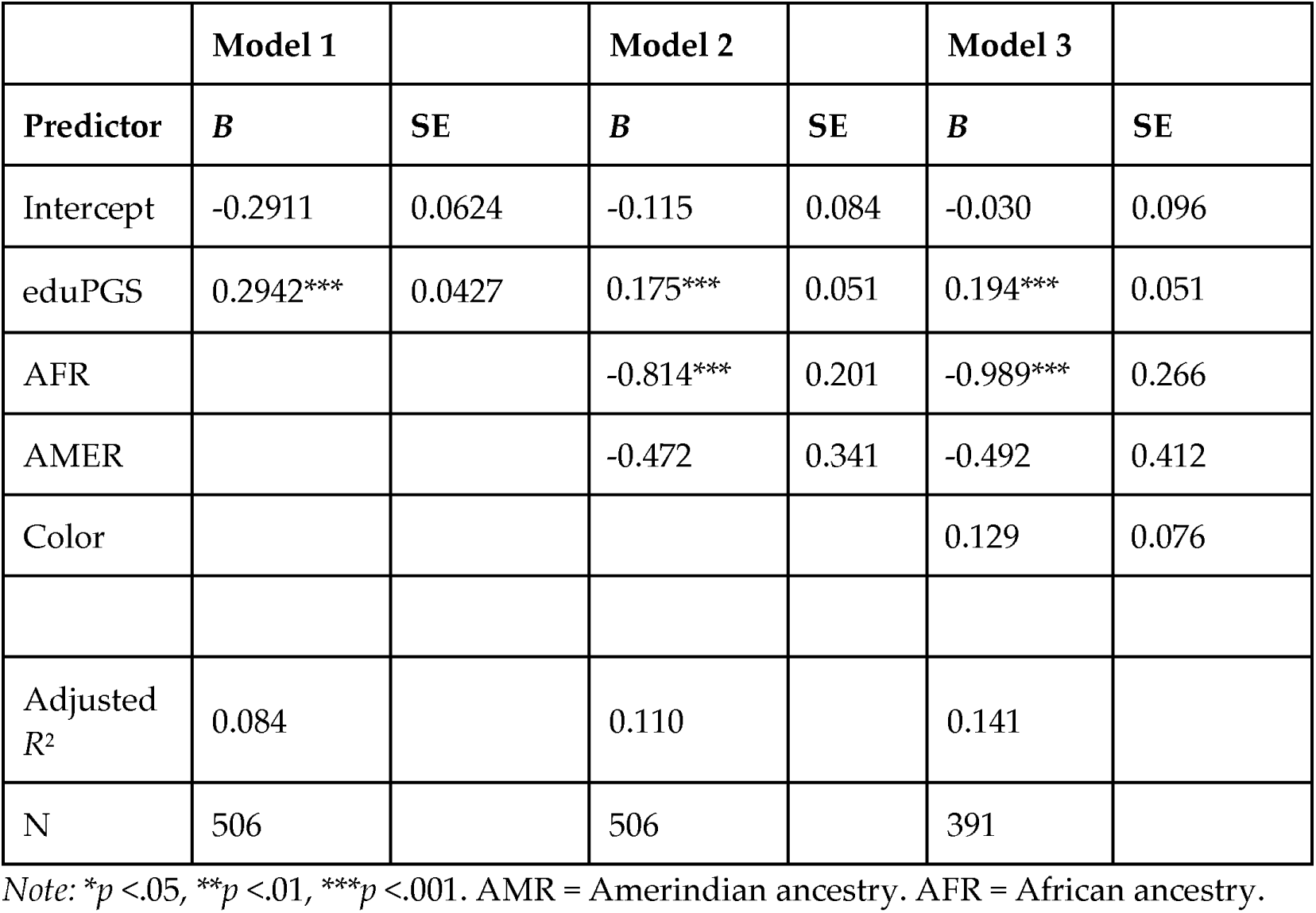
Regression Results for the Effect of eduPGS on Cognitive Ability among Hispanic Americans.

|  | Model 1 |  | Model 2 |  | Model 3 |  |
| --- | --- | --- | --- | --- | --- | --- |
| Predictor | $B$ | SE | $B$ | SE | $B$ | SE |
| Intercept | -0.2911 | 0.0624 | -0.115 | 0.084 | -0.030 | 0.096 |
| eduPGS | 0.2942*** | 0.0427 | 0.175*** | 0.051 | 0.194*** | 0.051 |
| AFR |  |  | -0.814*** | 0.201 | -0.989*** | 0.266 |
| AMER |  |  | -0.472 | 0.341 | -0.492 | 0.412 |
| Color |  |  |  |  | 0.129 | 0.076 |
| Adjusted $R^2$ | 0.084 | | 0.110 | | 0.141 | |
| N | 506 |  | 506 |  | 391 |  |

Note: \* $p < .05$ , \*\* $p < .01$ , \*\*\* $p < .001$ . AMR = Amerindian ancestry. AFR = African ancestry.

We used the MTAG 10k PGS for further analysis. While this variable did not have the highest validity among Hispanics or European-Africans, it had the highest predictive validity among the two largest groups, European and African Americans. Additionally, Lee et al. (2018; Table 4c) shows that MTAG eduPGS has higher predictive validity then GWAS based eduPGS for cognitive ability in two samples (The National Longitudinal Study of Adolescent to Adult and the Health and Health and Retirement Study). The relation between this eduPGS and between cognitive ability for the four major groups is shown in Figure 6. The slopes (B) and intercepts based on the regression equation were: Hispanic (*B* = 0.294, S.E. = 0.043; Intercept = -0.291; N = 506), European (*B* = 0.230, S.E. = 0.014; Intercept = -0.005; N = 4914), European-African (*B* = 0.284, 0.058; Intercept = 0.020; N = 228), and African American (*B* = 0.151, S.E. = 0.029; Intercept = -0.747; N = 2179). The slope for the Hispanic (*t*(5416) = 1.415, N.S.) and the European-African sample (*t*(5,138) = 0.905, N.S.) was not significantly steeper than the European one, though the power to detect a significant difference was relatively low. For the African American sample, the slope was significantly flatter (*t*(7,089) = 2.45, *p* < 0.05). As noted above, a number of factors can contribute to differences in SNP predictivity, including (1) lack of statistical power, (2) allele frequency differences, (3) linkage disequilibrium, or (4) differences in true causal variant effect sizes/directions (Zanetti & Weale, 2018).

Because we saw previously that ancestry is a robust predictor of cognitive ability in the admixed samples, we next ran multivariate regression to test to what extent the association between eduPGS and ability may be due to confounding with either global ancestry or skin color. Note, as pointed out by Lawson et al. (2020), if the admixed population’s ancestral groups have trait means that differ genetically, then correcting PGS for ancestry may downwardly bias the effect sizes. Model 1, Table 12, shows the effect of eduPGS alone for Hispanics. EduPGS is significantly related to *g* (*B* = 0.294, *N* = 506, *p* < 0.001). When adding ancestry covariates in Model 2, the association attenuates but remains significant (*B* = 0.175, *N* = 506, *p* < 0.001). Based on the formula provided by Clogg, Petkova, and Haritou (1995) for comparing Betas in nested models, the *z*-score for the difference was 1.79, which is significant (*p* = .037; one-tailed). Additionally, adding color in Model 3 did not further attenuate this association (*B* = 0.194, *N* = 391, *p* < 0.001). Note, the association in Model 2 for Hispanics was not substantially different from that in the equivalent model for Europeans (Model 2_European_: *B* = 0.230, *N* = 4,914).

Next we ran the same analysis on the two other heavily admixed groups, European-African and African Americans. Table 13, Model 1, shows the effect of eduPGS alone for European-African Americans (*B* = 0.284, *N* = 228, *p* < 0.001). Adding ancestry in Model 2 seems to attenuate the association somewhat but the effect remains significant (*B* = 0.215, *N* = 228, *p* < 0.01). When color was added in Model 3, the association also remained significant (*B* = 0.181, *N* = 164, *p* < 0.05).

**Table 13.**
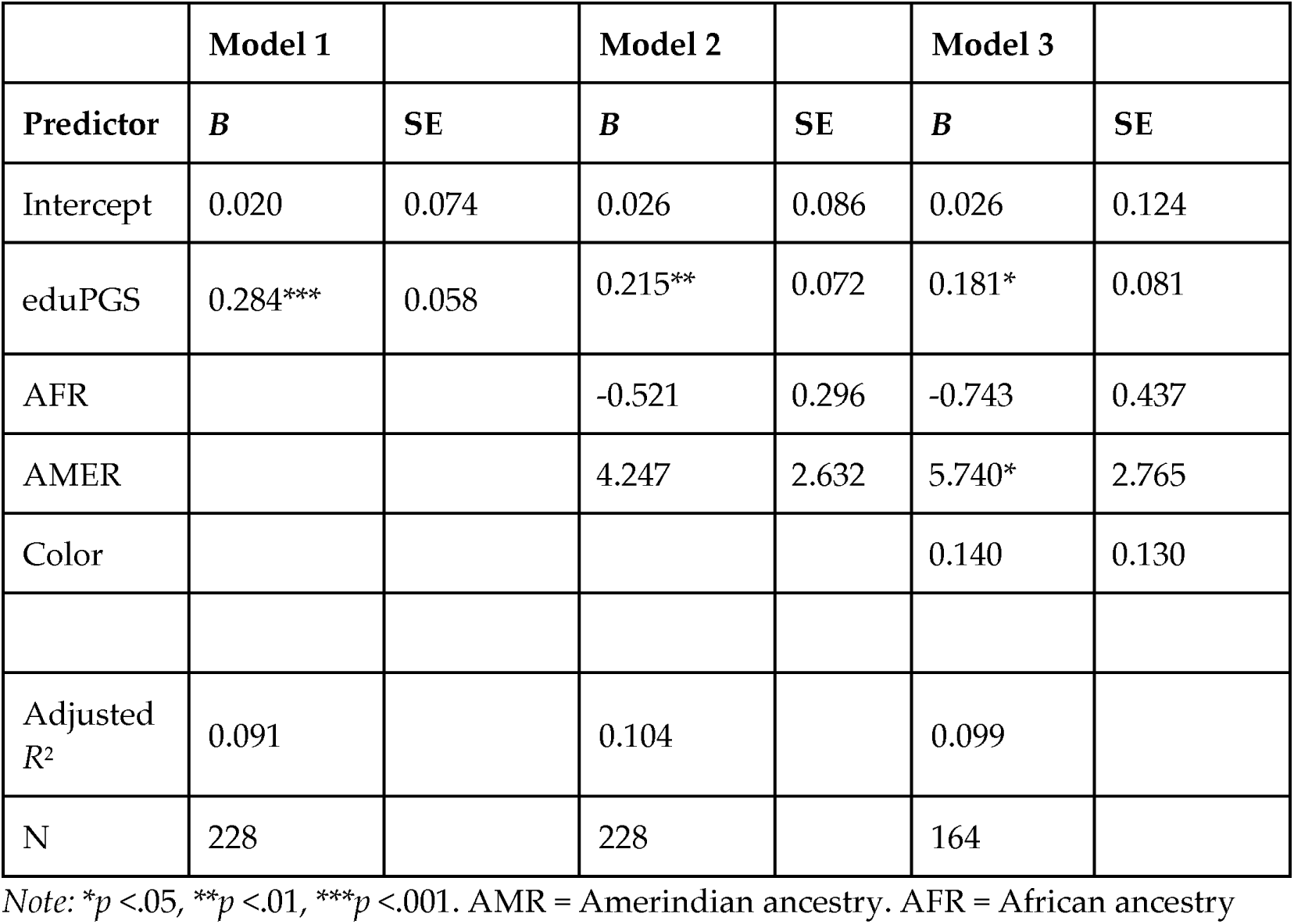
Regression Results for the Effect of eduPGS on Cognitive Ability among European-African Americans.

For African Americans, shown in Table 14, the effect of eduPGS on *g* (*B* = 0.151, *N* = 2,179, *p* < 0.001) again seems to be attenuated somewhat but remains significant (*B* = 0.126, *N* = 2,179, *p* < 0.001) with the inclusion of ancestry in Model 2. As seen in Model 3, adding color did not further attenuate this association (*B* = 0.133, *N* = 1,526, *p* < 0.001).

**Table 14.** Regression Results for the Effect of eduPGS on Cognitive Ability among African Americans.

|  | Model 1 |  | Model 2 |  | Model 3 |  |
| --- | --- | --- | --- | --- | --- | --- |
| Predictor | *B* | SE | *B* | SE | *B* | SE |
| Intercept | -0.747 | 0.055 | -0.423 | 0.171 | -0.229 | 0.204 |
| eduPGS | 0.151*** | 0.029 | 0.126*** | 0.030 | 0.133*** | 0.037 |
| AFR |  |  | -0.478* | 0.219 | -0.532 | 0.278 |
| AMER |  |  | 0.398 | 1.198 | -0.455 | 1.691 |
| Color |  |  |  |  | -0.066 | 0.043 |
| Adjusted $R^2$ | 0.012 | | 0.014 | | 0.021 | |
| N | 2179 |  | 2179 |  | 1526 |  |

Note: \* $p < .05$ , \*\* $p < .01$ , \*\*\* $p < .001$ . AMR = Amerindian ancestry. AFR = African ancestry.

Finally, for the combined sample, Model 1, Table 15, shows the effect of eduPGS alone. EduPGS is significantly related to *g* (*B* = 0.384, *N* = 7920, *p* < 0.001). When adding ancestry covariates in Model 2, the association is attenuated but remains significant (*B* = 0.221, *N* = 7,920, *p* < 0.001). Additionally, adding SIRE and color, in Model 4, did not further substantially attenuate this association (*B* = 0.218, *N* = 5,991, *p* < 0.001).

**Table 15.** Regression Results for the Effect of eduPGS on Cognitive Ability for the Combined Sample.

|  | Model 1 |  | Model 2 |  | Model 3 |  | Model 4 |  |
| --- | --- | --- | --- | --- | --- | --- | --- | --- |
| Predictor | $B$ | SE | $B$ | SE | $B$ | SE | $B$ | SE |
| Intercept | -0.322 | 0.012 | -0.117 | 0.016 | -0.117 | 0.017 | -0.144 | 0.025 |
| eduPGS | 0.384*** | 0.010 | 0.221*** | 0.013 | 0.219*** | 0.013 | 0.218*** | 0.014 |
| AFR |  |  | -0.795*** | 0.042 | -0.511*** | 0.110 | -0.501*** | 0.136 |
| AMER |  |  | -0.251 | 0.205 | 0.214 | 0.238 | 0.430 | 0.289 |
| SIRE_AA |  |  |  |  | -0.249** | 0.087 | -0.224* | 0.097 |
| SIRE EA_AA |  |  |  |  | 0.074 | 0.071 | 0.048 | 0.084 |
| SIRE HI_AA |  |  |  |  | -0.183 | 0.110 | -0.213 | 0.124 |
| SIRE HI_EA |  |  |  |  | -0.192 | 0.098 | -0.180 | 0.117 |
| SIRE HI_OT |  |  |  |  | -0.406* | 0.107 | -.309** | 0.117 |
| SIRE Other |  |  |  |  | -0.081 | 0.074 | -0.048 | 0.083 |
| Color |  |  |  |  |  |  | 0.006 | 0.026 |
| Adjusted R<sup>2</sup> | 0.159 |  | 0.195 |  | 0.196 |  | 0.196 |  |
| N | 7920 |  | 7920 |  | 7920 |  | 5991 |  |

Note: \* $p < .05$ , \*\* $p < .01$ , \*\*\* $p < .001$ . AMR = Amerindian ancestry. AFR = African ancestry.

### 3.5 Path Analysis

While fitting cross-sectional data to a path model cannot prove the causal assumptions, doing so can provide estimates of the effect magnitudes on the assumption that the model is correct (Bollen & Pearl, 2013). As such, we depict two sets of path model results fit with the **lavaan** R package (Rosseel, 2012). We limited the first path analyses to Hispanics to reduce transethnic bias in eduPGS validity. In the first model, we include European ancestry, color, and eduPGS as covariates. As color scores were not missing at random, we did not impute data, but rather handled missing data with listwise deletion. As with previous analyses, European ancestry was left unstandardized. The path model is shown in Figure 10. The path estimates are shown in Table 16. In the model, eduPGS partially explained the association between European ancestry and cognitive ability. Independent of eduPGS, European ancestry was also strongly positively associated with cognitive ability. As expected, European ancestry was strongly negatively associated with darker color. However, darker skin color had no significant independent effect on cognitive ability. Moreover, the sign of the Beta here was in the “wrong” direction relative to predictions from a colorism model. That is, darker color was unexpectedly associated with higher intelligence when controlling for ancestry. The model indicates that eduPGS is a plausible component; color, in contrast, does not seem to be a plausible mediator.

**Figure 10.**
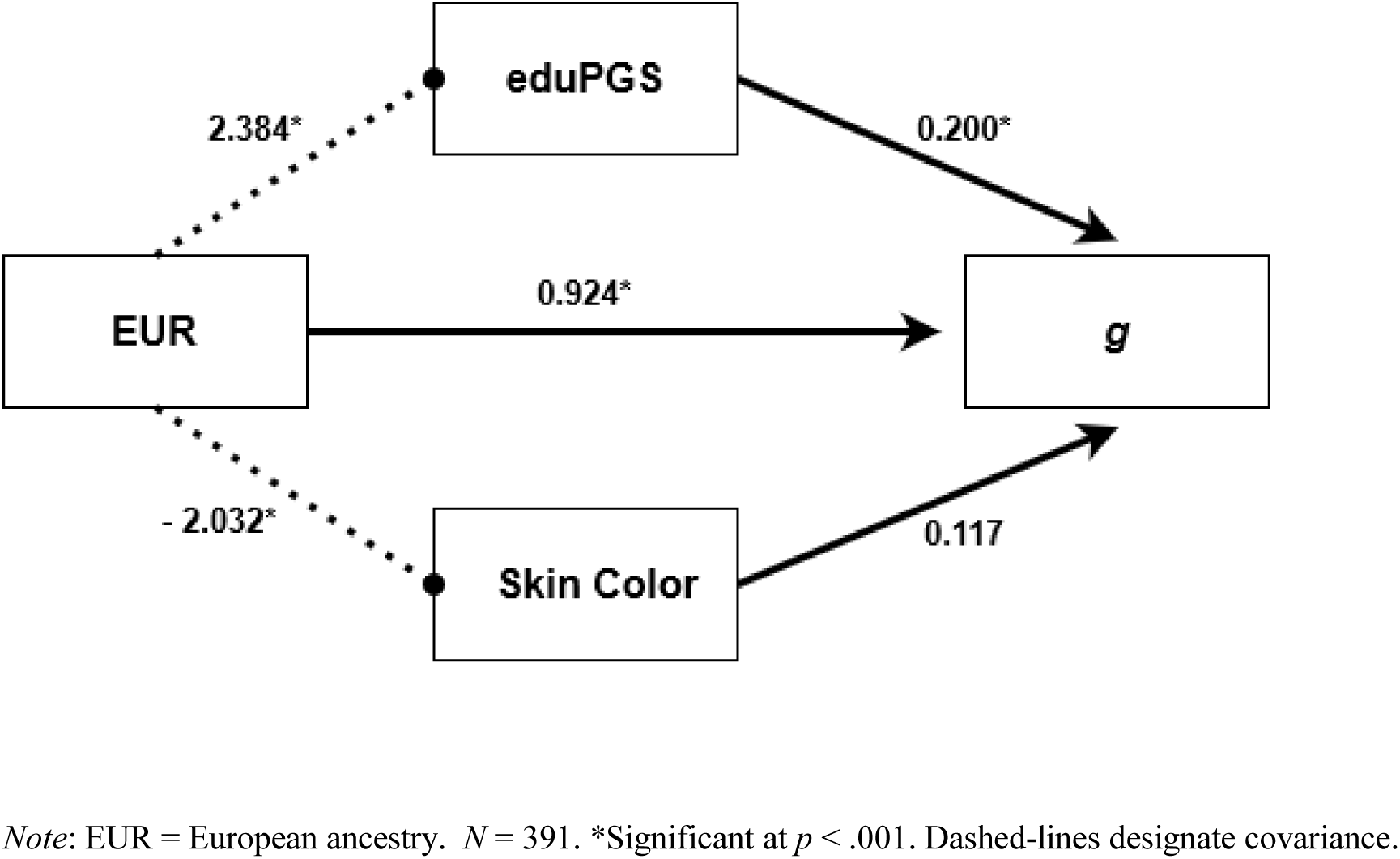
Path Diagram for the Relation between European Ancestry, Color, eduPGS, and g in the Hispanic American Sample.

**Table 16.** Detailed Results for the Path Diagram between European Ancestry, Color, eduPGS, and *g* for Hispanics.

| | | | Unstandardized Estimate | S.E. | $P$ value | Lower 95% CI | Upper 95% CI | Standardized Estimate |
| --- | --- | --- | --- | --- | --- | --- | --- | --- |
| EUR | → | $g$ | 0.924 | 0.260 | 0.000 | 0.414 | 1.433 | 0.256 |
| EUR | → | eduPGS | 2.442 | 0.168 | 0.000 | 2.112 | 2.773 | 0.591 |
| eduPGS | → | $g$ | 0.200 | 0.051 | 0.000 | 0.101 | 0.299 | 0.229 |
| Skin Color | → | $g$ | 0.117 | 0.075 | 0.118 | -0.030 | 0.264 | 0.098 |
| EUR | → | Skin Color | -2.032 | 0.114 | 0.000 | -2.256 | -1.809 | -0.670 |
| Skin Color | ~ | eduPGS | 0.009 | 0.032 | 0.774 | -0.054 | 0.073 | 0.015 |

Note: EUR = European ancestry. Tilde designates covariance.

**Table 17.** Detailed Results for the Path Diagram between European Ancestry, Color, eduPGS, and *g* for the Combined Sample.

|  |  |  | Unstandardized Estimate | S.E. | P value | Lower 95% CI | Upper 95% CI | Standardized Estimate |
| --- | --- | --- | --- | --- | --- | --- | --- | --- |
| EUR | → | $g$ | 0.718 | 0.079 | 0.000 | 0.563 | 0.872 | 0.232 |
| EUR | → | eduPGS | 2.384 | 0.032 | 0.000 | 2.320 | 2.447 | 0.688 |
| eduPGS | → | $g$ | 0.222 | 0.014 | 0.000 | 0.194 | 0.250 | 0.248 |
| Skin Color | → | $g$ | -0.001 | 0.026 | 0.983 | -0.051 | 0.050 | 0.000 |
| EUR | → | Skin Color | -2.404 | 0.018 | 0.000 | -2.440 | -2.369 | -0.865 |
| Skin Color | ~ | eduPGS | -0.008 | 0.006 | 0.176 | -0.019 | 0.004 | -0.017 |

Note: EUR = European ancestry. Tilde designates covariance.

In Figure 11, we repeat this analysis with the complete sample. Since SIRE had little consistent effect, independent of ancestry, we do not include SIRE variables in the path analysis. The results are comparable, except that the Beta for color is *B* = -.001 instead of *B* = .117 (both nonsignificant).

**Figure 11.**
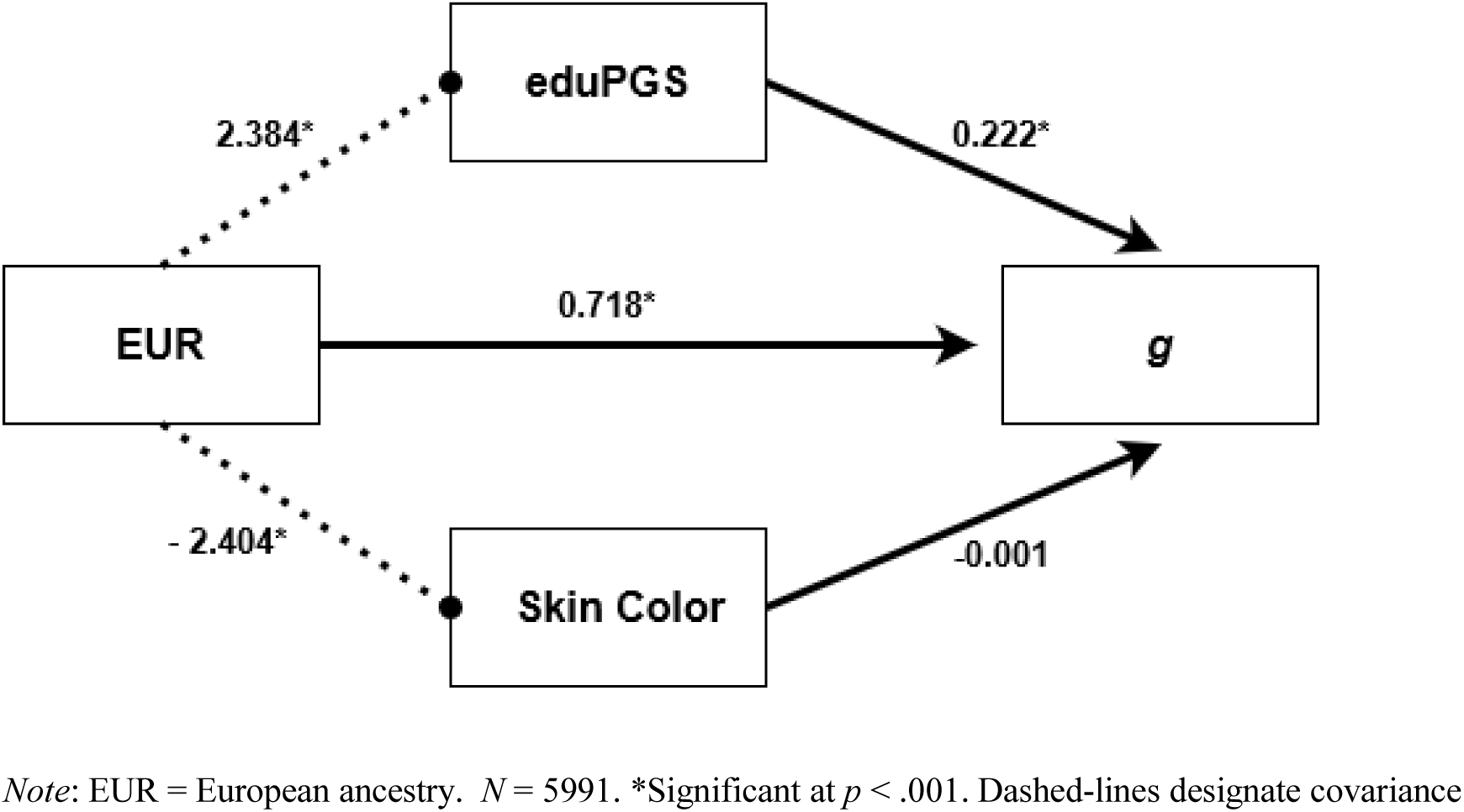
Path Diagram for the Relation between European Ancestry, Color, eduPGS, and g in the Complete Sample.

As an alternative model, we include parental education instead of color. Color was dropped as it was not a significant predictor of *g* and because there were limited cases with color scores. While most parents are likely biological parents, not all are, and so the adolescent’s ancestry is not exactly equivalent to the midparent ancestry in this case. As such, we represent the relationship between the adolescent’s European ancestry and their parent’s education as covariance (indicated by dashed lines). For Hispanics, the path model is shown in Figure 12, and the estimates appear in Table 18. In the model, eduPGS again partially explained the association between European ancestry and *g*. However, parental education was also a significant predictor. The covariance between parental education and adolescent eduPGS was significant. With the current data, however, it is not possible to disentangle the causal paths between European ancestry, parental education, adolescent eduPGS, and adolescent *g*. This is because adolescent eduPGS approximates biological mid-parent eduPGS. And it is expected that, within populations, mid-parent eduPGS will be causally related to parental educational levels, which, in turn, will be genetically correlated with adolescent *g* (see, e.g., Trzaskowski et al., 2014). Nonetheless, this model also indicates that eduPGS plausibly has constitutive explanatory relevance regarding the relation between European ancestry and *g*.

**Figure 12.**
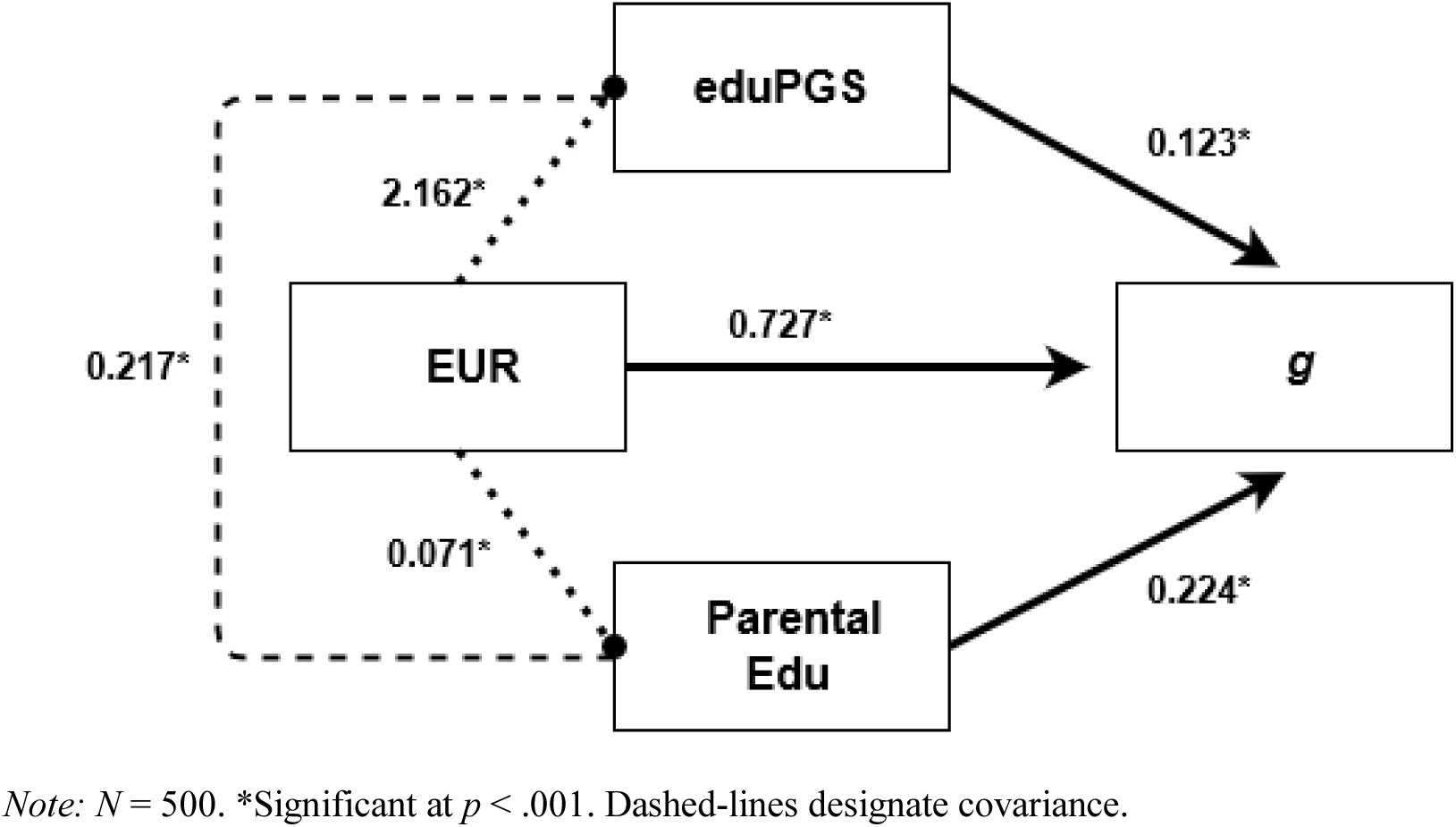
Path Diagram for the Relation between European Ancestry, eduPGS, Parental Education, and g in the Hispanic American Sample.

**Table 18.**
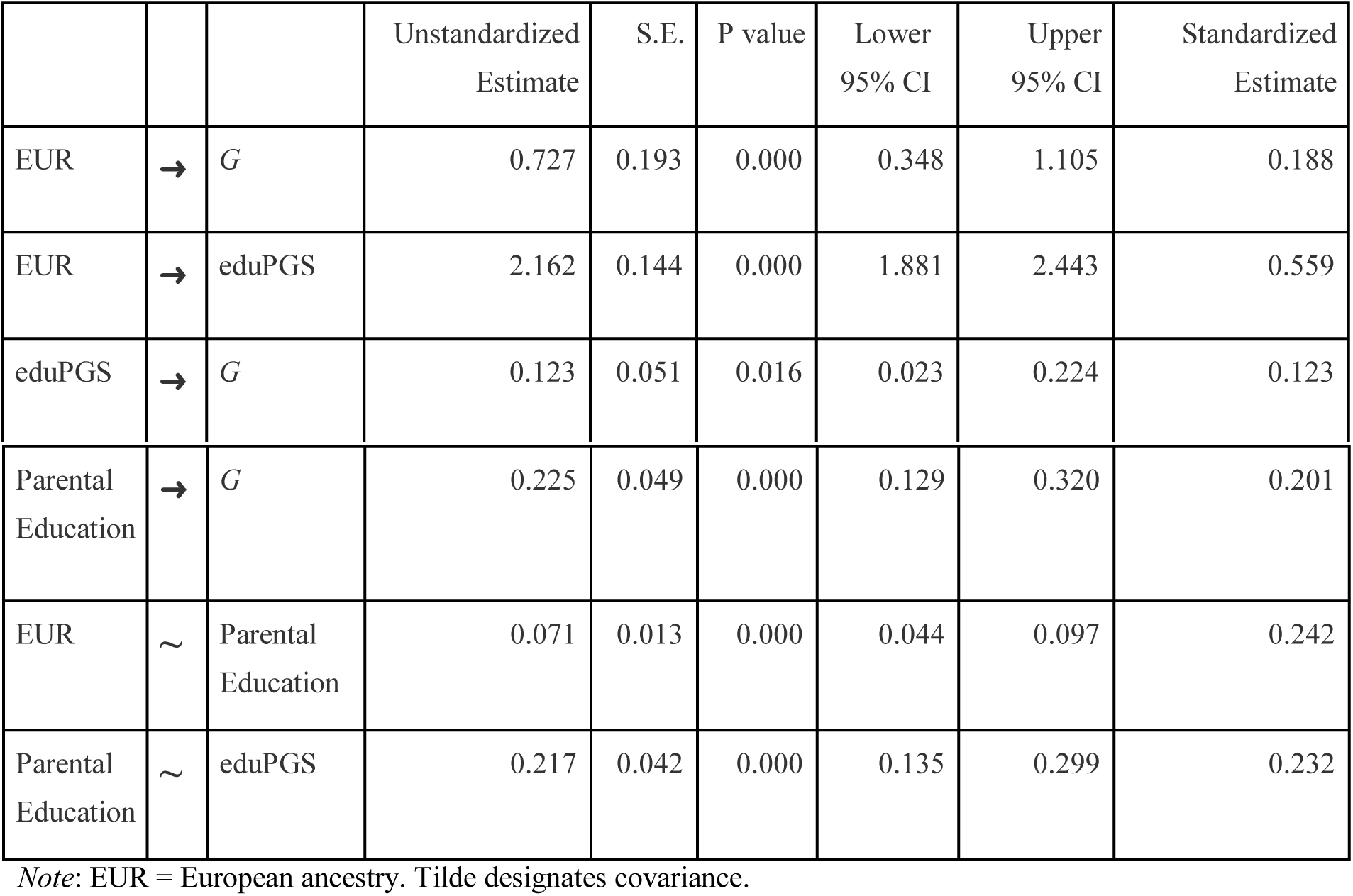
Detailed Results for Path Diagram with Parental Education.

|  |  |  | Unstandardized Estimate | S.E. | P value | Lower 95% CI | Upper 95% CI | Standardized Estimate |
| --- | --- | --- | --- | --- | --- | --- | --- | --- |
| EUR | → | $G$ | 0.727 | 0.193 | 0.000 | 0.348 | 1.105 | 0.188 |
| EUR | → | eduPGS | 2.162 | 0.144 | 0.000 | 1.881 | 2.443 | 0.559 |
| eduPGS | → | $G$ | 0.123 | 0.051 | 0.016 | 0.023 | 0.224 | 0.123 |
| Parental Education | → | *G* | 0.225 | 0.049 | 0.000 | 0.129 | 0.320 | 0.201 |
| EUR | ~ | Parental Education | 0.071 | 0.013 | 0.000 | 0.044 | 0.097 | 0.242 |
| Parental Education | ~ | eduPGS | 0.217 | 0.042 | 0.000 | 0.135 | 0.299 | 0.232 |

Note: EUR = European ancestry. Tilde designates covariance.

Again, we repeat this analysis with the combined sample. The path model is shown in Figure 13, and the estimates appear in Table 19. The results are comparable, except that the Beta for color is *B* = -.001 instead of *B* = .117 (both nonsignificant).

**Figure 13.**
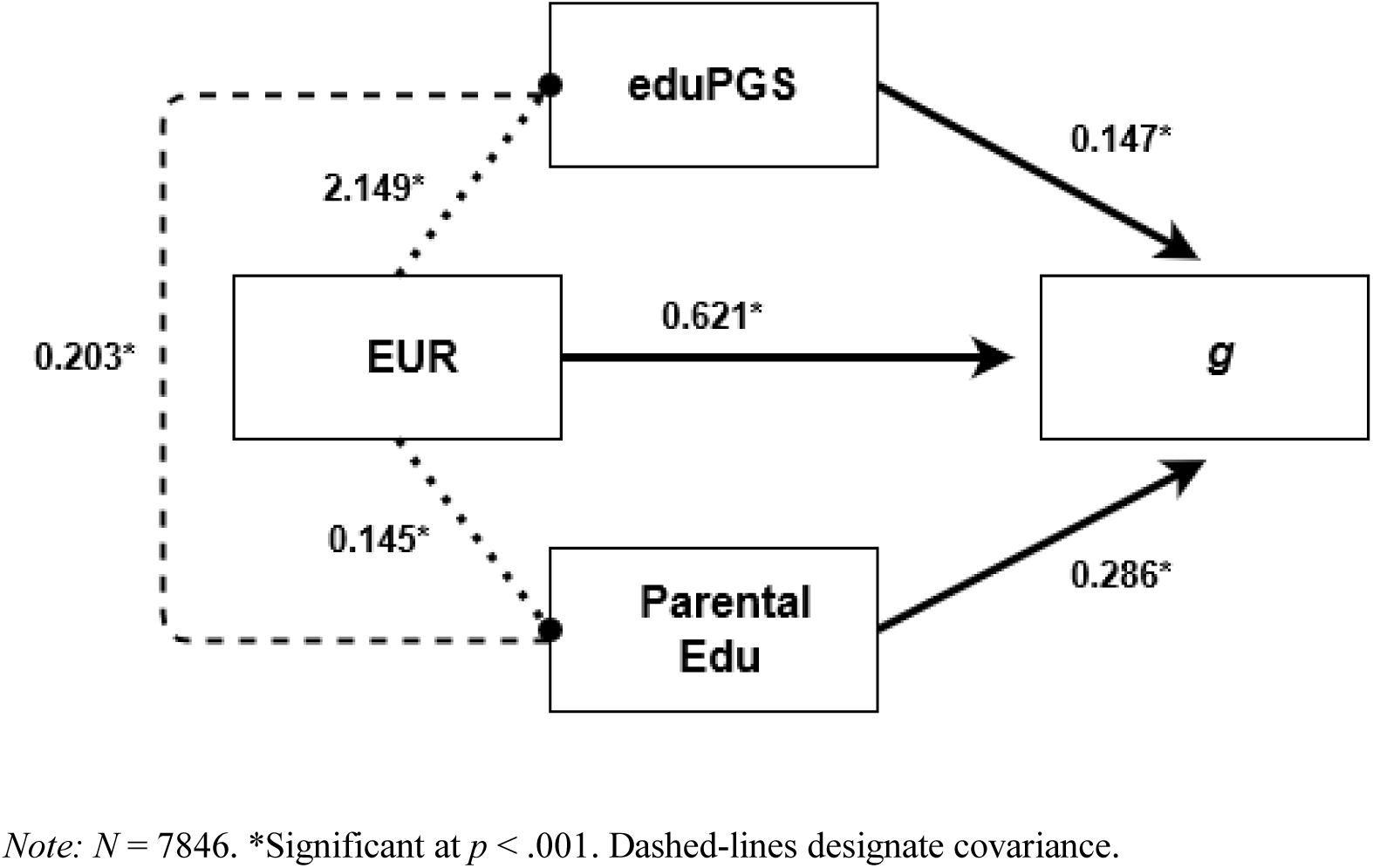
Path Diagram for the Relation between European Ancestry, eduPGS, Parental Education, and g in the Combined Sample.

**Table 19.**
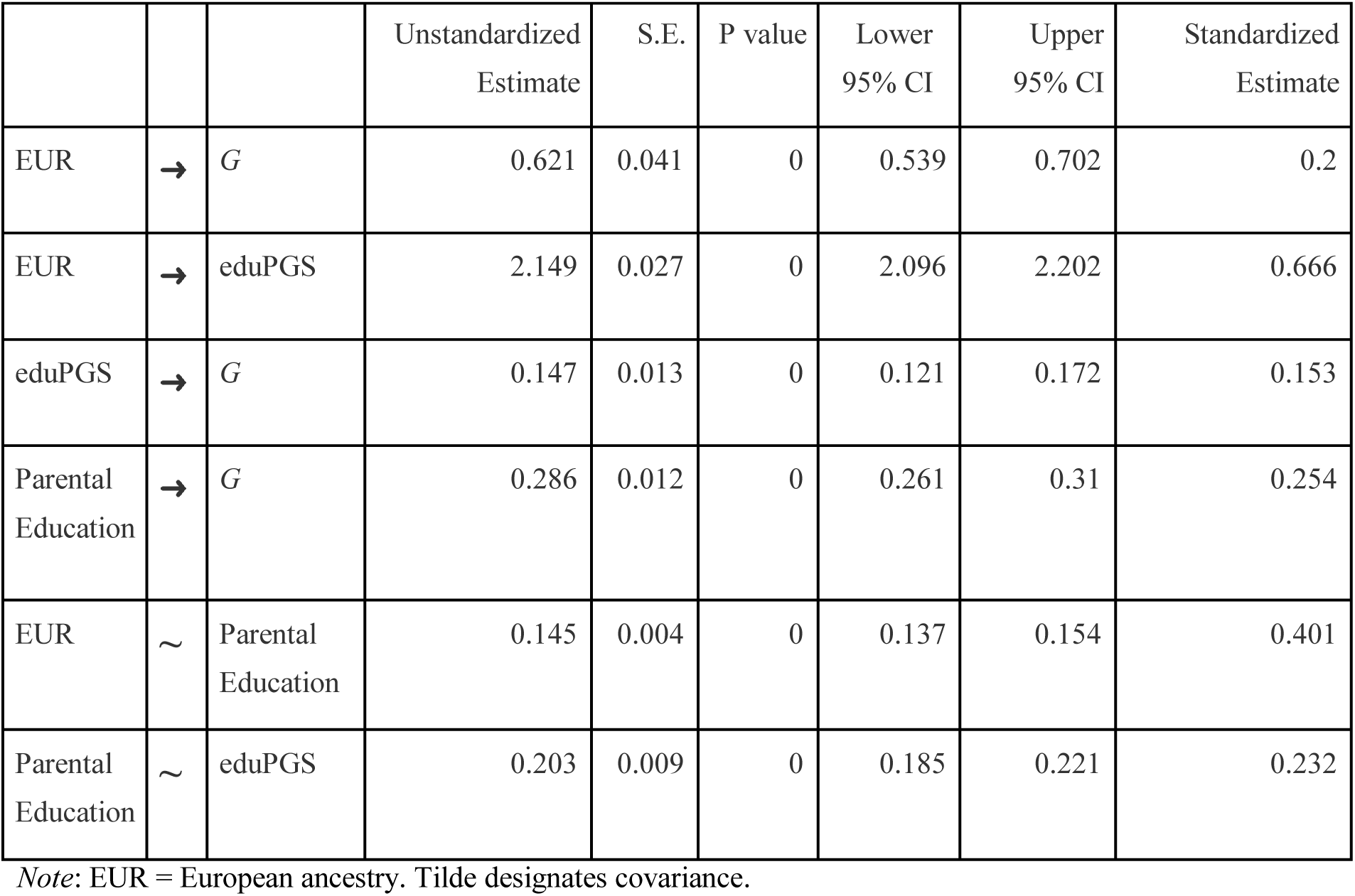
Detailed Results for Path Diagram with Parental Education.

|  |  |  | Unstandardized Estimate | S.E. | P value | Lower 95% CI | Upper 95% CI | Standardized Estimate |
| --- | --- | --- | --- | --- | --- | --- | --- | --- |
| EUR | → | *G* | 0.621 | 0.041 | 0 | 0.539 | 0.702 | 0.2 |
| EUR | → | eduPGS | 2.149 | 0.027 | 0 | 2.096 | 2.202 | 0.666 |
| eduPGS | → | *G* | 0.147 | 0.013 | 0 | 0.121 | 0.172 | 0.153 |
| Parental Education | → | *G* | 0.286 | 0.012 | 0 | 0.261 | 0.31 | 0.254 |
| EUR | ~ | Parental Education | 0.145 | 0.004 | 0 | 0.137 | 0.154 | 0.401 |
| Parental Education | ~ | eduPGS | 0.203 | 0.009 | 0 | 0.185 | 0.221 | 0.232 |

Note: EUR = European ancestry. Tilde designates covariance.

### 3.6 The Spearman-Jensen Hypothesis

If the association between eduPGS and cognitive ability is primarily genetic in nature, a Jensen effect between eduPGS effects and subtest *g*-loadings is expected, since genetic effects tend to primarily act through *g* (te Nijenhuis et al., 2019). Moreover, since the weak version of Spearman’s hypothesis fits the data well, for both the African/European and Hispanic/European differences, it is expected that the magnitude of the differences on subtests will positively correlate with the subtest *g*-loadings (i.e., there will be a Jensen Effect on score differences). Spearman’s hypothesis can also be extended to associations with genetic ancestry. If the association between ancestry (not just SIRE) and cognitive ability is primarily due to ancestry-related differences in *g*, a Jensen effect with respect to ancestry would also be expected. The relevance of determining whether this is the case has been previously discussed (Hu et al., 2019). The vectors of group differences (EA / E_AA, EA / AA, and EA / HI) are based on the means and standard deviations reported in Table 2. The other vectors are reported in Table 20.

**Table 20.**
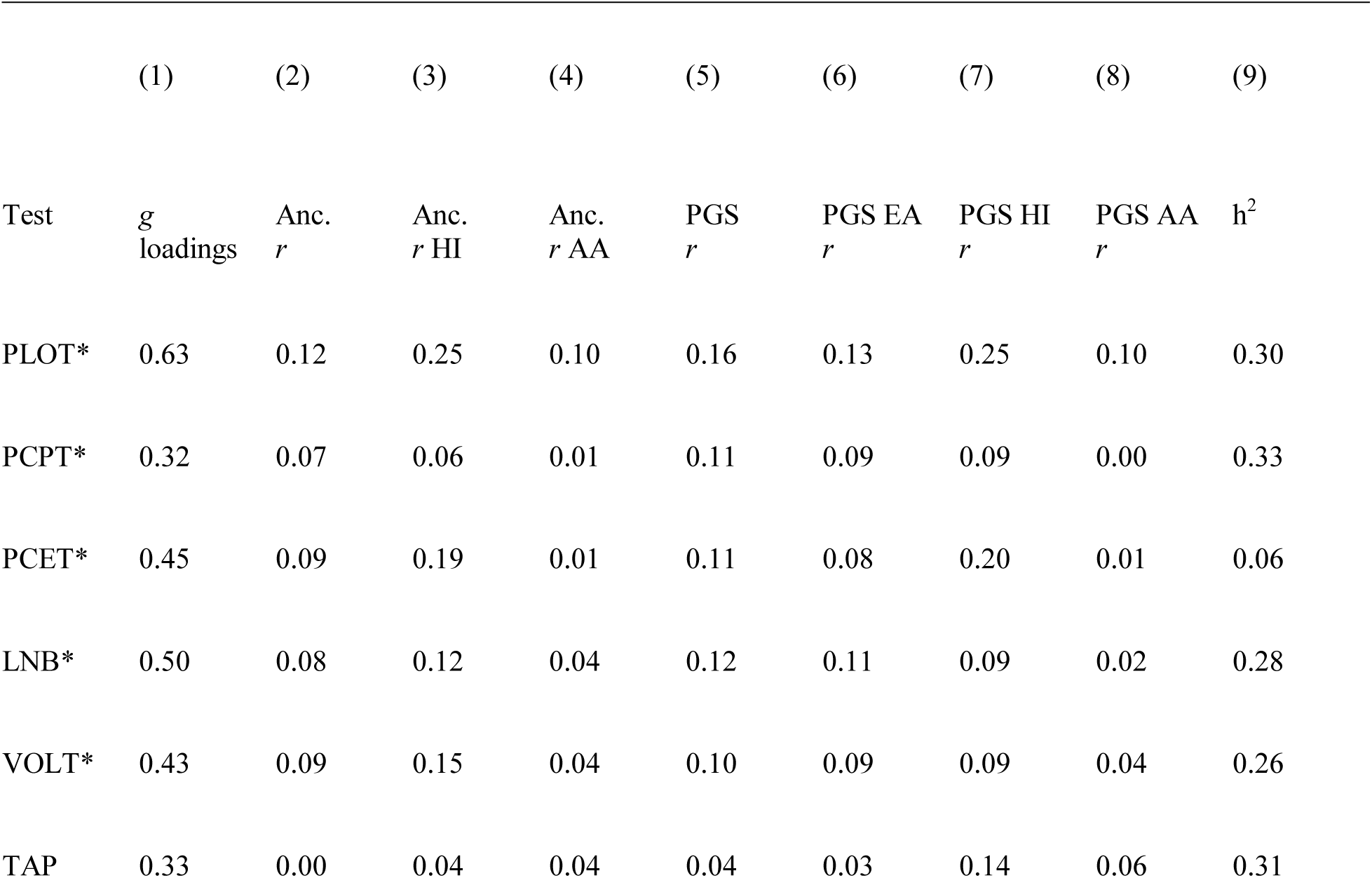
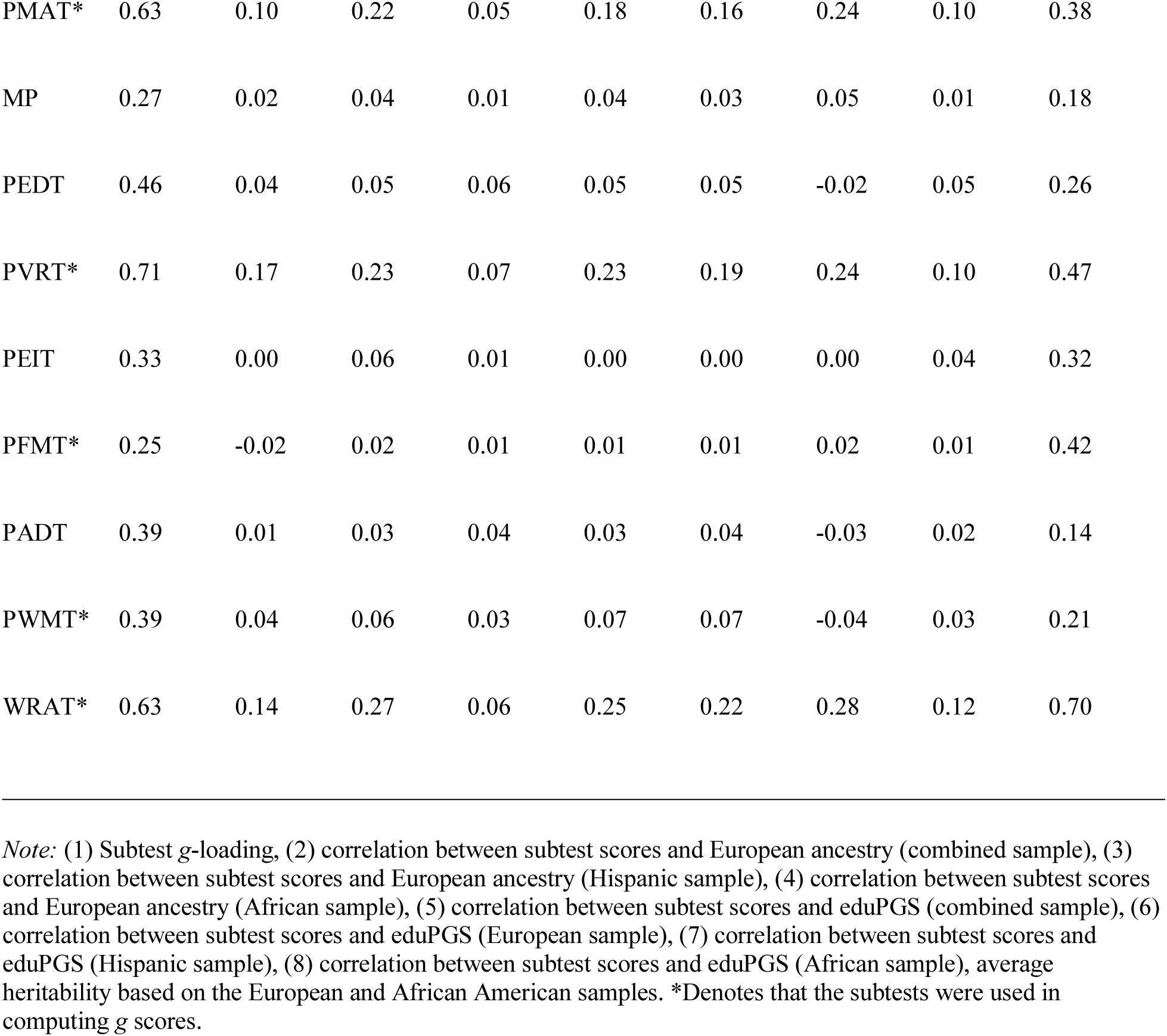
Rounded Vectors for the Method of Correlated Vector Analysis.

|  | (1) | (2) | (3) | (4) | (5) | (6) | (7) | (8) | (9) |
| --- | --- | --- | --- | --- | --- | --- | --- | --- | --- |
| Test | $g$<br>loadings | Anc.<br>$r$ | Anc.<br>$r$ HI | Anc.<br>$r$ AA | PGS<br>$r$ | PGS EA<br>$r$ | PGS HI<br>$r$ | PGS AA<br>$r$ | $h^2$ |
| PLOT* | 0.63 | 0.12 | 0.25 | 0.10 | 0.16 | 0.13 | 0.25 | 0.10 | 0.30 |
| PCPT* | 0.32 | 0.07 | 0.06 | 0.01 | 0.11 | 0.09 | 0.09 | 0.00 | 0.33 |
| PCET* | 0.45 | 0.09 | 0.19 | 0.01 | 0.11 | 0.08 | 0.20 | 0.01 | 0.06 |
| LNB* | 0.50 | 0.08 | 0.12 | 0.04 | 0.12 | 0.11 | 0.09 | 0.02 | 0.28 |
| VOLT* | 0.43 | 0.09 | 0.15 | 0.04 | 0.10 | 0.09 | 0.09 | 0.04 | 0.26 |
| TAP | 0.33 | 0.00 | 0.04 | 0.04 | 0.04 | 0.03 | 0.14 | 0.06 | 0.31 |
| PMAT* | 0.63 | 0.10 | 0.22 | 0.05 | 0.18 | 0.16 | 0.24 | 0.10 | 0.38 |
| MP | 0.27 | 0.02 | 0.04 | 0.01 | 0.04 | 0.03 | 0.05 | 0.01 | 0.18 |
| PEDT | 0.46 | 0.04 | 0.05 | 0.06 | 0.05 | 0.05 | -0.02 | 0.05 | 0.26 |
| PVRT* | 0.71 | 0.17 | 0.23 | 0.07 | 0.23 | 0.19 | 0.24 | 0.10 | 0.47 |
| PEIT | 0.33 | 0.00 | 0.06 | 0.01 | 0.00 | 0.00 | 0.00 | 0.04 | 0.32 |
| PFMT* | 0.25 | -0.02 | 0.02 | 0.01 | 0.01 | 0.01 | 0.02 | 0.01 | 0.42 |
| PADT | 0.39 | 0.01 | 0.03 | 0.04 | 0.03 | 0.04 | -0.03 | 0.02 | 0.14 |
| PWMT* | 0.39 | 0.04 | 0.06 | 0.03 | 0.07 | 0.07 | -0.04 | 0.03 | 0.21 |
| WRAT* | 0.63 | 0.14 | 0.27 | 0.06 | 0.25 | 0.22 | 0.28 | 0.12 | 0.70 |

*Note:* (1) Subtest *g*-loading, (2) correlation between subtest scores and European ancestry (combined sample), (3) correlation between subtest scores and European ancestry (Hispanic sample), (4) correlation between subtest scores and European ancestry (African sample), (5) correlation between subtest scores and eduPGS (combined sample), (6) correlation between subtest scores and eduPGS (European sample), (7) correlation between subtest scores and eduPGS (Hispanic sample), (8) correlation between subtest scores and eduPGS (African sample), average heritability based on the European and African American samples. \*Denotes that the subtests were used in computing *g* scores.

Table 21 shows the vector correlations (using unrounded vectors). The results based on the ten subtest MI model appear above the diagonal; while, those for all 15 subtests appear below. As can be seen, all associations are strongly positive. The strong association between *g*-loadings and eduPGS is consistent with the finding that Lee et al.’s (2018) eduPGS is associated with genetic *g* (de la Fuente, 2020). Consistent with other research, there is a strong Jensen effect on ethnic differences (te Nijenhuis, van den Hoek, & Dragt, 2019) and on ancestry-related differences within ethnic groups (Hu et al., 2019; Lasker et al., 2019). Generally, the effect of eduPGS, like that of ancestry and SIRE group differences, is pronounced on the most *g*-loaded and more heritable subtests.

**Table 21.**
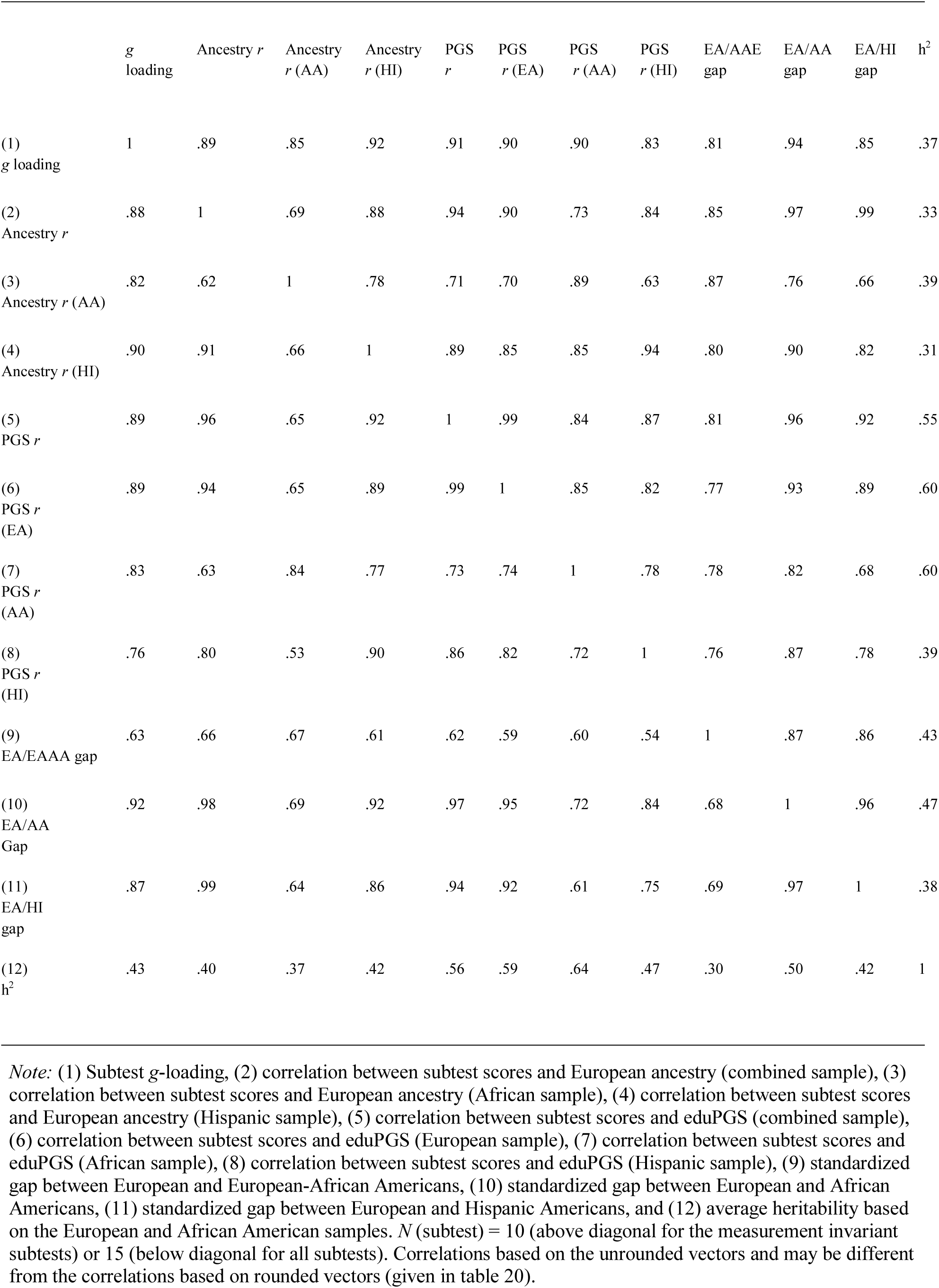
Results from Method of Correlated Vectors Analysis.

## 4. Evaluation of Bias in Education-related PGS (eduPGS)

### 4.1. Evaluation of Bias in the TCP Sample

PGS may be biased due to the source population with which they were computed (i.e., ascertainment bias). There are a couple of obvious mechanisms by which this bias can occur. First, European-based eduPGS may be biased against non-Europeans due to the inclusion of European-specific variants (Thomson, 2019). These population-specific variants might have very low frequencies in non-European populations. Second, out-of-African based eduPGS may be biased against African populations due to an overrepresentation of derived (due to new mutations) versus ancestral (shared with other primates) variants in the out-of-African populations (Kim et al., 2018; Thomson, 2019). As a robustness check, we investigate both possibilities.

First, we computed MTAG eduPGS excluding variants with minor allele frequency (MAF) < 0.01 (leaving 7,636 overlapping variants) and <0.05 (leaving 7,172 overlapping variants) among African lineages, using the 1000 Genomes reference samples to determine the African MAF. As a result, these eduPGS exclude variants not also present in African populations. As seen in Table 22, this exclusion had no substantive effect on the mean eduPGS differences between groups.

**Table 22.**
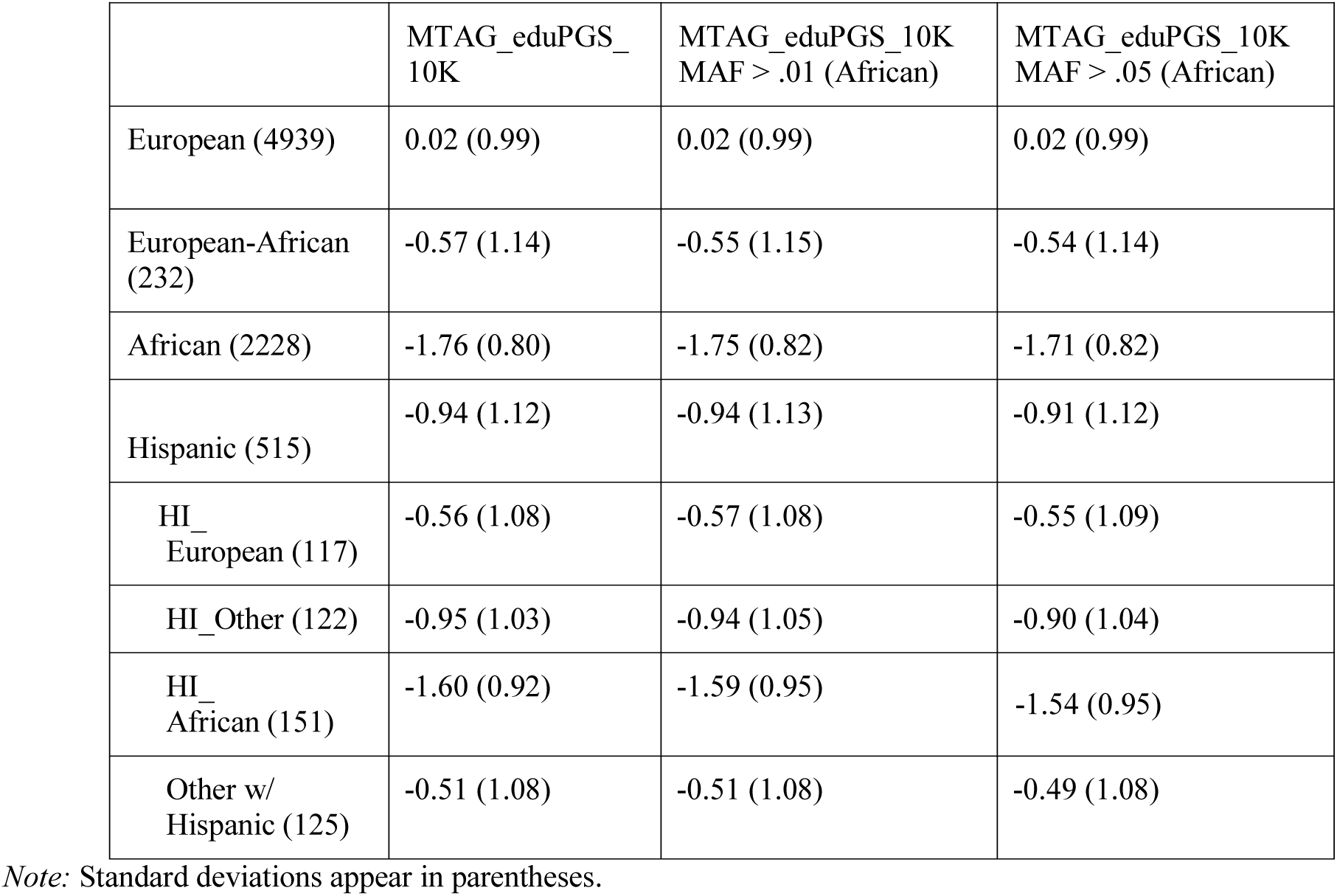
EduPGS scores for European, Hispanic, and African American Participants.

|  | MTAG_eduPGS_10K | MTAG_eduPGS_10K<br>MAF > .01 (African) | MTAG_eduPGS_10K<br>MAF > .05 (African) |
| --- | --- | --- | --- |
| European (4939) | 0.02 (0.99) | 0.02 (0.99) | 0.02 (0.99) |
| European-African (232) | -0.57 (1.14) | -0.55 (1.15) | -0.54 (1.14) |
| African (2228) | -1.76 (0.80) | -1.75 (0.82) | -1.71 (0.82) |
| Hispanic (515) | -0.94 (1.12) | -0.94 (1.13) | -0.91 (1.12) |
| HI_ European (117) | -0.56 (1.08) | -0.57 (1.08) | -0.55 (1.09) |
| HI_Other (122) | -0.95 (1.03) | -0.94 (1.05) | -0.90 (1.04) |
| HI_ African (151) | -1.60 (0.92) | -1.59 (0.95) | -1.54 (0.95) |
| Other w/ Hispanic (125) | -0.51 (1.08) | -0.51 (1.08) | -0.49 (1.08) |

*Note:* Standard deviations appear in parentheses.

Kim et al. (2018) demonstrated that when allelic risk scores are based on out-of-African populations, African populations show elevated frequencies of disease-associated loci for ancestral (shared with other primates) alleles, and reduced frequencies for derived (due to new mutations after the split with primates) alleles, even when there are no underlying trait differences. They conclude that “systematic allele frequency differences between populations need not be due to any underlying difference in risk” (*p*. 5) and propose corrections for bias due to ancestral versus derived allele status. It has been argued that the eduPGS differences may also be due to a similar form of ascertainment bias (Thomson, 2019). To investigate this, we compute eduPGS by derived and ancestral status. To be clear, we computed one PGS with only those SNPs where the enhancing allele is derived, and then another PGS with only those SNPs where the enhancing allele is ancestral. In this case, risk alleles and trait-enhancing alleles are the same thing; medical versus cognitive GWAS studies just use different terminology. The results are shown in Table 23. As seen, contrary to the findings of Kim et al. (2018), with eduPGS, non-European populations have both lower derived and ancestral eduPGS scores and, moreover, the differences are largest for the derived ones. As a result, when Kim et al.’s (2018, *p*.12) correction is applied, the polygenic score gaps change little (compare Table 22 and Table 23). Thus, this form of ascertainment bias does not explain the eduPGS differences.

**Table 23.**
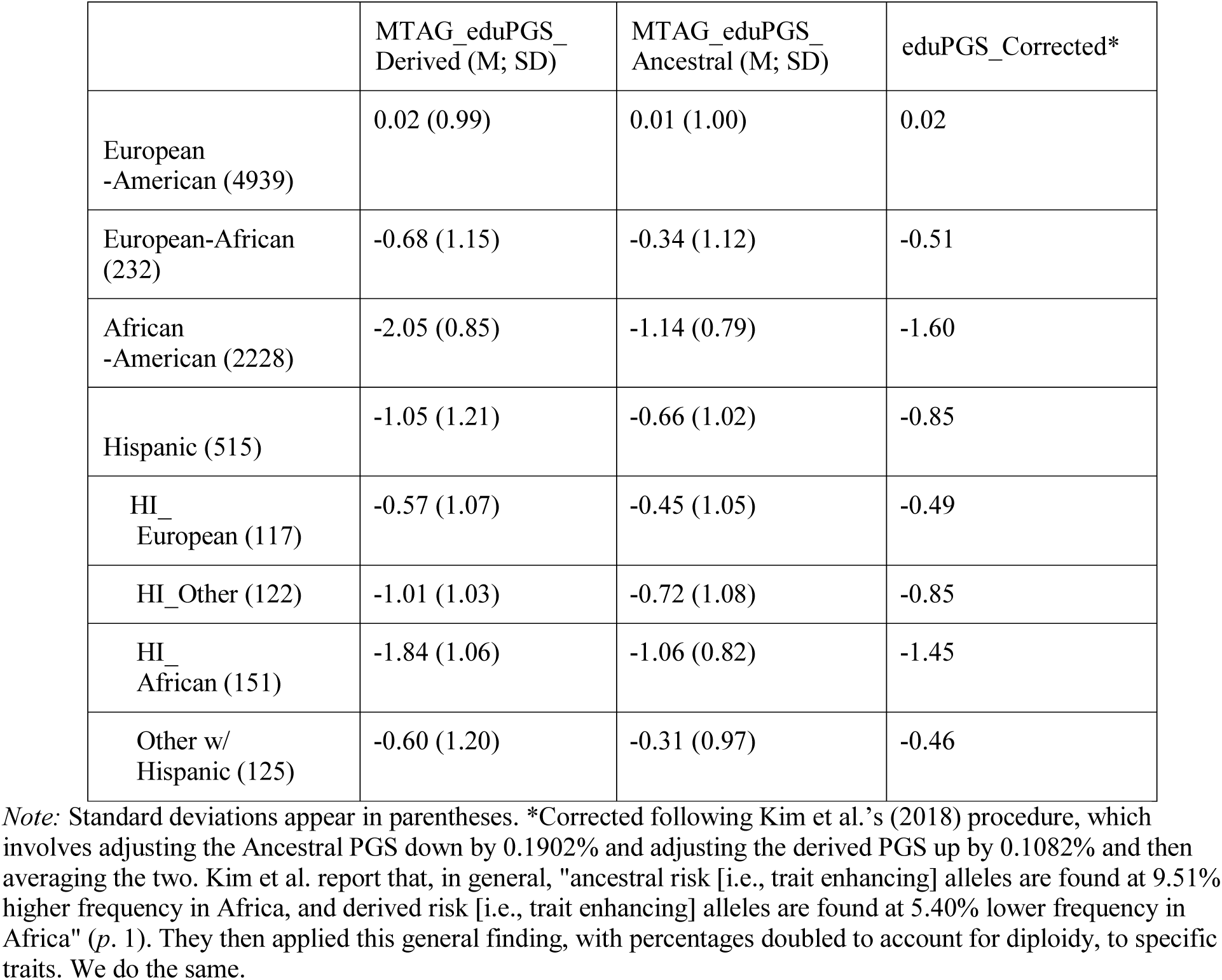
EduPGS scores for European, European-African, African, and Hispanic American Participants.

Additionally, we computed the validities to see if the corrections affected these. The validities by eduPGS are reported in Table 24. As seen, the validities for the different eduPGS were approximately the same for a given SIRE group.

**Table 24.**
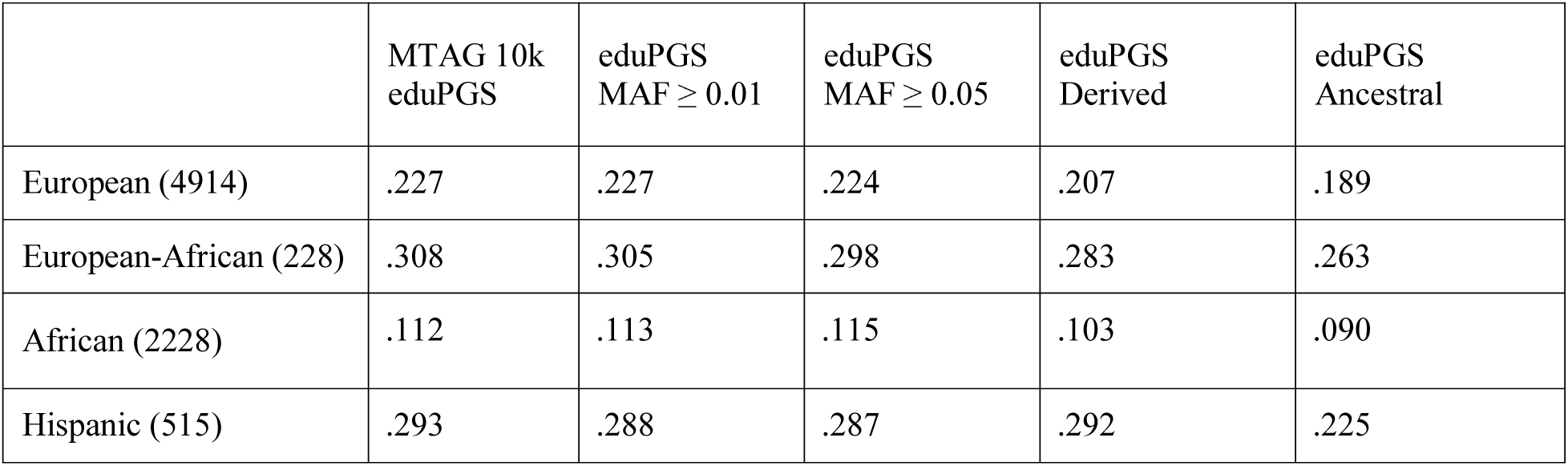
Validities of MAF ≥ 0.01, MAF ≥ 0.05, Derived, Ancestral eduPGS for Predicting among European, European-African, Hispanic, African and Hispanic Americans.

| | MTAG 10k<br>eduPGS | eduPGS<br>$MAF \geq 0.01$ | eduPGS<br>$MAF \geq 0.05$ | eduPGS<br>Derived | eduPGS<br>Ancestral |
| --- | --- | --- | --- | --- | --- |
| European (4914) | .227 | .227 | .224 | .207 | .189 |
| European-African (228) | .308 | .305 | .298 | .283 | .263 |
| African (2228) | .112 | .113 | .115 | .103 | .090 |
| Hispanic (515) | .293 | .288 | .287 | .292 | .225 |

Finally, it has been argued that the eduPGS differences may be upwardly biased by using the discovery sample-based direction of effects for SNPs (Thompson, 2019). To investigate this we computed the Betas for the MTAG SNPs, used as predictors of *g*, for European (*N*= 4,939) and African Americans (*N*= 2,228) separately. The SIRE specific Betas are provided in the Supplemental Material. We then examined mean differences and validities for the variants which showed transracially concordant effects across ethnic groups as compared to those which showed discordant effects. African Americans had lower eduPGS than EA based on the concordant SNPs (-3.13 and 0.00, respectively), but slightly higher eduPGS based on the discordant SNPs (0.49 and 0.00, respectively). Moreover, among African Americans, the concordant SNPs showed higher validity for *g* than they did for the discordant SNPs. Cross validation confirmed that the concordant eduPGS were more predictive than the discordant eduPGS among African Americans. A possible interpretation is that the discordant eduPGS, which are less likely to be causal, contain more LD decay related effects. Regardless, there is no evidence, based on this sample, that the eduPGS differences are being inflated by the inclusion of SNPs with transethnically discordant effects as some have argued (Thomson, 2019). However, this issue will have to be revisited when SNP effect sizes based on larger non-European samples are available.

### 4.2. Evaluation of Bias in 1000 Genomes Samples

Since ascertainment bias and confounding related to population stratification is of significant concern (Thompson, 2019), we ran supplementary analyses which leveraged the 1000 Genomes data to explore the effects of score construction on the magnitudes of population differences. Because the main ancestral groups of European-Africans, Africans and Hispanics in the current sample are Africans and Europeans, we restricted focus to individuals of Northern and Central European descent from Utah (CEU) and Yoruba Nigerians (YRI) from the 1000 Genomes Project. Methods and detailed results are reported in Supplemental File 2.

4.2.1. *Population-GWAS vs. Within family weights*

A reader suggested computing eduPGS using Lee et al.’s (2018) within-family effect sizes. These weights were based on a GWAS for education using 22,000 sibling pairs (as compared to the 1.1 million individuals for the regular estimates). As Lee et al. (2018) note, these within-family estimates are smaller than the corresponding estimates from the population GWAS. The authors explore different reasons for this (Suppl. Note *pp*. 21-38) and reason that the lower validity is likely due in part to a within-family reduction of gene-by-environmental correlation. This conclusion is consistent with results from a number of other studies (e.g., Liu et al., 2018; Selzam, Ritchie, Pingault, Reynolds, O’Reilly, & Plomin, 2019).

In context to this analysis, the rationale for using within-family Betas is that they are more robust to confounding from structure. Using within-family Betas does not completely address the problem of population stratification, since there could be bias due to SNP selection. For example, Zaidi and Mathieson (2020) found that "[w]hile sibling-based association tests are immune to stratification, the hybrid approach of ascertaining variants in a standard GWAS and then re-estimating effect sizes in siblings reduces but does not eliminate bias" (abstract).

To examine the impact of using population-GWAS versus within family Beta weights, we created eduPGS for both CEU and YRI individuals using the regular MTAG Betas, which we will refer to as population-GWAS Betas, and the within-family Betas. To do this, we used the MTAG-based SNPs as prior. The SNPs were filtered for MAF >0.01 for both CEU and YRI. These were then weighted by the population-GWAS Betas reported by Lee et al. (2018) and the within family weights provided by the authors upon request. The standardized scores, decomposed also by ancestral and derived status, are shown in Table 25. Based on a Welch Two Sample *t*-test, these differences were all highly statistically significant. Finally, to address the concern raised by Zaidi and Mathieson (2020), we computed the differences using the 4,413 within-family SNPs that had a *p*-value < .05 along with the within family weights. These thus are pure within-family based eduPGS and so should show no population structure related bias. The s for β CEU and YRI were 0.529 and -.485, respectively with a β difference of 1.01. Results are shown in Figure S3.

**Table 25.**
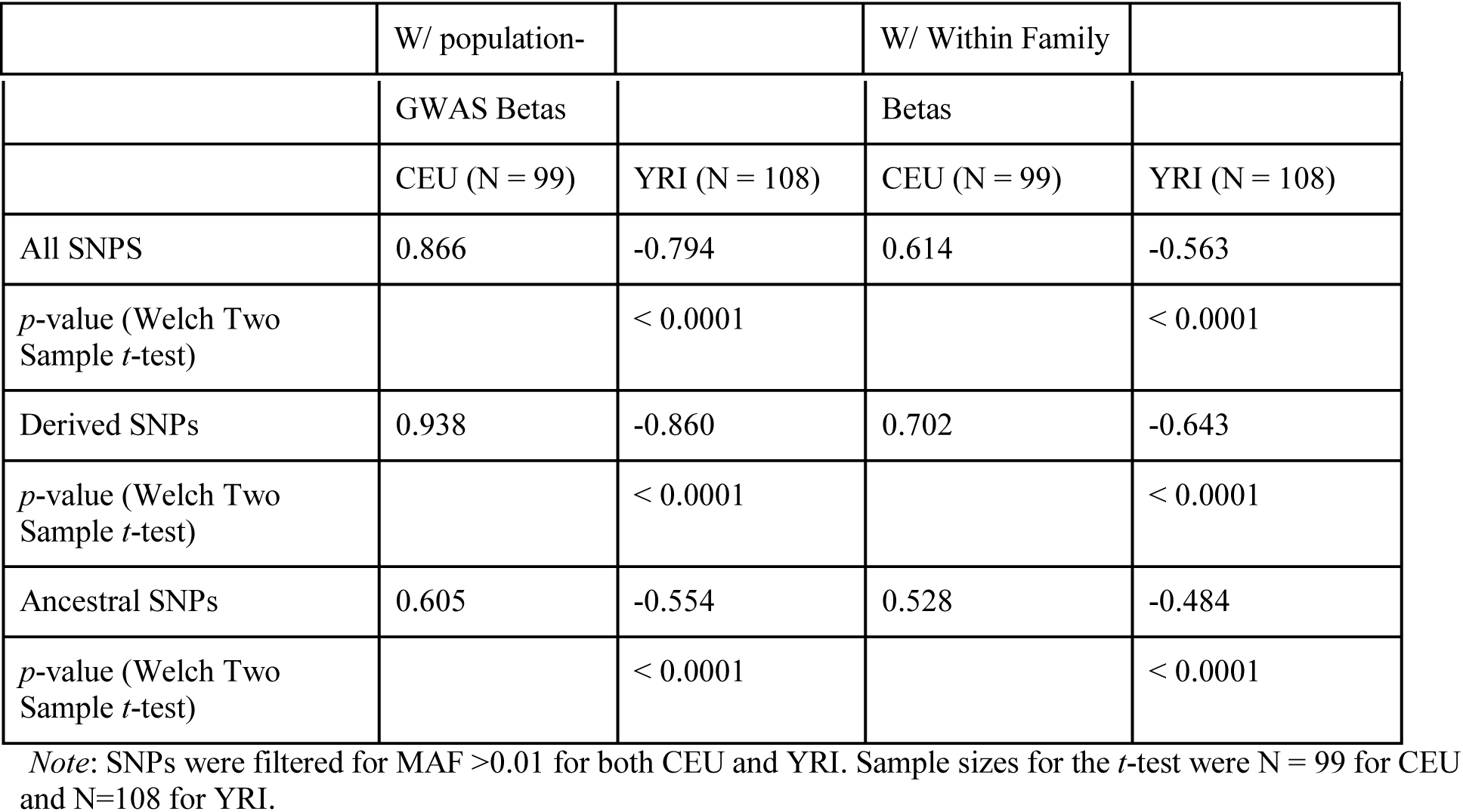
Mean MTAG-based PGS for CEU and YRI Calculated using population-GWAS and Within Family Betas.

#### 4.2.2. Trans-Ethnically Concordant Betas

Previous polygenic selection studies have applied European GWAS βs to different world populations (e.g., Berg and Coop, 2014). In theory, however, taking into account information about population-specific effects (i.e., βs for both European and non-European comparison β samples) should yield more accurate results both on the individual and population levels (Grinde et al., 2018). Indeed, the PGS βs for SNPs in European samples will often show opposite or β discordant effects in non-European samples. It has been argued that including SNPs with transracially discordant effects may bias the group differences and so that “the polygenic scores should be computed only from those GWAS hits that have directionally consistent effects in the races that are being compared” (Thompson, 2019). Thus, for this analysis, we use the two largest TCP samples, European and African Americans, to classify MTAG βs into trans-ethnically concordant and discordant ones. We then recomputed concordant and discordant eduPGS and compared the magnitude of the 1000 Genomes CEU and YRI differences. We computed eduPGS for the concordant and discordant SNPs separately, using weights from Lee et al. (2018). More detail on the methods is provided in Supplementary File 2.

As shown in Table 26 and Table 27, the differences were largest in the trans-ethnically concordant SNPs. In line with the results from 4.1, the discordant PGS showed no CEU-YRI difference (95% C.I. = -0.009, 0.005), while the concordant CEU-YRI difference was around 3% (95% C.I. = 0.025, 0.039). Thus, the presence of discordant variants may possibly be masking CEU-YRI differences as would be the case if individual differences were caused by variants common across ethnic groups and, also, if the transethnic differences were larger for causally-relevant SNPs. However, again, this issue will have to be reevaluated when SNP effect sizes based on larger non-European samples are available. At present, we can only say that given the data available, the eduPGS differences are not inflated as a result of inclusion of SNPs with transethnically discordant effects.

**Table 26.** Concordant, Discordant, and “Naive” PGS Frequencies by Population

| Population | PGS 10k MTAG | PGS concordant | PGS discordant |
| --- | --- | --- | --- |
| CEU | 0.5063 | 0.5050 | 0.5093 |
| YRI | 0.4904 | 0.4726 | 0.5109 |
| Difference (CEU - YRI) | 0.0159 | 0.0325 | -0.0016 |

**Table 27.** . Results of *t*-test for PGS Frequency CEU-YRI difference

|  | *T* | *P*-value | 95% CI for Mean Difference | Df |
| --- | --- | --- | --- | --- |
| PGS naïve | 6.288 | $3.37 \times 10^{-10}$ | 0.0109, 0.0208 | 8418 |
| PGS concordant | 9.013 | $2.2 \times 10^{-16}$ | 0.0254, 0.0395 | 4305 |
| PGS discordant | -0.453 | 0.651 | -0.0088, 0.0055 | 3993 |

*Note:* Sample sizes for the *t*-test were the number of concordant and discordant SNPs. Using the number of individuals, instead, did not change the interpretation of the results.

#### 4.2.3. Cross-population Validity

Cross-population analyses are frequently used to assess the validity of PGS (e.g., Berg et al., 2019, Figure 1; Sohail et al., 2019; Figure 4). Results based on height and other PGS have led some to conclude that PGS are meaningless for cross-population comparisons. However, whether this is so needs to be evaluated on a case by case basis. Thus, in the third analysis, we compared the cross-population predictive validity, for measured population cognitive ability, of the concordant, discordant, 10K MTAG, and MTAG-lead PGS. Further, we examine if eduPGS computed from Lee et al.’s (2018) sibling analysis data predicts national cognitive scores. While sibling analyses are robust to population structure-related confounding, none of Lee et al.’s within-family SNPs met the minimum for GWAS significance (5e-8) and so the eduPGS computed from them provides a very noisy signal. Methods and additional results are detailed in Supplementary File 2.

We were able to construct eduPGS for 18 countries based on the 1000 Genomes data. For the 16 countries with data, Lynn and Becker’s (2019) national IQs correlated with World Bank’s (2017) cognitive test scores at *r* = .86 (*N* = 16). Since there were more cases for the former (18), we report those results here. The highest predictive validity (*r* = .82) was attained by the eduPGS based on MTAG-lead SNPs, followed by eduPGS based on all MTAG SNPs (*r* = .81). The correlation between concordant and discordant 10K MTAG eduPGS and measured population IQ were *r* = .79 and *r* = .21, respectively; this difference was statistically significant (*t* = 5.95, *p* < 0.01), with, as expected, the concordant eduPGS being more predictively valid. The within-family based eduPGS also had a validity (*r* = .474) which, while statistically significant, was significantly lower than the 10k MTAG eduPGS one (*t* = 3.785 *p* < 0.01).

In sum, MTAG-based eduPGS, except the trans-ethnically discordant one, is predictive on the population level. Generally, while it has been found that, in the case of some other PGS, “differences in polygenic risk scores across populations are significant but not supported by epidemiological or anthropometric studies of the same traits” (Martin et al., 2017, *p*. 644) this appears to not be the case for MTAG-based eduPGS. Nonetheless, results based on the within-family based eduPGS highlights Duncan et al.’s (2019) caution that different eduPGS can give markedly different results, rendering interpretation uncertain.

## 5. Discussion

### 5.1 **General Discussion**

To better understand how eduPGS function in admixed American populations, and to determine if there is a robust confound vs. causal problem, we examined the association between intelligence-related polygenic scores, global ancestry, and general cognitive ability in East Coast Hispanic and non-Hispanic European, European-African, and African American samples. Prior to analysis, we conducted MGCFA to assure that measurement invariance held between the main ethnic groups. We were able to confirm full factorial invariance in the case of the European-Hispanic and European-African differences. Moreover, the weak form of Spearman’s hypothesis seemed to be the best fitting model in context to both the European / African and European / Hispanic differences.

Among Hispanics and in the combined Hispanic and non-Hispanic American sample, the association between European/African ancestry and cognitive ability was robust to controls for SIRE, color, and parental education. For Amerindian ancestry, in both the Hispanic-only and the combined samples, the association with cognitive ability became nonsignificant when SIRE was added as a covariate. In the Hispanic-only sample, however, the effect remained directionally consistent with recently reported results (Kirkegaard et al. 2019; Warne, 2020). Statistically, the insignificance, in this case, was a result of the higher standard errors of Amerindian ancestry, as compared to African, which was due to the reduced variance in Amerindian ancestry. Similarly, in a small, “non-Hispanic Caucasian” sample, with little variance in ancestry, European ancestry versus (apparently) Mexican/Mexican-American ancestry was only weakly and non-significantly associated with IQ (*r* = .09, *N* =120; Wang, Pandika, Chassin, Lee & King, 2016).

For the combined sample, however, the effect of Amerindian ancestry turned positive with SIRE controls. Statistically this was due to Amerindian ancestry being slightly positively correlated with general intelligence in the non-Hispanic White sample (*r* = .014; *N* = 4914, *N.S*.) and to the much larger non-Hispanic White than Hispanic sample size. However, in this case, the Amerindian ancestry may have been of northern extraction and thus not directly comparable with that in the other samples. Moreover, the amount and variance of Amerindian ancestry among Philidephian non-Hispanic Whites (*M* = 0.01%; *SD* = 0.04%) was marginal. Generally, the relation between cognitive ability and Amerindian ancestry needs to be better explored in predominantly European-Amerindian “Mestizo” samples. The results to date are ambiguous.

Many older studies have shown that the social and phenotypic indexes of admixture are associated with cognitive ability in admixed American samples. This is the case for Latin Americans in the United States and throughout Central and South America, in addition to Native American and Afro-descent groups in English speaking America (e.g., Paschal & Sullivan, 1925; Vincenty, 1930; Curti, 1960; Grinder, Spotts, & Curti, 1964; Green, 1972; Kirkegaard & Fuerst, 2017; Hailu, 2018). Additionally, a meta-analysis of epidemiological studies also showed that European ancestry was positively (and African and Amerindian negatively) associated with education and economic indices throughout Latin and Anglo-American admixed populations (Kirkegaard, Wang, & Fuerst, 2017). Considering these results, the finding here and elsewhere that European genetic ancestry positively predicts cognitive ability among Latin Americans should not be completely surprising. On the other hand, genetic ancestry was previously reported to not be significantly associated with IQ in both Latin and Anglo American countries, based on older studies using blood-groups (e.g., Rothhammer & Llop, 1976; Loehlin, Vandenberg, & Osborne, 1973; Scarr, Pakstis, Katz, & Barker, 1977). These results are regularly cited as (often “direct”) evidence against non-environmental models for group differences by several authors (e.g., Templeton, 2001; Nisbett, 2009; Mackintosh, 2011; Nisbett et al., 2012; Lilienfeld et al., 2014; Coleman, 2016; Halpern & Kanaya, 2019). Thus, while not completely surprising, the results are nonetheless informative and not self-explanatory.

Here, explanations for the association between ancestry and cognitive ability by confounding with either geography or racial phenotype (see, e.g., Conley & Fletcher, 2017) are not viable. This was a local population sample from the Philadelphia area, so geographic confounding is not a substantial concern. We did find a significant negative association between cognitive ability and darker skin color. However, rather than the association between ancestry and cognitive ability being explained by color, the association between color and cognitive ability was statistically explained by ancestry. These results are key in that there is a large literature on “colorism” which purports to demonstrate color or pigment-based discrimination by showing mere correlations between color and social outcomes (e.g., Marira and Mitra, 2013). Moreover, it has been argued by some that such discrimination might account for potential associations between ancestry and cognitive ability (e.g., Conley & Fletcher, 2017). However, our results concur with the competing distributional model described by Hu et al. (2019, Figure 1), in which the association between color and cognitive ability is a proxy (versus a cause) of that between ancestry and cognitive ability. This finding has implications for genetic research (Lawson et al., 2020), as it suggests that color-based discrimination is not likely an additional source of confounding.

We found that eduPGS was significantly associated with cognitive ability within the European, European-African, African, and Hispanic samples. For the MTAG 10k eduPGS, which we use for further analyses, the associations in the admixed populations were attenuated with the inclusion of ancestry. Regardless, the association remained significant and not substantially different from that in the European sample (*B*_European_ = .230 vs. *B*_European-African_ = .215 and *B*_Hispanic_ = .175), except in the case of African Americans (*B*_African_ = .126), where, while significant, the association was less than that for European Americans. Moreover, path analysis indicated that eduPGS partially statistically explained the association between European ancestry and cognitive ability, whereas color did not (note, among Hispanics, darker color was unexpectedly positively, though insignificantly, associated with *g* in the path model, though this finding did not replicate among the much larger combined sample). Additionally, the explanatory power of eduPGS was not fully accounted for by parental education, a variable which is both genetically and environmentally correlated with adolescent intelligence within groups and thus may be likewise between groups.

The eduPGS differences may be spurious owing to difficult to control ascertainment bias and confounding related to population stratification. For example, Berg et al. (2019) found attenuated effects for height PGS when applying UK versus pan-European-based PGS to Eurasian samples. The reason seems to be that controls for ancestry components do not always fully capture population structure effects, however, some confounding effects can be avoided by computing PGS using a more homogenous population and then applying these to the more heterogeneous populations of interest (see: Berg and Coop, 2014). In our case, though, we start with eduPGS based on European origin samples and then apply these to samples of different continental ancestry. As such, we already incorporated this component of Berg et al.’s (2019) analysis into ours. We further investigated whether differences might be due to biasing effects of discovery population-specific variants, whether the allele associated with higher values of the trait is derived or ancestral, failure of SNP sign concordance between populations, and discrepancy between population-GWAS and within-family coefficients. The forms of ascertainment bias and confounding related to population stratification addressed by our procedures do not explain the group differences in eduPGS. However, it is beyond the scope of this paper to explore all forms of possible confounding. Indeed the point of the paper is not to resolve the issue but to illustrate how ancestry, eduPGS, and *g* are statistically tangled, a situation which will require further research.

The regression and path analysis results raise some possible concerns regarding the use and interpretation of eduPGS in admixed American populations. Ancestry covaries with trait and eduPGS scores. This could either be due to trait-relevant genetic differences between the ancestral groups of the admixed populations or to a mix of both environmental differences and also ascertainment bias and confounding related to population stratification. If the former, controlling for ancestry can attenuate the effect of eduPGS; however, if the later, leaving ancestry unadjusted can inflate it. Thus, whether controlling for ancestry in this context is appropriate depends on the magnitude of the quantitative genetic variance in the trait between ancestral groups (Lawson et al., 2020). Generally, our results are consistent with either a confounding or causal model (depicted in Figure 1).

The differences in eduPGS between ancestral groups are large enough that this is an issue which future research should try to resolve. While the magnitudes of differences for the true causal SNPs are unknown, magnitudes can be calculated for presently known educational related SNPs. We can use the fixation index, a measure of population differentiation, to do this. Supplementary File 3 shows the Fst for the SNPs from Lee et al. (2018) for 1000 Genomes super-populations. The Fst value between Europeans and Africans, which are the two main ancestral groups for the admixed populations here, is .1090 As detailed in Supplementary File 3, for typically reported heritabilities (*h*^2^ = .5; Polderman et al., 2015; Pesta et al., 2020) this magnitude of population differentiation gives medium to large expected mean differences (i.e., when environments are equal), in this case, equivalent to *d* = 0.68. Thus, if transethnically unbiased eduPGS differences turn out to be commensurate with those based on Lee et al.’s (2018) eduPGS, medium to large phenotypic differences are expected under conditions of environmental equality. Controlling for ancestry might then bias eduPGS effects.

Related to this point, a reader suggested that we should run analyses to detect polygenic selection as done by Berg et al. (2019). However, whether differences are due to drift or selection is not necessarily relevant to whether there are genetic differences between populations. Moreover, given the expected differences discussed above, selection, in the form of stabilizing or convergent selection *between populations* is needed to show trait equality, not trait differences. That is, the evolutionary default or null expectation would be that trait-causing SNP frequency differences will be commensurate with random SNP ones, not that selection acted to homogenize differences in this particular trait (Edelaar & Björklund, 2011; Leinonen, McCairns, O’hara, & Merilä, 2013; Edge & Rosenberg, 2015; Rosenberg, Edge, Pritchard, & Feldman, 2019). As Rosenberg et al. note: “[P]henotypic differences among populations are predicted under neutrality to be similar in magnitude to typical genetic differences among populations” (*p*. 30). We are not aware of anyone who has established selection acting between populations such to make causally-relevant genetic differences (related to education and intelligence) substantially smaller than expected by drift. Whether there was divergent selection is undetermined (e.g., Guo et al., 2018; Racimo, Berg, & Pickrell, 2018).

Finally, MCV indicated that the effect of eduPGS was strongly *g*-loaded, as was the effect of ancestry, and group differences. This is consistent with all these effects acting primarily by way of *g*. These results for eduPGS are not self-evident. This is because it has been shown that some of the effect of eduPGS is a shared environmental effect (Domingue & Fletcher, 2019); however, the effect of adoption, a shared environmental effect, exhibits an anti-Jensen effect (te Nijenhuis, Jongeneel-Grimen, & Armstrong, 2015). Thus, that eduPGS would act like typical genetic effects, and be most pronounced on *g*-loaded subtests, is not obvious. In addition to eduPGS, both ancestry and group differences exhibited Jensen Effects, which is consistent with differences being primarily in *g* (i.e., Spearman’s hypothesis). A practical implication of this is that good measures of *g* are needed to capture the full statistical effects of cognitive ability in context to research on eduPGS and related variables. Moreover, the positive manifold of Jensen Effects is at least consistent with a causal model. To reconcile it with a confounding model one would need to additionally propose a mechanism by which environmental induced phenotypic differences produced Jensen Effects (e.g., Flynn, 2019).

Considering these results, it may be that individual ancestry tracks cognitive ability in admixed populations across the Americas. If so, proper interpretation of the predictive accuracy of eduPGS in American admixed populations will require a better understanding of the causal pathways underwriting this association. We argue that admixture mapping is the appropriate next step (for a rationale, see: Kirkegaard et al., 2019). That said, different sub-ancestral components (e.g., North vs. South Amerindian / European, West versus East African) may yield different associations between global ancestry and cognitive ability, so additional global ancestry studies are warranted to better understand the pattern of effects.

## 5.2 Limitations

Owing to the sample sizes of the subgroups, some of the ancillary analyses, such as the comparison of eduPGS validities, were underpowered. Replications should be attempted with larger samples; until then, caution is warranted regarding interpretation. Importantly, we did not attempt to determine the cause of the covariance between eduPGS, ancestry, and *g*. We can only say that the cause is presently undetermined and that it needs to be resolved for a proper interpretation of the predictive accuracy of eduPGS in admixed American populations. While we suggest local admixture mapping as a way to narrow the uncertainty, further research should also explore other forms of confounding.

## Acknowledgements

Drs. Gur, Hakonarson, and collaborators request that publications resulting from these data cite their original publication: [TBD]. Support for the collection of the data sets was provided by grant RC2MH089983 awarded to Raquel Gur and RC2MH089924 awarded to Hakon Hakonarson. All subjects were recruited through the Center for Applied Genomics at The Children’s Hospital in Philadelphia. dbGaP accession <u>phs000607.v1.p1</u>

We also thank Jordan Lasker for advice on running the Multi-group Confirmatory Factor Analysis.

## Footnotes

1 Regarding terminology, since the relation between ancestry and eduPGS is better characterized as constitutive, rather than causal, eduPGS could be better characterized as being a component with constitutive explanatory relevance (Craver, 2007; Ylikoski, 2013; Weinberger, 2019), rather than a mediator as defined by Pearl (2014).

