## Supplementary material for "Intelligence-associated Polygenic Scores Predict g, Independent of Ancestry, Parental Educational Levels, and Color among Hispanics in comparison to European, European-African, and African Americans": SM 3

Updated: July 5, 2020

Supplementary File 3 for “Intelligence-associated Polygenic Scores Predict g, Independent of Ancestry, Parental Educational Levels, and Color among Hispanics in comparison to European, European-African, and African Americans”:

**Calculating Expected Phenotypic Differences.**

1. Formulas

The formal relation between the combined within and between group heritability, heritability between groups, and genetic and phenotypic differences is given by Defries (1972a), McClearn and Defries (1973), and Loehlin, Lindzey, & Spuhler (1975):

${h^{2}}_{\text{G}} = h^{2}* \frac{r}{t}$	(1)

where ${h^{2}}_{\text{G}}$ is the between group heritability, $h^{2}$is the combined heritability, *r* is the genetic intraclass correlation, and *t* is the phenotypic intraclass correlation, which is equivalent to the square of the point biserial correlation (i.e., *r*_pbs_^2^). This formula can be expressed in terms of within groups heritability ${h^{2}}_{\text{w}}$:

${h^{2}}_{\text{G}} = {h^{2}}_{\text{w}} * \frac{\left( 1-t \right)r}{\left( 1-r \right)t}$	(2)

where ${h^{2}}_{\text{w}}$is the average of the heritabilities within both groups. Equations (1) & (2) are simplified, but can be expanded to include gene-environment covariance (COV_GE_) (Defries, 1972b). In that case, the between group heritability is not ${h^{2}}_{\text{G}}$but is equal to:

${h^{2}}_{\text{G}}$+ *h*_G_ * *e*_G_ * *r*A_G_E_G_

where h_G_ and e_G_ are the square root of the between groups heritability and between group environmentality, respectively, and *r*A_G_E_G_ is the gene-environment correlation between groups. Thus,

in the case of positive COV_GE_, equations (1) and (2) will underestimate genetic differences between groups (McClearn and Defries, 1973). This formula can be further expanded to include other non-additive genetic variance components such as dominance (e.g., Wright, 1952). See the exchange between Defries and Jensen (Jensen, 1972) for when narrow or broad-sense within groups heritability is more appropriate. Here we will work with the simplified equation.

The intraclass correlations (*r* and *t*) can be interpreted in terms of one-way analysis of variance, where:

$ICC=\frac{MSb-MSw}{MSb + \left( n-1 \right)\mathrm{MS}w}$	(3)

where *MS*b represents the mean square between groups and *MS*w represents the mean square within groups. ICCs are equivalent to
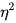
, which can be converted into Cohen’s *d* with the following equation:

$=\frac{(.5{d)}^{2}}{1 + {(.5d)}^{2}}$	or, equivalently, $d =2\surd\frac{}{1-}$	(4)
where Cohen’s *d* is:

$d=\frac{M1-M2}{{SD}_{pooled}}$	(5)

and *M*_1_ and *M*_2_ are the means for group 1 and group 2, respectively and SD_pooled_ is the pooled standard deviation. Alternatively,
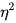
can be converted into a point-biserial correlation, and this can be converted into Cohen’s *d* with the following equation:

*r*_pbs_ = $\frac{d}{{{\surd(d}^{2}}_{\text{ }}+ 4 )}$	or, equivalently, *d* =	$\frac{2r}{{{\surd(1-r}^{2}}_{\text{ }})}$	(6)

Now, for diploid populations, *r*, the genetic intraclass correlation, in equation (1) and (2), is approximately 2*Fst* for the trait relevant alleles (Defries, 1972).

Though, more exactly it is:

*r* = 2F_st_ / (1+ F_IT_)	(7)

where F_st_ is the fixation index, or the between group variance in allele frequencies, and F_IT_ is the overall level of inbreeding in the total population (Hamilton, 1971; Cheverud, 1985).

Equation (2) can be rearranged to solve for *t* (the phenotypic variance). This gives:

$t = {h^{2}}_{\text{w}} * \frac{r}{{{rh}^{2}}_{\text{w}} + {h^{2}}_{\text{G}} - {{rh}^{2}}_{\text{G}}}$	(8)

Based on equation (8), one can solve for the expected gap, where environments are equal, which is done by setting ${h^{2}}_{\text{G}}$ to 1. This gives the following:

$t_{\text{expected}} = {h^{2}}_{\text{w}} * \frac{r}{{{rh}^{2}}_{\text{w}} + 1 - r}$	(for ${h^{2}}_{\text{G}}$ =1)	(9)

When the within groups heritability is high and/or the *Fst* value is low, this is approximately equal to:

$t_{\text{expected}} \sim{h^{2}}_{\text{w}}*r$	(for ${h^{2}}_{\text{G}}$ =1)	(10)

When $t_{\text{expected}}$ is converted into an effect size which expresses the relation between *x* and *y* in a linear form, this is called a “genotypic gap”.

Equation (9) and (10) can be related to the equation for expected differences given by Turkheimer (1991, eq. 6), where:

P_1observed_ = ${{\surd h}^{2}}_{\text{w}} Ĥ_{\text{1}}$ + ${{\surd e}^{2}}_{\text{w}} Ê_{\text{1}}$	and	P_2observed_ = ${{\surd h}^{2}}_{\text{w}} Ĥ_{\text{2}}$ + ${{\surd e}^{2}}_{\text{w}} Ê_{\text{2}}$	(11)

and P_1_ and P_2_ are the standardized observed phenotypic values for group 1 and group 2, respectively and $Ĥ$and $Ê$ are the standardized genetic and environmental values for the respective groups.

When ${Ê_{\text{1}}= Ê}_{\text{2}}$, and so ${h^{2}}_{\text{G}}$ = 1:

P_1observed -_P_2observed_ = *r*_pbs_observed_ = P_1expected -_P_2expected_ *= r*_pbs_expected_phenotypic_	(12)

And so:

*r*_pbs_expected_phenotypic_ = ${{\surd h}^{2}}_{\text{w}} {(Ĥ}_{\text{1}}{- Ĥ}_{\text{1}}$) = ${{\surd h}^{2}}_{\text{w}}$* *r*_pbs_genetic_	(13)

Except, since we now include between group variance, ${{\surd h}^{2}}$ needs to replace ${{\surd h}^{2}}_{\text{w}}$ in equation (13), giving:

*r*_pbs_expected_phenotypic_ = ${\surd h}^{2} {(Ĥ}_{\text{1}}{- Ĥ}_{\text{1}}$) = ${\surd h}^{2}$* *r*_pbs_genetic_	(14)

which, when squaring both sides, approximates equation (10). From the above, it can be seen that the $d_{\text{expected}}$ or the “genotypic gap” is equal to √ ${h^{2}}_{\text{G}}$ * d_observed,_ where the √ ${h^{2}}_{\text{G}}$ can be interpreted as the correlation between phenotype and genotype between groups, i.e.:

$d_{\text{expected}}= \sqrt{{h^{2}}_{\text{G}}}* d_{\text{observed}}$	(15)

This is because we can rewrite equation (1) as:

${h^{2}}_{\text{G}}*t= h^{2} * r$	(16)

Which by (14) gives:

${{\surd h}^{2}}_{\text{G}}*\sqrt{t}= {\surd h}^{2} * r_{\text{pbs\_genetic}} = r_{\text{pbs\_phenotypic}}$	(17)

and recovers the results from equation (8).

Note, it is obvious that ${h^{2}}_{\text{G}}$ is not equal to the real-world percentage of a difference that is attributable to genes. The inference makes the r^2^ interpretative fallacy (Hunter & Schmidt, 2004), which results because variance-explained does not represent a linear relation between x and y. Rather percentage genetic, in the ordinary sense, is given by:

Percentage genetic =	$d_{\text{expected}}$ / d_observed_	(18)

2. Education SNP Fst values:

Using the 1000 Genomes data, we computed the Fst values for the 10k MTAG

SNPs. These are shown in Table 1 below.

Table 1. Fst Values for the 10k MTAG SNPs by 1000 Genomes Population Pairs.

_____________________________________________________________________________________

| Population_1 | Population_2 | Edu_Fst |  | Edu_Fit |
| --- | --- | --- | --- | --- |
| AFR | EAS | 0.1402 |  | 0.1470 |
| AFR | EUR | 0.1090 |  | 0.1153 |
| AFR | SAS | 0.1018 |  | 0.1125 |
| AFR | AMR | 0.0984 |  | 0.1160 |
| EAS | EUR | 0.0964 |  | 0.1030 |
| AMR | EAS | 0.0714 |  | 0.0899 |
| EAS | SAS | 0.0626 |  | 0.0741 |
| EUR | SAS | 0.0342 |  | 0.0451 |
| AMR | SAS | 0.0296 |  | 0.0528 |
| AMR | EUR | 0.0226 |  | 0.0412 |

_____________________________________________________________________________________

*Note*: AFR = African, EAS = East Asian, SAS = South Asian, Eur = European, and AMR = admixed American (Mexican, Puerto Ricans, Colombian, and Peruvian) populations.

1. Example:

For Africans (AFR) and European (EUR), the MTAG SNPS the FST = .1090. By equation (7), *r* = 2(.1090) /(1+.1153) = .1955. Given a ${h^{2}}_{\text{w}}$ *=* .5, then t_expected_ from equation (9) is:

t_expected_  =    .5   *  $\frac{.1955}{(.1955)(.5) + 1 - (.1955)}$       =    .1083

Given equations (3) and (4), this equals *d* = 0.68 or a 10.21 point difference on a metric with a standard deviation of 15.

And so, by equation (17), the percentage of the observed group differences which is attributable to genes would be = $10.21$ / d_observed_
