## Supplementary material for "Intelligence-associated Polygenic Scores Predict g, Independent of Ancestry, Parental Educational Levels, and Color among Hispanics in comparison to European, European-African, and African Americans": SM 1

Updated: July 5, 2020

Supplementary File 1 for “Intelligence-associated Polygenic Scores Predict g, Independent of Ancestry, Parental Educational Levels, and Color among Hispanics in comparison to European, European-African, and African Americans”:

1. **Psychometric Assessment of the Penn Computerized Neurocognitive Battery**

Multi-group confirmatory factor analysis (MGCFA) is a technique often used to assess the measurement invariance (MI) of different psychological assessments including intelligence tests (Dolan, 2000; Dolan & Hamaker, 2001; Lubke, Dolan & Kelderman, 2001; Frisby & Beaujean, 2015). When MI holds, the assessment being examined is generally considered unbiased in the groups being compared. Resultantly, MI is taken to imply that the constructs measured in both groups are alike. MI is assessed by fitting a series of increasingly restrictive models and analyzing model fit at each step (van de Schoot, Lugtig & Hox, 2012). We assessed MI for Hispanic and European American comparisons with the Penn Computerized Neurocognitive Battery (PCNB).

Model fit was assessed with multiple indices. We adopted the same procedure, model, and criteria as in Lasker, Pesta, Fuerst, & Kirkegaard (2019), who examined MI for non-Hispanic European Americans and non-Hispanic African Americans. For assessing measurement invariance, we adopted Cheung and Rensvold’s (2002) and Chen’s (2007) criteria. Specifically, we regarded a ΔCFI of greater than -0.01, a ΔMc greater than -0.02, and a ΔRMSEA greater than 0.01 as evidence that measurement invariance was untenable. See Putnick & Bornstein (2016) for review.

For this analysis, we ran MGCFA on the set of individuals with sufficient cognitive data for imputation (506 Hispanic and 4,914 European Americans) without limiting the analysis to only those who passed quality controls for computing genetic ancestry. This set includes individuals with imputed subtest scores (when fewer than half of the subtests were missing). Our results are presented in tables S1-S2 and the **lavaan** model syntax is provided at the end of this supplement.

**Table S1. Bifactor Solution for Hispanic and European Americans on the Philadelphia Computerized Neurocognitive Battery.**

| Model | MI Step | χ2 | df | CFI | ΔCFI | RMSEA | ΔRMSEA | Mc | ΔMc | SRMR |
| --- | --- | --- | --- | --- | --- | --- | --- | --- | --- | --- |
| 1 | Configural | 115.45 | 48 | 0.992 | - | 0.023 | - | 0.994 | - | 0.013 |
| 2 | Metric | 135.60 | 65 | 0.991 | -0.001 | 0.020 | -0.003 | 0.994 | 0 | 0.015 |
| 3 | Scalar | 171.63 | 71 | 0.988 | -0.003 | 0.023 | 0.003 | 0.991 | -0.003 | 0.017 |
| 4 | Strict | 252.43 | 81 | 0.979 | -0.009 | 0.028 | 0.005 | 0.984 | -0.007 | 0.020 |
| 5 | Latent Variances | 252.43 | 81 | 0.979 | 0 | 0.028 | 0 | 0.984 | 0 | 0.020 |
| 6 | Means | 376.71 | 85 | 0.964 | -0.015 | 0.036 | 0.008 | 0.973 | 0.011 | 0.029 |
| 5a | Strong | 268.94 | 84 | 0.977 | -0.002 | 0.029 | 0.001 | 0.983 | -0.001 | 0.021 |
| 5b | Weak | 254.38 | 82 | 0.979 | 0 | 0.028 | 0 | 0.984 | 0 | 0.021 |
| 5c | Contra | 374.09 | 83 | 0.964 | -0.015 | 0.036 | 0.008 | 0.974 | -0.010 | 0.029 |

*Note*: Combined *N* = 5,420, with 506 Hispanic Americans and 4,914 European Americans. The latent variance model merely changes the identification constraint to the variances from a single loading for each modeled factor.

Models 5a to 5c further assess Spearman's hypothesis (Jensen, 1998; Frisby & Beaujean, 2015), which is more fully discussed by Lasker et al. (2019). The strong model (5a) leaves only *g* to vary between groups. The weak model (5b) leaves *g* to vary and constrains complex cognition; it should be noted that it was possible to constrain any set of the broad factors without a meaningful decrease in model fit (ΔCFI for weak models ranged from 0 to -0.001 out of all six possible models). The contra model (5c) constrains *g* and carries the broad factor constraints from the weak model. Contra model fits were always worse (approximately and absolutely) than the fits of comparable weak models (ΔCFI ranged from -0.005 to -0.015). 5a-c are each compared to model 5. Neither the strong nor weak model fits worse than the model with latent variances constrained in terms of approximate fit; however, using a χ2 test, the weak model does not fit worse while the strong model does (like the contra model). The fit for the chosen contra model could be rejected with a ΔCFI of <-0.01 accompanied by a notably elevated χ^2^. These results tentatively support either the weak or strong model over the contra model with approximate fits and absolutely support the weak model with a χ^2^ test.

Tables S2 and S3 show. respectively, the standardized mean differences based on the model with constrained latent variances and the weak Spearman’s hypothesis model. In the weak Spearman’s hypothesis model, the European-Hispanic American difference in *g* is 0.668 *g* (positive values favor European Americans and vice-versa). There are also small to moderate differences in executive functioning (*g* = -0.342) and episodic memory (*g* = -0.234) net of *g* which favor Hispanics. In the contra or baseline models, broad factors more strongly favor European Americans, as the differences associated with *g* in the latent variances or weak models are distributed among the other factors in the absence of *g*.

The values of ω_h_ and ω_t_ for this battery were 0.69 and 0.77 respectively; 90% of the reliable variance was thus attributable to *g*. The ECV for *g* was 70%, PUC was 0.78, and H was 0.76, with these values being uniformly too low for complex cognition (ECV = 9%, H = 0.27), executive functioning (9%, 0.24%), and episodic memory (12%, 0.31). Using the method from Dolan (2001), in the latent variances model, an average of 67% of the between-group differences were accounted for by *g*; 65% of the differences in the indicators for complex cognition, 62% for executive functioning, and 58% for episodic memory.

In the selected weak Spearman’s hypothesis model, 73% of the group differences are accounted for by g and the proportion of the differences in the indicators for executive functioning were unchanged. The correlation between the vector of group differences and the vector of *g* loadings is *r* = 0.524. This same correlation for the complex cognition, executive functioning, and episodic memory loadings are, respectively, *r* = -0.420, *r* = 0.113, *r* = -0.198, and overall, *r* = -0.159 with all values and *r* = -0.269 without the 0 for PCPT. Values for Mardia’s b1p and b2p were 16.403 and 132.853 (Mardia, 1980).

***Table S2. Factor Score Differences between Hispanic and European Americans based on the Model with Constrained Latent Variances***

| Factor | Estimate | SE | Lower 95% CI | Upper 95% CI |
| --- | --- | --- | --- | --- |
| g | 0.614 | 0.071 | 0.474 | 0.754 |
| Complex Cognition | 0.123 | 0.088 | -0.049 | 0.296 |
| Executive Functioning | -0.259 | 0.126 | -0.506 | -0.012 |
| Episodic Memory | -0.192 | 0.078 | -0.344 | -0.040 |

*Note: Positive values indicate higher white scores and vice-versa. Estimates are in terms of Hedge’s g.*

***Table S3. Factor Score Differences between Hispanic and European Americans based on the Weak Spearman’s Hypothesis Model***

| Factor | Estimate | SE | Lower 95% CI | Upper 95% CI |
| --- | --- | --- | --- | --- |
| g | 0.668 | 0.064 | 0.543 | 0.793 |
| Complex Cognition | 0 | - | - | - |
| Executive Functioning | -0.342 | 0.124 | -0.585 | -0.099 |
| Episodic Memory | -0.234 | 0.075 | -0.382 | -0.087 |

*Note: Positive values indicate higher white scores and vice-versa. Estimates are in terms of Hedge’s g.*

2. **Subtest Means and Standard Deviations by Hispanic Subgroup**

We additionally provide the means and standard deviations for the subtests by Hispanic subgroup. Scores for all 15 subtests are provided, with an asterisk placed next to the 10 for which measurement invariance was found to hold.

***Table S4. Subtest Means and Standard Deviations by Hispanic Subgroup***

_____________________________________________________________________________________________

|  | **HI** |  | **HI_EA** |  | **HI_AA** |  | **HI_OT** | | **Other** |  |
| --- | --- | --- | --- | --- | --- | --- | --- | --- | --- | --- |
| g | -0.57 | 1.13 | -0.33 | 1.17 | -0.84 | 0.98 | -0.65 | 1.17 | -0.39 | 1.13 |
| PLOT* | -0.36 | 1.04 | -0.17 | 1.05 | -0.62 | 1.03 | -0.40 | 1.03 | -0.21 | 0.98 |
| PCPT* | -0.26 | 1.26 | -0.19 | 1.14 | -0.29 | 1.32 | -0.46 | 1.38 | -0.46 | 1.17 |
| PCET* | -0.26 | 1.08 | -0.25 | 1.12 | -0.43 | 1.04 | -0.25 | 1.06 | -0.09 | 1.08 |
| LNB* | -0.29 | 1.12 | -0.23 | 1.12 | -0.42 | 1.19 | -0.33 | 0.99 | -0.15 | 1.13 |
| VOLT* | -0.27 | 1.06 | -0.08 | 1.00 | -0.51 | 1.09 | -0.33 | 1.10 | -0.12 | 0.99 |
| TAP | 0.02 | 0.98 | 0.14 | 0.96 | 0.07 | 0.91 | -0.07 | 1.09 | -0.05 | 0.97 |
| PMRT* | -0.25 | 0.98 | -0.13 | 1.02 | -0.39 | 0.89 | -0.25 | 1.00 | -0.22 | 1.02 |
| MP | -0.07 | 1.18 | -0.02 | 1.17 | 0.05 | 0.83 | -0.31 | 1.65 | -0.02 | 0.99 |
| PEDT | -0.15 | 1.09 | 0.02 | 1.15 | -0.21 | 1.17 | -0.25 | 1.06 | -0.13 | 0.95 |
| PVRT* | -0.58 | 1.18 | -0.35 | 1.23 | -0.74 | 1.14 | -0.67 | 1.28 | -0.52 | 1.06 |
| PEIT | 0.02 | 1.02 | -0.10 | 1.13 | -0.01 | 0.91 | 0.12 | 1.13 | 0.08 | 0.91 |
| PFMT* | 0.06 | 1.04 | -0.03 | 1.02 | -0.02 | 1.03 | 0.32 | 0.98 | -0.01 | 1.09 |
| PADT | -0.06 | 1.02 | 0.03 | 1.04 | -0.13 | 1.11 | -0.09 | 1.00 | -0.02 | 0.92 |
| PWMT* | -0.11 | 1.16 | 0.06 | 0.99 | -0.09 | 1.00 | -0.37 | 1.47 | -0.03 | 1.12 |
| WRAT* | -0.48 | 1.05 | -0.34 | 1.01 | -0.75 | 1.04 | -0.52 | 1.01 | -0.25 | 1.07 |

*_____________________________________________________________________________________________*

*Note*: *Denotes the subtests, from the 10-subtest measurement invariant model, used to compute *g* scores. HI_EA = Hispanic European, HI_AA = Hispanic African, HI_EA = Hispanic Other, and Other = any other also with Hispanic ethnicity marked.

*References*

Chen, F. F. (2007). Sensitivity of Goodness of Fit Indexes to Lack of Measurement Invariance. *Structural Equation Modeling: A Multidisciplinary Journal*, *14*(3), 464–504.

Cheung, G. W., & Rensvold, R. B. (2002). Evaluating Goodness-of-Fit Indexes for Testing Measurement Invariance. *Structural Equation Modeling: A Multidisciplinary Journal*, *9*(2), 233–255.

Dolan, C. V. (2000). Investigating Spearman’s Hypothesis by Means of Multi-Group Confirmatory Factor Analysis. *Multivariate Behavioral Research*, *35*(1), 21–50.

Dolan, C. V., & Hamaker, E. L. (2001). *Investigating black-white differences in psychometric IQ: Multi-group confirmatory factor analyses of the WISC-R and KABC and a critique of the method of correlated vectors*.

Frisby, C. L., & Beaujean, A. A. (2015). Testing Spearman’s hypotheses using a bi-factor model with WAIS-IV/WMS-IV standardization data. *Intelligence*, *51*, 79–97.

Jensen, A. R. (1998). *The g Factor: The Science of Mental Ability*. In *The g Factor: The Science of Mental Ability*. Westport, CT, US: Praeger Publishers/Greenwood Publishing Group.

Lasker, J., Pesta, B. J., Fuerst, J. G. R., & Kirkegaard, E. O. W. (2019). Global Ancestry and Cognitive Ability. *Psych*, *1*(1), 431–459.

Lubke, G. H., Dolan, C. V., & Kelderman, H. (2001). Investigating Group Differences on Cognitive Tests Using Spearman’s Hypothesis: An Evaluation of Jensen’s Method. *Multivariate Behavioral Research*, *36*(3), 299–324.

Mardia, K. V. (1980). Tests of unvariate and multivariate normality. In *Analysis of Variance*: *Vol.* *1*. *Handbook of Statistics* (pp. 279–320).

Putnick, D. L., & Bornstein, M. H. (2016). Measurement invariance conventions and reporting: The state of the art and future directions for psychological research. *Developmental Review*, 41, 71-90.

van de Schoot, R., Lugtig, P., & Hox, J. (2012). A checklist for testing measurement invariance. *European Journal of Developmental Psychology*, 9(4), 486–492.

Model Syntax

F1 =~ WRAT + PVRT + PMAT

F2 =~ PMAT + PCET + VOLT + PLOT

F3 =~ VOLT + PWMT + PFMT + LNB

g =~ WRAT + PVRT + PMAT + PCET + VOLT + PLOT + LNB + PWMT + PFMT + PCPT
