## Supplementary material for "Intelligence-associated Polygenic Scores Predict g, Independent of Ancestry, Parental Educational Levels, and Color among Hispanics in comparison to European, European-African, and African Americans": SM 2

Updated: September 20, 2020

Supplementary File 2 for “Intelligence-associated Polygenic Scores Predict g, Independent of Ancestry, Parental Educational Levels, and Color among Hispanics in comparison to European, European-African, and African Americans”

Evaluation of Bias and Validity Using the 1000 Genomes populations

In these analyses, we leverage the 1000 Genomes populations to investigate whether

eduPGS differences can be accounted for by bias in SNP Betas (βs) & SNPs, whether the differences can be accounted for by the inclusion of SNPs with trans-ethnically discordant β weights, and whether eduPGS have cross-national predictive validity.

Analysis 1. Differences in population-GWAS vs. Within Family Weighted eduPGS

Rationale:

PGS scores are calculated by weighting the trait-associated SNP allele frequencies by each SNP’s effect on the predicted trait (i.e., the SNP βs). However, population structure may bias the βs. This form of bias can be partially circumvented by using βs calculated from within-family analyses, which, in principle, are robust to the effects of population structure (Sohail et al., 2019). Thus, in this analysis, we compare the differences between eduPGS computed with population-GWAS vs. within family βs weights.

Method:

Lee et al. (2018) report the βs for the 10k MTAG SNPs based on their analysis of 1.1 million (mostly) unrelated individuals. The predicted traits were cognitive ability, self-reported math ability, and highest math class taken. On request, the authors also provided the βs for their analysis of 22,000 sibling pairs. The predicted trait was self-reported years of education. We applied these weights to the SNP frequencies for the 1000 Genomes Northern and Central European descent from Utah (CEU) and Yoruba Nigerian (YRI) samples. We used Europeans and Africans because the ancestral populations of the admixed groups were primarily European and African in origin. Before doing this, we filtered the 10k MTAG SNPs to those for which both population-GWAS and within family βs were available. Moreover, the SNPs were filtered for MAF >0.01 for both CEU and YRI. We then computed eduPGS based on the within family and population-GWAS βs separately for each individual. We further repeated this analysis for the 5 unadmixed European and 5 unadmixed African 1000 Genomes populations.

Results:

Figure S1 and S2 depict, respectively, the population-GWAS and within family weighted eduPGS for CEU and YRI individuals. The difference in betas came to β = 1.66 and β = 1.18, respectively. (Note, these βs are based on total sample *SDs*, which are larger than the average of the within population *SDs*). In both cases, the differences were highly significant and large by conventional interpretative standards. This difference between the two represents a 29% reduction in the eduPGS gap size. For comparison, Lee et al. (2018) report that the within-family effect sizes are 40% smaller than the population-GWAS ones (among Europeans). Thus, this magnitude of reduction in gap size is consistent with the reduction in validity among Europeans.

Figure S1: Plot of population-GWAS Weighted, MTAG SNP eduPGS for CEU and YRI 1000 Genomes Individuals.

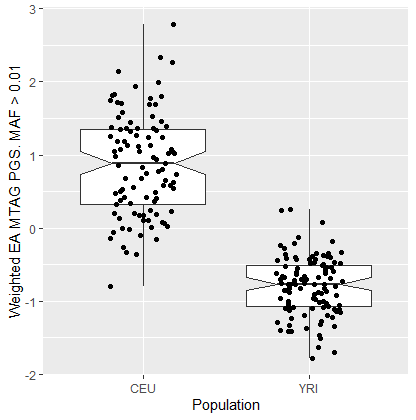

Figure S2: Plot of Within-family, Weighted MTAG SNP eduPGS for CEU and YRI 1000 Genomes Individuals.

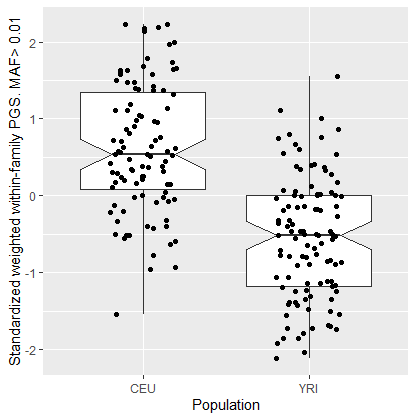

Further, Table S1 gives the eduPGS means for all 10K MTAG SNPs, for the derived SNPs, and for the ancestral SNPs. For all sets, the differences are significant and large, as determined using Welch’s Two Sample *t*-test.

Table S1. Mean MTAG-based PGS for CEU and YRI Calculated using population-GWAS and Within Family Betas.

|  | W/ population-GWAS |  | W/ Within Family Betas |  |
| --- | --- | --- | --- | --- |
|  | CEU (N = 99) | YRI (N = 108) | CEU (N = 99) | YRI (N = 108) |
| All SNPS | 0.866 | -0.794 | 0.614 | -0.563 |
| *p*-value (Welch’s Two Sample *t*-test) |  | < 0.0001 |  | < 0.0001 |
| Derived SNPs | 0.938 | -0.860 | 0.702 | -0.643 |
| *p*-value (Welch’s Two Sample *t*-test) |  | < 0.0001 |  | < 0.0001 |
| Ancestral SNPs | 0.605 | -0.554 | 0.528 | -0.484 |
| *p*-value (Welch’s Two Sample *t*-test) |  | < 0.0001 |  | < 0.0001 |

*Note*: SNPs were filtered for MAF >0.01 for both CEU and YRI. Scores represent standard scores calculated using the standard deviation in the total sample. Sample sizes for the *t*-test were N = 99 for CEU and N=108 for YRI.

Further, to see if use of MTAG SNPs were biasing the results, we computed the differences using the 4,413 within-family SNPs that had a *p*-value < .05 along with the within family weights. These thus are pure within-family based eduPGS and so should show no population structure related bias. The βs for CEU and YRI were 0.529 and -.485, respectively with a β difference of 1.01. Results are shown in Figure S3.

Figure S3: Plot of Within-family Weighted, Within-Family SNP eduPGS for CEU and YRI 1000 Genomes Individuals.

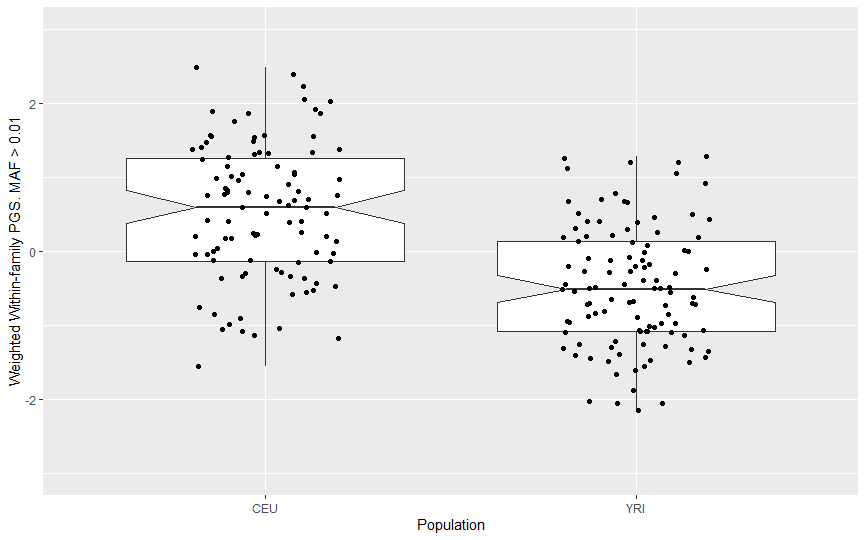

We additionally computed the MTAG-based PGS for the 5 African and 5 European 1000 Genomes populations. These results are shown below in Table S2. As seen in Table S2, there are large eduPGS differences between the 1000 Genomes European and African populations. The European-African difference in betas for population-GWAS and within family weighted MTAG PGS came to β = 1.66 and β = .93, respectively. This difference between the two represents a 44% reduction in the eduPGS gap size. Note, the standardized scores were computed using the total sample, so the scores for CEU and YRI are different in Table S1 (with 2 populations) than in S2 (with 10 populations); also, the standard deviations (within populations) is less than 1.0, because a significant portion of the variance was between populations, so the β difference is not an effects size which uses average or pooled *SDs*.

Table S2. Mean MTAG-based PGS for European and African 1000 Genomes populations calculated using population-GWAS and Within Family Betas.

__________________________________________________________________________________

| Population | *N* | MTAG SNP Frequencies (Unweighted)  *M* | *SD* | MTAG Standard Scores (population-GWAS Weights)  *M* | *SD* | MTAG Standard Scores (Within Family Weights) *M* | *SD* |
| --- | --- | --- | --- | --- | --- | --- | --- |
| CEU | 99 | 0.510 | 0.0063 | 0.800 | 0.6550 | 0.707 | 0.7530 |
| FIN | 99 | 0.510 | 0.0105 | 0.876 | 0.8480 | 0.345 | 1.0700 |
| GBR | 91 | 0.509 | 0.0064 | 0.694 | 0.6310 | 0.498 | 0.8080 |
| IBS | 107 | 0.511 | 0.0059 | 0.949 | 0.6440 | 0.359 | 0.7670 |
| TSI | 107 | 0.509 | 0.0055 | 0.806 | 0.5560 | 0.439 | 0.8120 |
| EUR_Average | 503 | 0.510 | 0.0071 | 0.829 | 0.6723 | 0.467 | 0.8488 |
| YRI | 108 | 0.494 | 0.0040 | -0.768 | 0.3740 | -0.365 | 0.7240 |
| ESN | 99 | 0.492 | 0.0043 | -0.839 | 0.4200 | -0.405 | 0.7100 |
| GWD | 113 | 0.493 | 0.0039 | -0.795 | 0.4010 | -0.553 | 0.6660 |
| LWK | 99 | 0.492 | 0.0055 | -0.839 | 0.4740 | -0.522 | 1.4900 |
| MSL | 85 | 0.492 | 0.0040 | -0.919 | 0.3930 | -0.484 | 0.6990 |
| AFR_Average | 504 | 0.493 | 0.0044 | -0.827 | 0.4133 | -0.466 | 0.9107 |

__________________________________________________________________________________

*Note*: SNPs were filtered for MAF >0.01 for both CEU and YRI. Scores represent standard scores calculated using the means and standard deviation in the total sample. Note also, populations *SDs* are less than 1 because these *SDs* include only within-population variance.

Interpretation:

Though reduced in size, the within-family weighted eduPGS differences remain large. Moreover, this magnitude of reduction is roughly consistent with the reduced effect sizes that within-family eduPGS have, as compared to population-GWAS ones, among Europeans. Generally, the differences are unlikely to be due to population structure-related bias in the SNP βs. Moreover, they are unlikely to be due completely to population structure-related to SNP selection, since differences were substantial when using both within family weights and SNPs.

Analysis 2. Effect of Transethnically Concordant versus Discordant alleles on 1000 Genomes Population eduPGS Differences

Rationale:

Previous polygenic selection studies have applied European GWAS βs to different world populations (Graham and Coop, 2014; Piffer, 2019). In theory, however, taking into account information about population-specific effects (i.e., βs for both European and non-European comparison samples) should yield more accurate results both on the individual and population levels (Márquez-Luna et al., 2017; Grinde et al., 2019).

Indeed, the PGS βs for SNPs in European samples will often show opposite or discordant effects in non-European samples. SNPs which show effects which are directionally concordant across ethnic groups are more likely to be causal (when individual differences are caused by variants common across ethnic groups). As such, computing PGS using SNPs with only concordant effects may either increase or decrease apparent eduPGS gaps. For this analysis, we use the two largest TCP samples, European and African Americans, to classify MTAG βs into trans-ethnically concordant and discordant ones. We then recomputed concordant and discordant eduPGS and compare the magnitude of the 1000 Genomes CEU and YRI differences.

Method:

Using the TCP sample, we computed the 10k MTAG SNP betas for *g* separately for European and African Americans. These results are provided in the supplementary excel file. The 10k MTAG SNPs were then split into two sets: 1) concordant SNPs, which had the same direction of effect for TCP African and European Americans and 2) discordant SNPs, which had opposite directions for TCP African and European Americans. Polygenic scores were then computed for the concordant SNPs only and the discordant SNPs only. In computing the eduPGS, the SNPs were weighted with the βs reported by Lee et al. (2018). The rationale was that GWAS βs are more reliable than the TCP βs because they are based on a much larger sample size, thus TCP data is only used to identify concordant / discordant SNPs status. For comparison, eduPGS were additionally computed, with the same weights, based on all 10k MTAG SNPS.

Results:

There were 4,307 and 3,994 concordant and discordant SNPs, respectively. This suggests there is a slight overrepresentation of concordant SNPs. The binomial probability of having 4,307 or more discordant SNPs out of 8,301 is *p* = 0.0003. Table S3 reports the eduPGS based on all SNPs, the concordant only, and discordant only.

Table S3. Polygenic Scores by Concordance Status and Population.

| Population | PGS 10k MTAG | PGS concordant | PGS discordant |
| --- | --- | --- | --- |
| CEU | 0.5063139 | 0.505047 | 0.5092652 |
| YRI | 0.4904064 | 0.4725855 | 0.5109076 |
| Difference (CEU - YRI) | 0.01590744 | 0.03246145 | -0.001642405 |

An independent two-group *t*-test was employed to assess the significance of the PGS differences between CEU and YRI as shown in Table S4. Both methods yielded significant PGS differences between CEU and YRI.

Table S4. Results of t-test for Polygenic Score CEU-YRI Difference.

|  | *T* | *P* value | 95% CI for Mean Difference | df |
| --- | --- | --- | --- | --- |
| PGS 10k MTAG | 6.2883 | 3.371*10-10 | 0.01093991, 0.02084975 | 8418 |
| PGS concordant | 9.013 | 2.2*10-16 | 0.02538434, 0.03949752 | 4305 |
| PGS discordant | -0.4527 | 0.6508 | -0.008755425, 0.005470615 | 3993 |

*Note*: Sample sizes for the t-test were the number of concordant and discordant SNPs. Using the number of individuals, instead, did not change the interpretation of the results.

As shown in Table S5, a two-way ANOVA showed the interaction to be significant.

Table S5. Two-way anova (Response: PGS)

|  | Sum of Squares | *F* value | *P* value |
| --- | --- | --- | --- |
| Population | 1.07 | 10.727 | 0.0010578 |
| Method (concordant vs discordant) | 1.88 | 18.811 | 1.451e-05 |
| Population*Method | 1.21 | 12.089 | 0.0005085 |

Interpretation:

The concordant PGS had a higher CEU-YRI difference than both the discordant PGS and the combined eduPGS. In fact, the discordant PGS showed no CEU-YRI difference (95% C.I. = -0.009, 0.005), while the concordant CEU-YRI difference was around 3% (95% C.I. = 0.025, 0.039). A two-way ANOVA showed this interaction to be significant. Generally, the results from this analysis are consistent with the hypothesis that the ability of the MTAG eduPGS to predict population differences in IQ and scholastic achievement is driven by the subset of SNPs with trans-ethnically homogeneous effect and that the discordant SNPs reduce the magnitude of the polygenic score differences. However, it needs to be noted that the power to correctly classify SNPs was low, so this analysis will have to be repeated when better data is available.

Analysis 3. Predictive Validity of the Polygenic Scores across Populations

Rationale:

Cross-population analyses are frequently used to assess the validity of PGS (e.g., Berg et al., 2019, Figure 1; Sohail et al., 2019; Figure 4). At times, it is found that PGS differences are directionally inconsistent with observed trait differences across populations. When this is found, the cross-population validity of the PGS is called into question (Martin et al., 2017).

Thus, in the second analysis, we compared the cross-population predictivity for measured population IQ / test scores.

In addition to 10K MTAG and MTAG-lead eduPGS, we look at the relation by concordant and discordant status, as it is expected that the discordant 10K MTAG eduPGS will be less predictive than the concordant one, as found on the individual level. Moreover, we see if eduPGS computed from Lee et al.’s (2018) sibling analysis data predicts national cognitive scores. To do so, we computed eduPGS using all 81,130 within-family SNPs and the 4,413 within-family SNPs that had a *p*-value < .05. Note, while sibling analyses are robust to population structure-related confounding, none of these SNPs met the minimum for GWAS significance (5e-8) and so the eduPGS computed from them provides a very noisy signal.

Method:

The Measured population cognitive scores for 18 countries were copied from Lynn and Becker (2019) and World Bank (2017). EduPGS were calculated for the 26 1000 Genomes populations, using the 10K MTAG, MTAG-lead SNPs, the concordant and discordant 10K MTAG SNPs, and within-family based weighted SNP frequencies. The latter were computed using all within-family SNPs along with the within family β weights. Scores for multiple ethnic groups were reported for four countries: USA (European, Mexican, African, and Asian-Indian American), UK (European, Indian, and Sri Lankan British), China (North Han, South Han, and Dai), and Nigeria (Esan and Yoruba). For these, eduPGS were weighted as shown in Table S6 to create national eduPGS. Intra-national group scores were not used as 1) scores are not psychometrically comparable to international ones (Wicherts & Wilhelm, 2007; Täht & Must, 2013), though when available they are reported with the sources noted, and 2) migrant populations can not be assumed to be representative of national ones.

Results:

Table S6 reports the eduPGS and country cognitive scores.

Table S6: Polygenic and Cognitive Scores at the Country level

___________________________________________________________________________

| Population |  | MTAG 10k eduPGS | MTAG Lead eduPGS | MTAG Concordant eduPGS | MTAG Discordant eduPGS | Lee et al.'s (2018) Within-Family (All) | Lee et al.'s (2018) Within-Family (*P* < .05) | Ethnic IQs | Lynn & Becker's (2019) NIQs | World Bank's (2017) Test Scores |
| --- | --- | --- | --- | --- | --- | --- | --- | --- | --- | --- |
| Afr.Car.Barbados | | -1.376 | -1.343 | -1.418 | -0.219 | 0.0000151 | 0.0002438 |  | 91.69/ 90.15* |  |
| Bengali Bangladesh | | -0.021 | -0.122 | 0.058 | -0.269 | 0.0000240 | 0.0003013 |  | 74.36 | 368 |
| Colombian |  | 0.368 | 0.305 | 0.557 | -0.614 | 0.0000266 | 0.0003316 |  | 82.99 | 424 |
| Finland |  | 1.043 | 0.986 | 1.117 | -0.187 | 0.0000246 | 0.0002973 |  | 100.55 | 548 |
| Gambian |  | -1.469 | -1.425 | -1.544 | -0.115 | 0.0000141 | 0.0002370 |  | 60 | 338 |
| Iberian, Spain | | 0.264 | 0.186 | 0.502 | -0.562 | 0.0000262 | 0.0003050 |  | 93.87 | 514 |
| Japan |  | 0.672 | 0.993 | 0.613 | 0.814 | 0.0000242 | 0.0003177 |  | 106.43 | 563 |
| Vietnam |  | 1.024 | 1.211 | 0.467 | 2.137 | 0.0000234 | 0.0003104 |  | 89.53 | 519 |
| Luhya, Kenya | | -1.513 | -1.400 | -1.615 | -0.058 | 0.0000145 | 0.0002505 |  | 75.22 | 455 |
| Mende, Sierra Leone | | -1.632 | -1.585 | -1.672 | -0.262 | 0.0000149 | 0.0002521 |  | 60 | 316 |
| Peruvian, Lima | | -0.500 | -0.676 | -0.047 | -1.540 | 0.0000286 | 0.0003246 |  | 81.42 | 407 |
| Punjabi, Pakistan | | 0.187 | 0.014 | 0.284 | -0.327 | 0.0000242 | 0.0002940 |  | 80.05 | 339 |
| Puerto Rican | | 0.230 | 0.190 | 0.477 | -0.925 | 0.0000251 | 0.0003124 |  | 81.89 |  |
| Toscani, Italy | | 0.907 | 0.796 | 1.105 | -0.707 | 0.0000263 | 0.0003210 |  | 94.16 | 514 |
| Nigeria |  | -1.471 | -1.317 | -1.637 | 0.203 | 0.0000157 | 0.0002475 |  | 67.83 | 325 |
| Esan, Nigeria | 0.01 | -1.545 | -1.364 | -1.609 | -0.167 | 0.0000142 | 0.0002343 | (??.??) |  |  |
| Yoruba, Nigeria | 0.2 | -1.467 | -1.315 | -1.638 | 0.222 | 0.0000158 | 0.0002481 | (??.??) |  |  |
| USA |  | 0.458 | 0.324 | 0.671 | -0.824 | 0.0000268 | 0.0003099 |  | 97.43 | 523 |
| Utah Whites | 0.63 | 0.924 | 0.748 | 1.139 | -0.787 | 0.0000286 | 0.0003221 | (100.00) |  |  |
| Mexican in L.A. | 0.11 | -0.077 | -0.181 | 0.293 | -1.306 | 0.0000273 | 0.0003029 | (91.45) |  |  |
| US Blacks | 0.13 | -1.347 | -1.301 | -1.262 | -0.664 | 0.0000178 | 0.0002575 | (85.00) |  |  |
| Gujarati Indian, Tx | 0.01 | 0.469 | 0.269 | 0.443 | 0.103 | 0.0000225 | 0.0003005 | (101.65) |  |  |
| China |  | 1.227 | 1.496 | 0.786 | 2.122 | 0.0000249 | 0.0003110 |  | 103.95 | 456 |
| Chinese, Bejing | 0.47 | 1.364 | 1.631 | 0.863 | 2.368 | 0.0000260 | 0.0003087 | (105.90) |  |  |
| Chinese, South | 0.47 | 1.089 | 1.360 | 0.708 | 1.876 | 0.0000238 | 0.0003134 | (105.90) |  |  |
| Chinese Dai |  | 0.739 | 1.047 | 0.491 | 1.271 | 0.0000231 | 0.0003004 | (93.90) |  |  |
| UK |  | 0.783 | 0.600 | 1.044 | -0.960 | 0.0000267 | 0.0003086 |  | 99.22 | 517 |
| British, GB | 0.82 | 0.790 | 0.609 | 1.060 | -0.993 | 0.0000268 | 0.0003090 | (100.00) |  |  |
| Indian Telegu, UK | 0.02 | 0.501 | 0.249 | 0.400 | 0.379 | 0.0000243 | 0.0002925 | (??.??) |  |  |
| Sri Lankan, UK | | 0.376 | 0.118 | 0.227 | 0.531 | 0.0000250 | 0.0003024 | (??.??) |  |  |

_________________________________________________________________________

*Note:* USA ethnic scores from Fuerst (2014), with second-generation “Hispanic” substituted for Mexican; Chinese ethnic scores from Lynn and Cheng (2014). USA eduPGS calculated as the weighted eduPGS of European, Mexican, African, and Asian-Indian Americans. Chinese eduPGS calculated as the weighted. eduPGS of North and South Han. UK eduPGS calculated as the weighted eduPGS of White and Indian eduPGS.Nigerian eduPGS calculated as the weighted eduPGS of Esan and Yoruba eduPGS (??.??) indicates unknown intranational scores. *Lynn and Becker (2019) report an IQ of 91.69 for Barbados. However, this becomes 90.15 when including the 114 generation 2 sample WASI scores reported by Wabler et al. (2018). We use this later score.

Table S6 reports the correlation matrix. As shown, the Lead SNP eduPGS has high predictive validity for both Lynn and Becker’s National IQs (*r* = .817) and World Bank’s Test Scores (*r* = .747). These values were equivalently high for the MTAG 10K eduPGS, at *r* = .804 and *r* = .734, respectively. As predicted, the correlations for the discordant eduPGS, unlike the concordant ones, were low at *r* = .213 (Lynn & Becker, 2019) and *r* = .142 (World Bank, 2017), respectively. The difference between the concordant (*r* = .784) and discordant (*r* = .213) correlations with Lynn and Becker’s NationalIQs was significant (*t* =5.93, *p* <0.01; dependent samples). The within-family based eduPGS, based on all SNPS, also had lower validity for both Lynn and Becker’s National IQs (*r* = .648) and World Bank’s Test Scores (*r* = .565), as did the ones that had a *p* < .05, with *r* = .638 (Lynn & Becker, 2019) and *r* = .596 (World Bank, 2017), respectively.

Figure S6. Correlation matrix for eduPGS and Population IQs.

__________________________________________________________________________________

|  | 1 | 2 | 3 | 4 | 5 | 6 |  |
| --- | --- | --- | --- | --- | --- | --- | --- |
| 1. MTAG 10K eduPGS | 1.00 |  |  |  |  |  |  |
| 2. MTAG Lead eduPGS | .988 (18) | 1.00 |  |  |  |  |  |
| 3. MTAG Concordant eduPGS | .971 (18) | .930 (18) | 1.00 |  |  |  |  |
| 4. MTAG Discordant eduPGS | .246 (18) | .371 (18) | .011 (18) | 1.00 |  |  |  |
| 5. Lee et al. (2018) Within Family | .840 (18) | .776 (18) | .921 (18) | -.190 (18) | 1.00 |  |  |
| 6. Lee et al. (2018) Within Family (*p* < .05) | .853 (18) | .814 (18) | .905 (18) | -.063 (18) | .964 (18) | 1.00 |  |
| 7. Lynn & Becker (2019) NIQ | .804 (18) | .817 (18) | .784 (18) | .213 (18) | .648 (18) | .638 (18) | 1.00 |
| 8. World Bank (2017) Test Scores | .734 (16) | .747 (16) | .727 (16) | .142 (16) | .565 (16) | .596 (16) | .885 (16) |

__________________________________________________________________________________

*Note:* Sample sizes in parentheses.

Figure S3 shows the regression plot for NIQ and MTAG 10K eduPGS, while figure S4 shows the regression plot for NIQ and the within-family based eduPGS. As seen, the cross population validity of the within-family eduPGS is tenuous. However, this eduPGS may not be reliable as the within-family SNP with the lowest *p* value (1.877e-05) did not even meet the conventional minimum for GWAS significance (5e-8).

*Figure S3. Regression Plot for Lynn and Becker’s (2019) NIQ and MTAG 10K eduPGS Scores Based on 1000 Genomes Samples.*

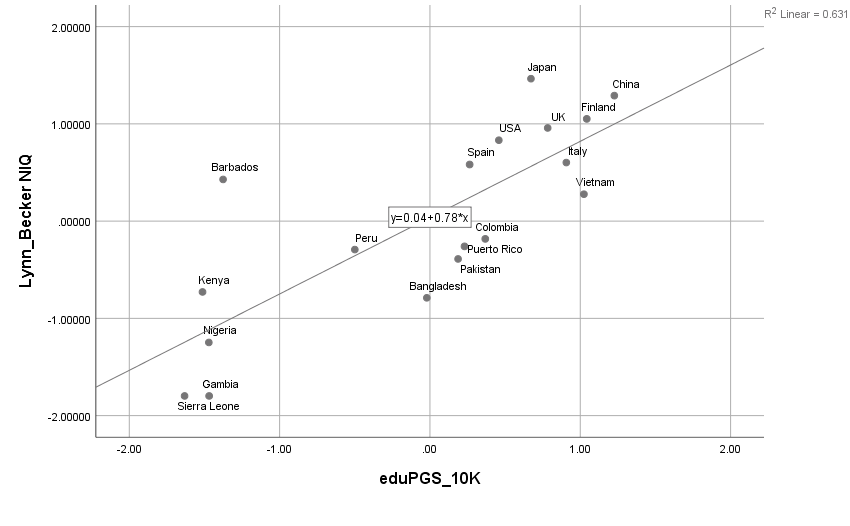

*Figure S4. Regression Plot for Lynn and Becker’s (2019) NIQ and Within-family Based eduPGS Scores (Using all 81,130 SNPs) Based on 1000 Genomes Samples.*

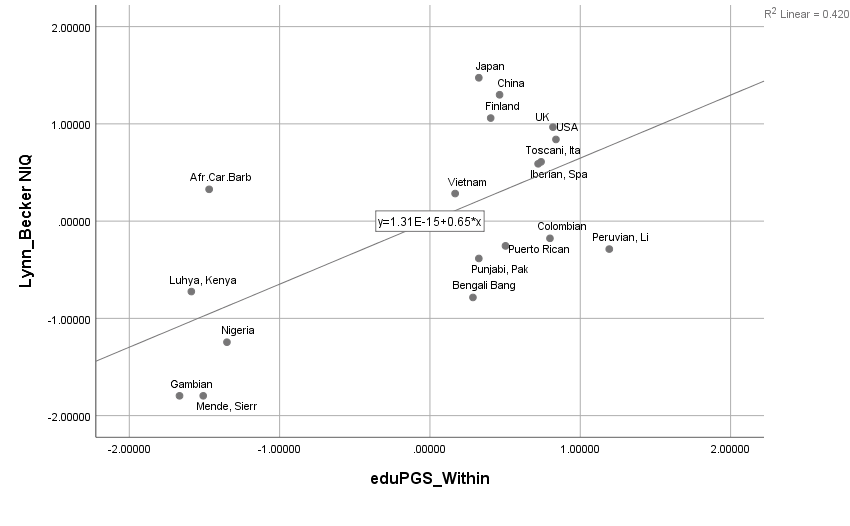

Interpretation:

MTAG-based eduPGS correlates highly with measured cognitive scores. Moreover, the population-level correlations were significantly higher for the concordant than discordant SNPs, consistent with the individual level results. However, the cross-population predictive validity for the within-family based eduPGS, calculated based on 22,000 sibling pairs, is tenuous, with high eduPGS for some low cognitive test scoring countries (e.g., Peru and Colombia). Since none of the SNPs for this eduPGS met the minimum for GWAS significance this may simply be a result of unreliability in the measure. In general, the cross-population validity of the MTAG-based eduPGS can not be rejected on the grounds that eduPGS differences are inconsistent with known phenotypic score differences (e.g., Martin et al., 2017). However, that eduPGS constructed using the smaller within-family sample shows a tenuous validity highlights Duncan et al.’s (2019) caution that different eduPGS can give markedly different results.

<https://www.worldbank.org/en/publication/human-capital>
